# Diverse regulatory pathways modulate “bet hedging” of competence induction in epigenetically-differentiated phase variants of *Streptococcus pneumoniae*

**DOI:** 10.1101/2022.03.07.483185

**Authors:** Min Jung Kwun, Alexandru V. Ion, Marco R. Oggioni, Stephen D. Bentley, Nicholas J. Croucher

## Abstract

Despite enabling *Streptococcus pneumoniae* to acquire antibiotic resistance and evade vaccine-induced immunity, transformation occurs at variable rates across pneumococci. Phase variants of isolate RMV7, distinguished by altered methylation patterns driven by the translocating variable restriction-modification (*tvr*) locus, differed significantly in their transformation efficiencies and biofilm thicknesses. These differences were replicated when the corresponding *tvr* alleles were introduced into an RMV7 derivative lacking the locus. RNA-seq identified differential expression of the type 1 pilus, causing the variation in biofilm formation, and inhibition of competence induction in the less transformable variant, RMV7_domi_. This was partly attributable to lower expression of ManLMN in RMV7_domi_, which promoted competence induction through importing *N*-acetylglucosamine. This effect was potentiated by orthologues of the gram-negative competence regulatory machinery. Furthermore, a phage-related chromosomal island was more active in RMV7_domi_, which inhibited transformation by increasing expression of the stress response proteins ClpP and HrcA. However, HrcA increased competence induction in the other variant, with its effects depending on Ca^2+^ supplementation or heat shock. Hence the heterogeneity in transformation efficiency likely reflects the diverse signalling pathways by which it is affected. This regulatory complexity will modulate population-wide responses to synchronising quorum sensing signals to produce co-ordinated yet stochastic “bet hedging” behaviour.

---

Competence for natural transformation was first identified in *Streptococcus pneumoniae* (the pneumococcus) in the early 20^th^ century (1). Cells can be “transformed” to express a new phenotype through the acquisition of exogenous DNA, which is integrated into their genome through homologous recombination following its import through the specialised cell-encoded competence machinery (2). Transformation has played a key role in the emergence of antibiotic-resistant *S. pneumoniae*, both through generating “mosaic” alleles of core loci (3–5) and the acquisition of specialised resistance genes (6). It has also enabled vaccine evasion through recombinations affecting the capsule polysaccharide synthesis (*cps*) locus altering surface antigens (7, 8).

Despite the ability of transformation to accelerate such adaptive evolution in *S. pneumoniae*, considerable variation in the rate of diversification of strains through this mechanism persists across the species (9). Epidemiological studies have found the *r*/*m* ratio of base substitutions introduced through homologous recombination, relative to those occurring through point mutation, varies from well over 10 (7, 9) to below 0.1 (9, 10) across the species. Similarly, *in vitro* assays have identified >100-fold differences in the transformation efficiency of *S. pneumoniae* genotypes, with substantial variation even between isolates of the same serotype or strain (11–14). Many isolates are routinely found not to be transformable under standard conditions (11). This is often the consequence of integrative mobile genetic elements (MGEs) disrupting genes necessary for transformation (6, 15–17), selfishly preventing themselves from being eliminated from the chromosome (17). Yet in other non-transformable isolates, the highly-conserved competence machinery is intact (11, 18). This suggests the variation in transformation rates also reflects differences in regulation of the competence system.

The best-characterised signal inducing transformation in *S. pneumoniae* is the competence stimulating peptide (CSP) pheromone, which acts as a quorum-sensing system that signals through the ComDE two-component regulator (19). This activates early competence genes after about ten minutes (20). These include *comX*, encoding an alternative sigma factor (21). ComX enables the RNA polymerase to recognise late competence genes (20), which feature a “combox” signal in their promoters (22, 23). This results in pneumococci entering a transient competent state around 20 minutes post-CSP induction, after which the relevant machinery is degraded (24), and the cells become temporarily refractory to induction (21).

Transformation efficiency is known to vary between isogenic pneumococci through phase variation in capsule production. Transparent colony variants produce less capsule, and are less virulent, and more transformable, than opaque colony variants (25). This short-term variation has been linked to rapid changes at the phase-variable inverting variable restriction (*ivr*) locus, encoding the conserved Type I *Spn*III restriction-modification system (RMS) and the IvrR recombinase that drives sequence inversions within the locus (26–31). These rearrangements switch the target recognition domains (TRDs) within the active HsdS specificity protein, which determines the DNA motif that is targeted by both the methylase and endonuclease activities of the system. Consequently, changes at this single locus have pleiotropic effects through altering genome-wide methylation patterns (27). These phase-variable RMSs can thereby maintain phenotypic heterogeneity within a genetically near-homogenous population (32), resulting in “bet hedging” that can increase the chances of a species surviving a changing environment (33, 34).

The second pneumococcal phase-variable Type I restriction-modification system (named *Spn*IV) (27), encoded by the translocating variable restriction (*tvr*) locus (28), varies through excision-reintegration mediated by the recombinase TvrR (35). This locus is active in almost all pneumococci, but the complement of TRDs varies between isolates, increasing the diversity of HsdS proteins, and possible methylation patterns, across the species (28, 35). This locus is inactive in the R6 laboratory isolate that is typically used to study pneumococcal competence (28, 35, 36). Here, we characterised clinical isolates in which the *tvr* locus is intact, to understand how phase variation in *Spn*IV activity might contribute to phenotypic heterogeneity in clonally-derived populations.

## Results

### Phase variants of *S. pneumoniae* RMV7 differ in their transformation efficiency

A previous screen for variation at the *tvr* locus identified a diverse panel of restriction-modification variants (RMVs) (35). Four RMV isolates underwent sufficiently rapid phase variation in culture to enable the isolation of pairs of genotypes that differed in the motif targeted by their *Spn*IV systems, which is determined by the *hsdS* gene nearest the 5’ end of the locus (35), but were isogenic across the rest of the genome. The differences in the arrangement of the *tvr* loci were “locked” in each variant through *tvrR* being knocked out, or disrupted, by a selectable and counter-selectable Janus cassette marker (35, 37). These pairs of otherwise-isogenic locked phase variants were screened for differences in their transformation efficiency (Fig. 1A). In the RMV6 and RMV7 pairs, the variant in which the active *tvr* locus HsdS comprised the TRDs III-i (directing the *Spn*IV system to target the bipartite motif TGAN_7_TATC) was found to have significantly higher transformation efficiency, following induction by exogenous CSP, than their counterpart. These genotypes both originate from the multidrug-resistant GPSC1 strain (38). The RMV6 variants were found to be distinguished by a mutation in *dltD*, which commonly occurs during *in vitro* culturing (39) and affects cell wall biology. By contrast, the serotype 19F RMV7 variants exhibited a ~100-fold difference in their transformation rate (Fig. 1A), despite having identical *dlt* operons, and were therefore characterised in greater detail.

**Figure 1.**
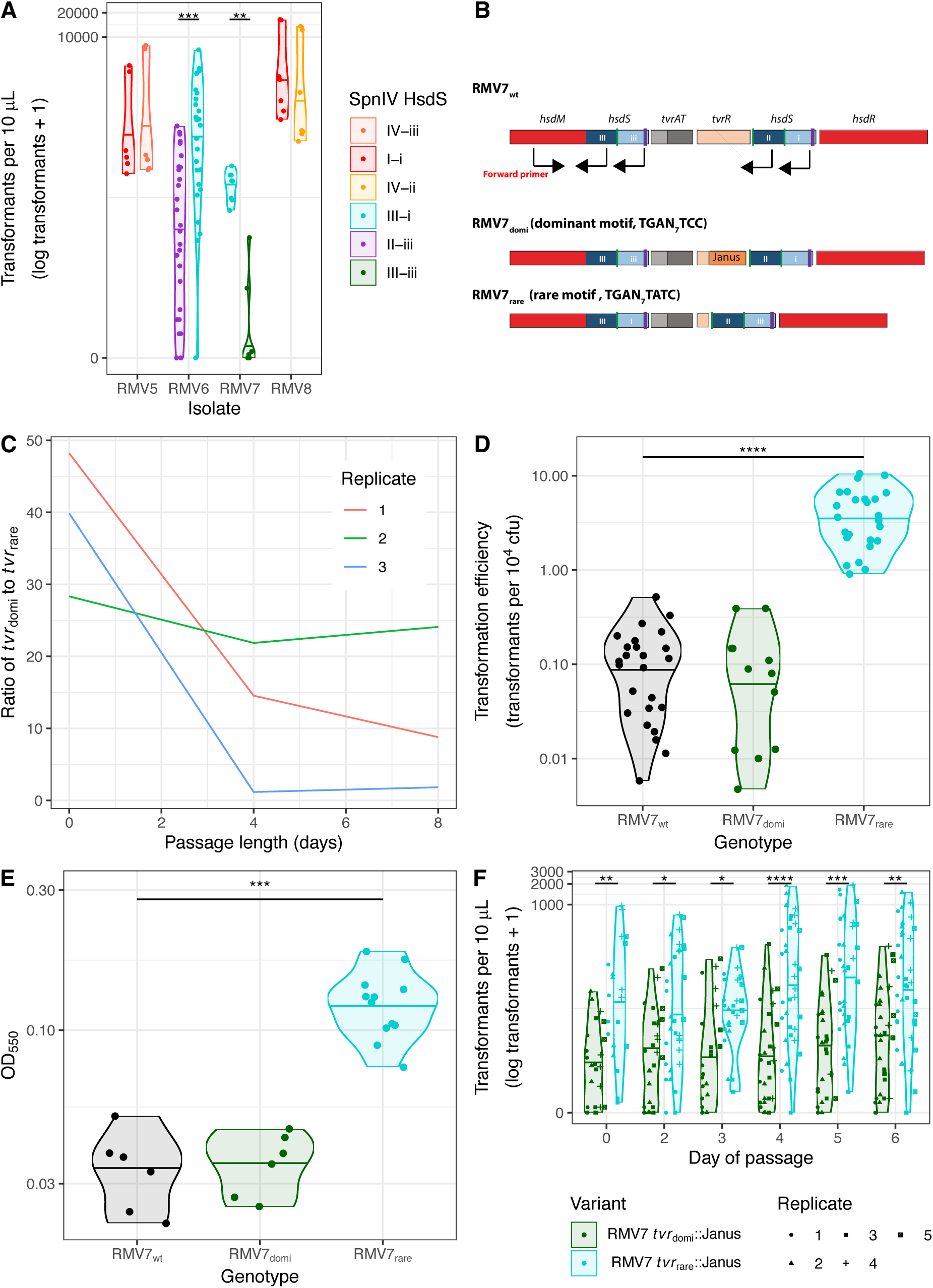
Phenotypic differences between locked *tvr* variants. (A) Violin plot showing the transformation efficiency of four pairs of *tvr* locus variants constructed from isolates RMV5, RMV6, RMV7 and RMV8. Each individual point represents an independent transformation experiment. The horizontal line within each violin shows the median for each genotype. The brackets indicate the statistical significance of the comparison between variants from the same isolate background, as calculated using a two-tailed Wilcoxon rank sum test. (B) Schematic of the *tvr* loci from RMV7_wt_, RMV7_domi_ and RMV7_rare_ to show the genes encoding the methylase (*hsdM*), endonuclease (*hsdR*), regulatory system (*tvrAT*) and recombinase (*tvrR*). The variants differ in their active *hsdS* genes, upstream of *tvrATR*. The RMV7_domi_ HsdS protein comprises the TRD combination III-iii (recognising motif TGAN_7_TCC), whereas that of RMV7_rare_ contains the TRDs III-I (recognising motif TGAN_7_TATC). The black arrows indicate the binding sites of a forward primer, in *hsdM*, and reverse primers, in *hsdS* fragments. (C) Line graph showing the ratio of RMV7_domi_ to RMV7_rare_ loci in eight-day passages of RMV7_wt_. The RMV7_domi_ variant was more common in each of the three replicates at every measured timepoint. (D) Violin plot showing the higher transformation efficiency of RMV_rare_ relative to RMV7_domi_ or RMV7_wt_. Each individual point represents an independent experiment in which the number of transformants, and number of overall colony-forming units (cfu), was calculated. The horizontal line within each violin shows the median for each genotype. Both mutants were compared with RMV7_wt_; the horizontal bracket shows a significant difference, as calculated from a two-tailed Wilcoxon rank sum test. (E) Violin plot showing the adhesion of variants to an abiotic surface, as quantified by the optical density at 550 nm after crystal violet staining of cells resisting washing from a microplate well. The RMV7_rare_ variant showed a significantly greater adhesion after 16 hours incubation at 35 °C, as assessed by a two-tailed Wilcoxon rank sum test. (F) Violin plots showing the transformation efficiency of knock-in mutants during a passage experiment. The *tvr* loci of RMV7 *tvr*_domi_::Janus and RMV7 *tvr*_rare_::Janus (both containing a *tvrR* gene interrupted by a Janus cassette) were each introduced into an RMV7 *tvr*::*cat* background (Fig. S5). This pair were separately passaged in liquid cultures over six days in five independent replicates. The number of transformants observed from three transformation assays, conducted each day for both variants, is shown by the individual points’ shapes and colours. The violin plots summarise the median and distribution of these values. The brackets indicate the statistical significance of the comparison between variants from the same day of the passage, as calculated using a two-tailed Wilcoxon rank sum test. Across all panels, significance is coded as: *p* < 0.05, *; *p* < 0.01, **; *p* < 10^−3^, ***; *p* < 10^−4^, ****. All *p* values were subject to a Holm-Bonferroni correction within each panel.

This substantial difference reflected the RMV7 variant carrying the alternative form of the *tvr* locus, with an active HsdS comprising the TRDs III-iii (which directs the *Spn*IV system to target the motif TGAN_7_TCC; Fig. 1B and S1), having an almost-undetectable transformation efficiency. Culturing the wild-type isolate (RMV7_wt_) over successive days demonstrated large changes in the relative frequency of these *tvr* variants, although the less transformable variant (RMV7 *tvrR*::Janus; henceforth, RMV7_domi_, carrying *tvr*_domi_) was typically dominant in prevalence relative to the rarer, more transformable variant (RMV7 Δ*tvrR*; henceforth, RMV7_rare_, carrying *tvr*_rare_; Fig. 1C). The unmodified RMV7_wt_ had a transformation efficiency similar to RMV7_domi_, concordant with the relative proportions of the variants observed *in vitro*, whereas that of RMV7_rare_ was confirmed to be ~50-fold higher (Fig. 1D). This difference could not be explained by high spontaneous mutation rates, nor by changes in the speed at which competence for transformation was activated (Fig. S2). Therefore, RMV7_domi_ and RMV7_rare_ exhibited distinctive phenotypes that correlated with their *tvr* locus arrangements.

RMV7_rare_ was also significantly more adhesive to an abiotic surface (Fig. 1E), which can be considered a proxy for biofilm formation (40). These differences in both adhesion and transformation could be explained by RMV7_rare_ being enriched for transparent phase variants. However, microscopy found no clear difference in colony morphology between the variants (Fig. S3). An alternative explanation for the phenotypic differences could be mutations that occurred during genetic manipulation of the isolates (41). Alignment of the two variants’ assemblies found them to be distinguished by seven non-synonymous single nucleotide polymorphisms and two premature stop codons outside the *tvr* locus, none of which were within genes known to directly affect the competence machinery (Table S1). Nevertheless, we tested how transformation was affected by mutations in RMV7_rare_ that were absent from both the RMV7_domi_ and RMV7_wt_ sequences. Neither a non-synonymous change in *phoB*, encoding a phosphate-sensitive response regulator (42), nor disruption of the transporter gene *pstS*, affect by a premature stop codon in RMV7_rare_, explained the contrasting transformation efficiencies of the variants (Fig. S4). Hence the differences between RMV7 variants could not be explained by point mutations or alterations in encapsulation.

To test whether the phenotypic differences were causatively associated with variation in the *Spn*IV RMS, the *tvr* locus of RMV7_wt_ was replaced with a chloramphenicol acetyltransferase (*cat*) resistance marker to generate RMV7_wt_ *tvr*::*cat*. The *tvr* loci of RMV7_domi_ and RMV7_rare_ (both modified by a Janus cassette within *tvrR*; see Methods) were separately introduced into RMV7_wt_ *tvr*::*cat*, generating the otherwise isogenic knock-in recombinants RMV7 *tvr*_domi_::Janus and RMV7 *tvr*_rare_::Janus, carrying the two different locked *tvr* loci (Fig. S5). An initial characterisation of these genotypes demonstrated RMV7_wt_ *tvr*::*cat* was substantially more transformable than RMV7_wt_, suggesting methylation at *tvr*_domi_ motifs caused the repression of competence induction (Fig. S5). The *tvr*_domi_::Janus and *tvr*_rare_::Janus mutants reproduced the phenotypic divergence between RMV7_domi_ and RMV7_rare_ in both biofilm formation and transformation (Fig. S5), providing further evidence that these differences were driven by epigenetic variation.

The ~17-fold difference in transformation efficiency between the *tvr*_domi_::Janus and *tvr*_rare_::Janus variants was nevertheless smaller than that measured between RMV7_domi_ and RMV7_rare_. Hence the transformability of these genotypes was assayed over five independent six-day passages, to test whether the difference between them would rise, in case any consequences of DNA methylation were slow to emerge. However, the observed disparity in the transformation efficiency of the *tvr*_domi_::Janus and *tvr*_rare_::Janus genotypes was stable (Fig. 1F). This smaller difference may represent changes in the expression of the introduced *tvr* loci, the effect of mutations outside the *tvr* locus in RMV7_domi_ or RMV7_rare_, else suggest that the effect of methylation on gene expression may be indirectly mediated via effects on nucleoid organisation. Nevertheless, both naturally-isolated and genetically-engineered pairs of RMV7 *tvr*_rare_ and *tvr*_domi_ variants replicated a significant and reproducible difference in biofilm formation and transformation efficiency.

### Epigenetic effects on the induction of competence genes

To understand how the *tvr* loci caused a difference in transformation, RNA-seq was used to quantify patterns of transcription in the recombinants RMV7 *tvr*_domi_::Janus and RMV7 *tvr*_rare_::Janus. Samples were taken pre-CSP, 10 minutes post-CSP, and 20 minutes post-CSP from each of three biological replicates (Fig. 2). The 18 RNA samples were sequenced as 200 nt paired-end multiplexed libraries on a single Illumina HiSeq 4000 lane. After alignment to the RMV7_domi_ genome, analysis of the RNA-seq data found the fragment length distributions (Fig. S6) and gene expression densities (Fig. S7) were consistent across samples (Table S2). Q-Q plots suggested a Benjamini-Hochberg corrected *q* value of 10^−3^ was an appropriate threshold for identifying significant transcriptional variation (Fig. S8-9). This identified 154 genes that significantly differed in their expression between the two genotypes prior to CSP exposure, or between the pre- and post-CSP samples from the same genotype (Fig. 2; Table S3).

**Figure 2.**
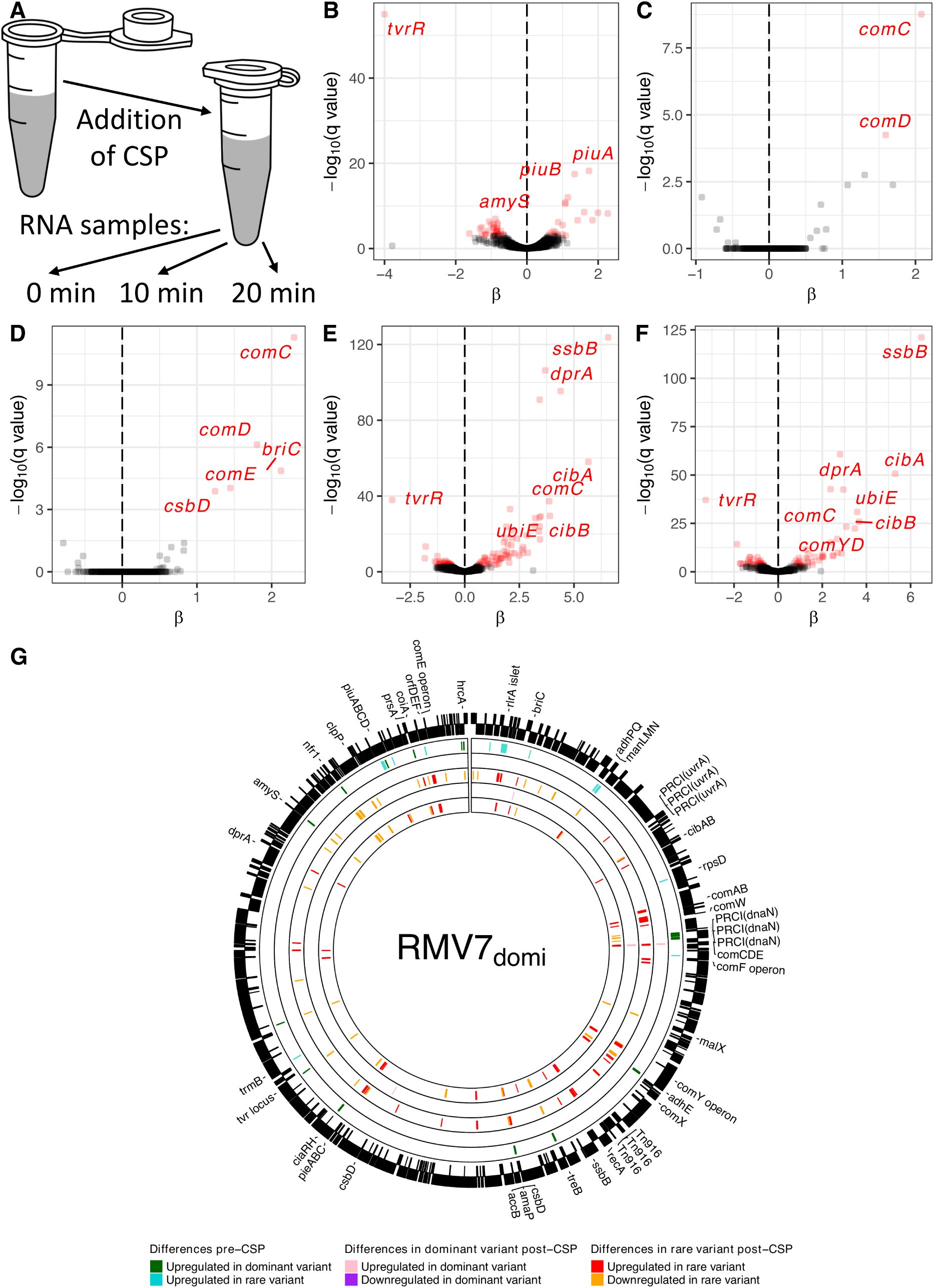
Genes exhibiting significant differences in transcription between RNA-seq samples. (A) Design of the RNA-seq experiment. (B)-(F) Volcano plots showing the variation in gene transcription between RNA-seq samples. The horizontal axis shows the natural logarithm of the fold difference in expression levels between the genotypes, β. The vertical axis shows the negative base 10 logarithm of the *q* value, corresponding to a Benjamini-Hochberg corrected *p* value. Points are coloured red where this value exceeds the threshold false discovery rate threshold of 10^−3^. The panels correspond to (B) the differences between the pre-CSP samples of *tvr*_domi_::Janus and *tvr*_rare_::Janus; the differences between the pre-CSP and 10 minute post-CSP samples for (C) *tvr*_domi_::Janus and (D) *tvr*_rare_::Janus; and the differences between the pre-CSP and 20 minute post-CSP samples for (E) *tvr*_domi_::Janus and (F) *tvr*_rare_::Janus. The significant difference in *tvrR* expression between the *tvr*_domi_::Janus and *tvr*_rare_::Janus mutants is an artefact of the different constructs used to disrupt *tvrR* in these two genotypes (see Methods). (G) Chromosomal distribution of differentially-expressed genes. The outer ring shows the annotation of RMV7_domi_ (accession code OV904788). Protein coding sequences are represented as black boxes, with the vertical positioning within the ring indicating the strand of the genome on which they are encoded. The next ring inwards shows significant pre-CSP differences in transcription: green genes were more highly expressed in RMV_domi_, and blue genes were more highly expressed in RMV_rare_. The next ring inwards show significant changes in gene expression 10 minutes post-CSP in RMV7_domi_: pink genes were upregulated, and purple genes were downregulated. The third ring inwards shows significant changes in gene expression 10 minutes post-CSP in RMV7_rare_: red genes were upregulated, and orange genes were downregulated. The two inner rings repeat this representation for changes in gene expression 20 minutes post-CSP.

Comparing the overall transcriptional patterns among datasets found the biggest separation distinguished the post-CSP RMV7 *tvr*_rare_::Janus transcriptomes from the others (Fig. S10), suggesting a major difference in the induction of the competence system between the variants. In RMV7 *tvr*_rare_::Janus, the early competence genes showed elevated expression 10 minutes post-CSP, including a >20-fold induction of *comCDE* (Fig. S11). The late competence genes exhibited more variable patterns of transcription (Fig. S12). Multiple nucleotide metabolism and transporter genes (*purA*, *tadA*, *dut*, *ribF*, *adeQ*; Fig. S13) were upregulated, and transcription of the competence-induced biofilm formation signal gene *briC* (43) rose >10-fold (Fig. S11). Transcription of a three gene cluster encoding another transporter, induced by CSP (44) and antimicrobial peptides (45), rose >3 fold, and consequently was named *pieABC* (peptide-induced exporter; CDSs IONPJBJN_01324-6 in RMV7_domi_, corresponding to SP_0785-787 in TIGR4; Fig. S13, Table S3).

By contrast, CSP significantly upregulated just five genes in RMV7 *tvr*_domi_::Janus: 5.7-fold and 6.5-fold increases in the transcription of the quorum sensing genes *comCDE* and *briC* respectively (Fig. S11), and a 2.9-fold increase in transcription of the stress response gene *csbD*. However, there was no sign of late competence genes being activated, which requires the alternative sigma factor ComX. However, expression of the *comX* gene itself can be difficult to determine through RNA-seq (Fig. S11), owing to the presence of two near-identical paralogues in pneumococcal genomes (46).

As an independent test of these transcriptional differences, qRT-PCR experiments were undertaken on the original RMV7_domi_ and RMV7_rare_ variants, and the control genotypes RMV7_wt_ and RMV7_wt_ *tvr*::*cat*. The qRT-PCR data showed genotypes of both lower (RMV7_wt_ and RMV7_domi_) and higher (RMV7_rare_ and RMV7_wt_ *tvr*::Janus) transformation efficiency up-regulated the early competence genes *comD* and *comX* in response to CSP (Fig. S14). However, the late competence genes *comEA* and *comYC* were only significantly upregulated in the more transformable genotypes. Hence the difference in transformability between the RMV7 variants was a consequence of late competence gene activation being blocked in RMV7_domi_ through effects of *tvr*_domi_ expression.

### Pre-CSP expression differences associate elevated intracellular stress with *tvr*_domi_

This difference in competence induction was likely caused by the 53 genes that significantly differed in their expression between RMV7 *tvr*_domi_::Janus and *tvr*_rare_::Janus prior to CSP exposure (Fig. 2; Table S3). These did not include any *cps* locus genes, which were non-significantly more highly expressed in RMV7 *tvr*_rare_::Janus (Fig. S15), confirming that the elevated transformation efficiency of this genotype did not reflect an enrichment of transparent phase variants (27, 47). An analysis of the distances from protein coding sequence start codons to the nearest upstream methylation site found no general relationship between differential expression and proximal methylation for either *tvr*_domi_ or *tvr*_rare_ motifs (Fig. S1 and S16). Of the four cases of significantly differentially-expressed genes being close to variable methylation sites, the modification was only likely to modulate transcription initiation at the *piuABCD* operon (Fig. S17). This encodes an iron transporter, and was >5-fold more highly expressed in RMV7_rare_ (Fig. 2B; Fig. S13). However, knocking out *piuA* did not reduce the transformation efficiency of RMV7_rare_, suggesting this change was independent of those affecting competence (Fig. S18).

Hence the differences in the pre-CSP transcriptomes are unlikely to be attributable to a small number of promoters that are strongly affected by direct modification, consistent with other genome-wide studies (27, 48). Instead, the differences likely represent the consequences of chromosome-level changes in DNA conformation or nucleoid interactions at particularly sensitive promoters (49). One notable aspect of the overall distribution of the *Spn*IV target motifs was that the *tvr*_rare_ motifs were uniformly distributed, whereas the *tvr*_domi_ motifs were enriched in one segment of the chromosome (Fig. S19). Such an uneven distribution could accentuate the effects of shifting patterns of modification.

Correspondingly, three transcriptional patterns suggested the *tvr*_domi_ methylation pattern was associated with dysregulation and increased intracellular stress. The first was the activation of heat shock responses. Transcription of the chaperone gene *prsA* and chaperone regulator gene *hrcA* were 3.4-fold and 4.1-fold higher in *tvr*_domi_::Janus, respectively. Correspondingly, the *groES*-*groEL* and *grpE*-*dnaK*-IONPJBJN_02152-*dnaJ* operons of the *hrcA* regulon were also more highly expressed in *tvr*_domi_::Janus, although the difference was only significant for IONPJBJN_02152 (Fig. S21; Table S3). The *accBC*-*yqhY*-*amaP*-*csbD* cell wall stress operon (50) (IONPJBJN_01032-6) was also upregulated in RMV7 *tvr*_domi_::Janus (Fig. S20). This was likely a consequence of increased expression of the *mgrA* gene, encoding a regulator, in RMV7 *tvr*_domi_::Janus, albeit this difference did not reach genome-wide significance (Fig. S20). Consistent with this activity of MgrA, the *rlrA* islet, encoding the type 1 pilus, was approximately five-fold lower in *tvr*_domi_::Janus (51).

The second indicator of stress in in RMV7 *tvr*_domi_::Janus was the 3.7-fold higher transcription of *ciaRH*, encoding a two-component system that enables cells to survive lysis-inducing conditions (52), and is known to inhibit competence (53). The third was the increased ~1.5-fold higher transcription of a phage-related chromosomal island (PRCI; also known as a phage-inducible chromosomal island, or PICI), integrated adjacent to *dnaN* (PRCI*_dnaN_*; Fig. S22; Table S3). This is one of two PRCIs associated with GPSC1 (7, 28), the other being integrated near *uvrA* (Fig. 2). The regulatory mechanisms of these elements are not thoroughly characterised (54), and in the absence of a helper prophage, it is unclear exactly what signal may have triggered this increased activity. Given integrative MGEs are likely under selection to reduce host cell transformability (17), the increased activity of PRCI*_dnaN_* could drive inhibition of the competence machinery. Hence PRCI*_dnaN_* and *ciaRH* were the primary candidates for causing the observed difference in transformability between the RMV7 variants.

### ManLMN links competence induction to carbon source metabolism

The higher expression of the *ciaRH* genes in the poorly-transformable genotypes RMV7_domi_ and RMV7_wt_, relative to the more transformable genotype RMV7_rare_, was confirmed by qRT-PCR (Fig S15). CiaRH binds at least 15 promoter sequences, five of which drive the expression of *cia*-dependent small RNAs (csRNA) that suppress the induction of competence by inhibiting CSP production (55). Their expression is difficult to determine using conventional RNA-seq approaches due to their size (56), but they are unlikely to affect competence induced by exogenous CSP (57). The remaining ten drive the expression of protein coding sequences (55) (CDSs), which were more highly expressed in RMV7 *tvr*_domi_::Janus compared to RMV7 *tvr*_rare_::Janus pre-CSP (Fig. S23). These included the extracytoplasmic chaperone and serine protease HtrA (55, 58, 59), which can block competence induction at low CSP concentrations through degrading extracellular CSP (58, 60). In agreement with some previous studies, elimination of *htrA* further increased the transformability of RMV7_rare_ (58), but the same mutation had no significant effect in RMV7_wt_ (Fig S24). This suggests HtrA inhibits the induction of competence, but is unlikely to explain much of the difference between these variants. Similarly, knock out of *ciaRH* itself had little effect in either genotype (Fig. S24), suggesting its expression was correlated with, rather than causative of, the difference in transformation efficiency between the variants.

Among the CiaRH regulon, the most significant difference in expression between RMV7_rare_ and RMV7_domi_ was the ManLMN carbon source importer operon (Table S3). The *manLMN* genes were more highly expressed in RMV7_rare_, as CiaRH binds promoter sequences within the operon (36, 55, 61, 62) and acts as a repressor (55, 63). Disrupting the *manLMN* locus reduced the transformation efficiency of RMV7_rare_ by >5-fold, while having little effect in RMV7_wt_ (Fig. 3A-B, S25). To test whether the observed transformation differences reflected a growth defect, *manLMN*::Janus mutants of both RMV7_wt_ and RMV7_rare_ were cultured in the same rich media (Fig. S26; Table S4). RMV7_domi_ grew more slowly than RMV7_rare_, consistent with the former suffering greater intracellular stress. However, removal of *manLMN* had little effect on growth in either variant, suggesting the transporter’s effect on transformation was through regulation rather than proliferation. Hence the lower expression of *manLMN* in RMV7_domi_ accounts for much of its reduced transformation efficiency relative to RMV7_rare_.

**Figure 3.**
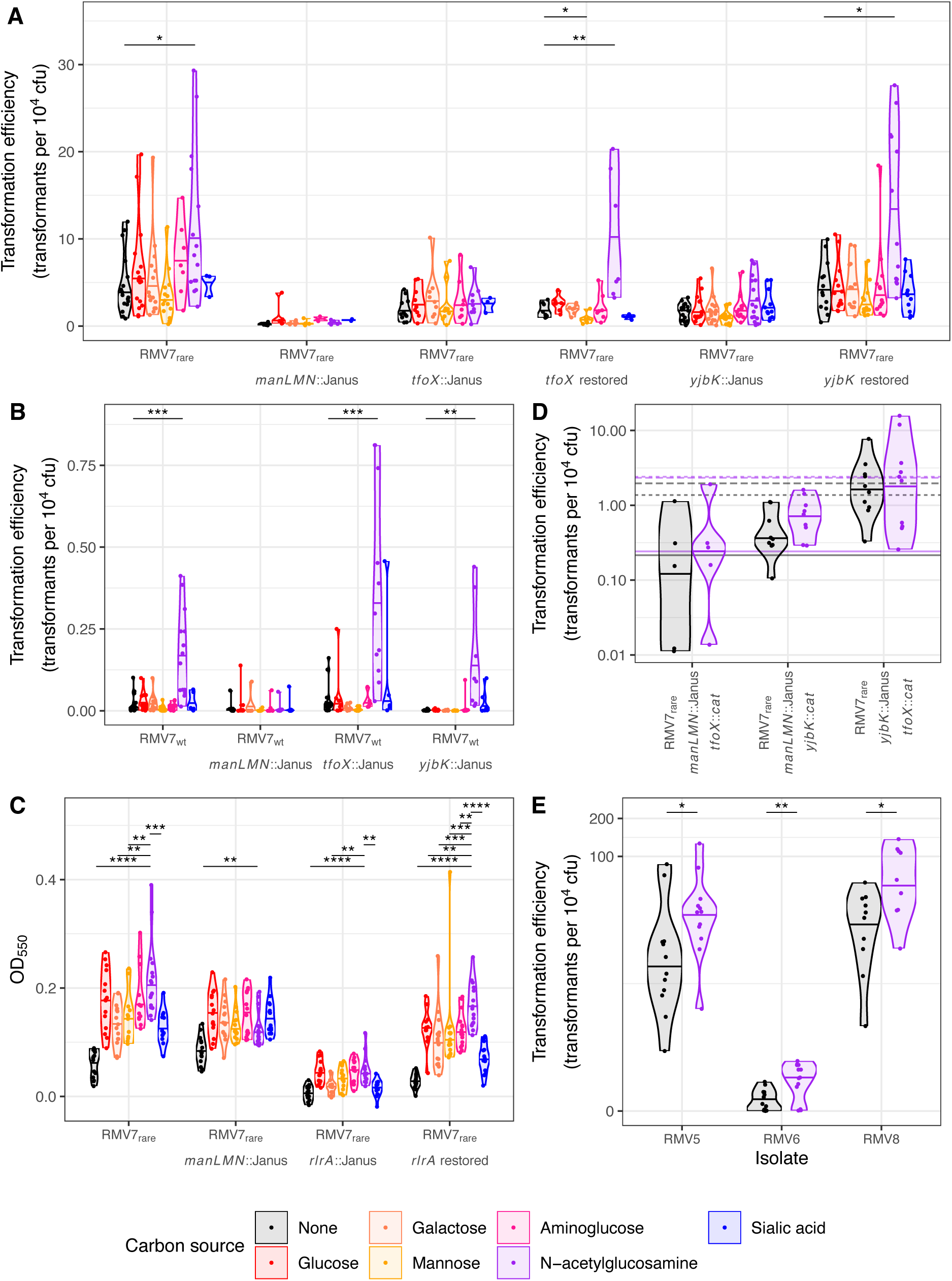
The dependence of transformation efficiency on import of carbon sources. (A) Violin plots showing the transformation efficiency of RMV7_rare_ relative to mutant derivatives. The counter selectable Janus cassette was used to disrupt the *manLMN*, *tfoX* and *yjbK* genes. This enabled the restoration of the *tfoX* and *yjbK* genes (similar data for *manL* are shown in Fig. S28). Each genotype was transformed in unsupplemented media, and in the presence of one of six carbon sources, as indicated by the plot colour (see key). Each point represents an independent experiment, and the horizontal line within the violin plots show the median for each combination of recipient cell genotype and carbon source. For each genotype, Wilcoxon rank-sum tests were used to test for evidence of changes in transformation efficiency caused by each carbon source, relative to the unsupplemented media. Significant differences are indicated by the black brackets at the top of the panel. (B) Violin plots showing the transformation efficiency of RMV7_wt_ relative to mutant derivatives. Data are displayed as in panel A. (C) Violin plots quantifying the effects of carbon source supplementation, and disruption of *manLMN* and the *rlrA* pilus islet, on adhesion of bacteria to an abiotic surface using OD_550_. The density of the biofilm formed following growth in GlcNAc-supplemented media was used as the comparator for Wilcoxon rank sum tests, as all carbon source supplements increased biofilm formation relative to unsupplemented media. (D) Violin plots showing the transformation efficiency of double mutants constructed in the RMV7_rare_ genotype. For comparison with panel A, the horizontal lines show the median transformation efficiencies of the corresponding single mutants in unsupplemented and GlcNAc-supplemented media: *manLMN*::Janus as solid lines; *tfoX*::Janus as dotted lines, and *yjbK*::Janus as a dashed line. Data are shown on a logarithmic scale to resolve the differences between these poorly-transformable genotypes. (E) Violin plots showing the effect of GlcNAc supplementation on the transformation efficiency of the most common *tvr* variants of the RMV5, RMV6 and RMV8 isolates. Across all panels, significance was coded as: *p* < 0.05, *; *p* < 0.01, **; *p* < 10^−3^, ***; *p* < 10^−4^, ****. All *p* values were subject to a Holm-Bonferroni correction within each panel.

ManLMN is a phosphotransferase system (PTS) transporter that can facilitate the import of glucose, mannose, galactose, fructose, aminoglucose and *N*-acetylglucosamine (GlcNAc) (64). Supplementation of liquid media with these carbon sources did not substantially affect growth profiles (Fig. S26), apart from the addition of glucose causing small ManLMN-dependent increases in the replication rate of both phase variants (Fig. S27). However, GlcNAc supplementation increased transformation efficiency ~10-fold in RMV7_wt_ (Fig. 3A) and ~2-fold in RMV7_rare_ (Fig. 3B). In both variants, this effect was dependent upon *manLMN*, as was confirmed through disrupting, and then restoring, *manL* (Fig. S28). This potentiation of the induction of competence by GlcNAc was also observed in the other RMV isolates (Fig. 3E). Therefore, ManLMN links carbon source availability to pneumococcal transformability.

### *N*-acetylglucosamine activates competence through TfoX and YjbK

Competence in *Vibrio cholerae* is induced GlcNAc disaccharides (65), thought to be generated from degradation of chitin (66). This is mediated through the Transformation Factor X (TfoX) protein (67, 68). An orthologue of this protein (TfoX*_Hflu_*, also called Sxy) is also central to induction of competence in *Haemophilus influenzae* (69) by 3’,5’-cyclic AMP (cAMP) signalling (70, 71). Intracellular concentrations of 3’,5’-cAMP rise in many Proteobacteria when the primary glucose PTS transporter is inactive, as the accumulation of phosphorylated EIIA PTS subunits stimulates adenylate cyclase activity (72). Search were undertaken for analogues of either of these pathways in *S. pneumoniae*.

An orthologue of the N terminal domain of TfoX*_Hflu_* was annotated, but not described, in *S. pneumoniae* ATCC 700669 (73). In RMV7, this gene (*tfoX*_Spn_; IONPJBJN_02097) is conserved in the same genomic location as in ATCC 706669, two genes upstream of the *comEA* competence operon (Fig. S29). The TfoX*_Spn_* amino acid sequence was predicted to form a four-strand beta sheet flanked by alpha helices (Fig. S30), resembling the N-terminal domain of gram-negative orthologues (Fig. S31). Disruption of *tfoX*_Spn_ in RMV7_rare_ both decreased the pneumococcus’ basal transformation efficiency in the absence of supplements (Fig. S25), and eliminated the GlcNAc-induced elevation in this rate (Fig. 3C). Restoring *tfoX* rescued RMV7_rare_’s responsiveness to GlcNAc.

A gene encoding a candidate adenylate cyclase, *yjbK*, was also identified in RMV7 (IONPJBJN_01639). This protein is predicted to have a β-barrel structure (Fig. S32), as observed for orthologous enzymes synthesising 3’,5’-cAMP (Fig. S33). The *yjbK* gene could be both disrupted, and restored, in RMV7_rare_. Transformation assays with these genotypes demonstrated the loss of YjbK reduced the transformation efficiency of RMV7_rare_ both in the absence of supplements, and following the addition of GlcNAc (Fig. 3D, S25). However, no 3’,5’-cAMP signalling pathway is known in Firmicutes (74). Correspondingly, an ELISA assay demonstrated 3’,5’-cAMP levels in *S. pneumoniae* were close to the lower detection threshold, far below those of *Escherichia coli*, and unaffected by *yjbK* disruption (Fig. S34). Additionally, exogenous 3’,5’-cAMP had no effect on transformation efficiencies in any RMV7 genotypes (Fig. S35). Therefore, it is unlikely that YjbK’s regulatory role is mediated through 3’,5’-cAMP production.

To test whether the effects of GlcNAc were the consequence of it acting as a signal, or as a metabolic substrate, *nagA* was disrupted in RMV7_wt_ and RMV7_rare_ (Fig. S36, S37). The import of GlcNAc by ManLMN generates intracellular GlcNAc-6-phosphate, which can be used in cell wall synthesis, or converted to glucosamine-6-phosphate by NagA. Glucosamine-6-phosphate is also generated by the import of aminoglucose by ManLMN, and must be converted to the glycolytic substrate fructose-6-phosphate, as both GlcNAc-6-phosphate and glucosamine-6-phosphate are cytotoxic (75). The *nagA*::Janus mutant in the faster-growing RMV7_rare_ variant exhibited a growth defect that was exacerbated by GlcNAc supplementation, demonstrating NagA was necessary for processing imported GlcNAc. Similarly, a *nagA*::Janus mutant in the more GlcNAc-responsive RMV7_wt_ variant could not be transformed in the presence of GlcNAc (Fig. S37). However, exogenous aminoglucose did not increase growth or transformation in these *nagA*^−^ mutants, nor in any *nagA*^+^ genotypes (Fig. 3), confirming the effect of GlcNAc on transformation was not caused by faster growth or glycolysis. By contrast, RMV7_rare_ *tfoX*::Janus and *yjbK*::Janus mutants grew faster than RMV7_rare_ (Fig. S36), and this difference was enhanced when media were supplemented with GlcNAc (Fig. S37), demonstrating their reduced transformation efficiencies did not reflect a growth defect. This heightened replication rate occurred despite the disruption of *tfoX* and *yjbK* not affecting *nagA* transcription, and decreasing the expression of *manL* slightly. Therefore TfoX and YjbK appear to be regulators, rather than metabolic enzymes. Hence the differential effects of GlcNAc and aminoglucose on cells is likely to reflect signalling effects specific to GlcNAc involving TfoX, YjbK, and other proteins.

### GlcNAc and ManLMN regulate pneumococcal physiology through multiple pathways

Disruption of *manLMN*, *tfoX* and *yjbK* in the laboratory genotype R6 was also found to reduce transformation efficiency, although detecting these effects required culturing in a low-sugar chemically-defined medium (see Methods; Fig. S38). However, disrupting *yjbK* and *tfoX* in RMV7_wt_ did not affect the GlcNAc-associated increase in transformation efficiency, although transformation was notably reduced in the *yjbK*::Janus mutant (Fig. 3E). These results suggested ManLMN affected competence through at least two pathways, at least one of which was dependent upon TfoX and YjbK.

To test this proposed arrangement of pathways, genotypes were constructed by combining pairs of mutations in *manLMN*, *tfoX* and *yjbK*. Transformation assays demonstrated the double mutants in which *manLMN* was disrupted behaved similarly to the *manLMN*::Janus single mutant (Fig. 3D). Disruption of both *tfoX* and *yjbK* did not cause a substantially greater effect than observed in either of the corresponding single mutants. This suggested both operated intracellularly within the same pathway (Fig. 3D), and their activity depended on the import of molecules by ManLMN. This suggests ManLMN affects transformation efficiency through multiple pathways, one of which involves both TfoX and YjbK, consistent with the *tfoX*::Janus and *yjbK*::Janus mutants exhibiting similar phenotypes (Fig. 3 and S36, S37). Hence ManLMN is a key pleiotropic regulator.

To test whether differences in ManLMN activity may also explain the difference in biofilm formation between the RMV7 variants, adhesion of RMV7_rare_ and derived *manLMN*::Janus mutants to an abiotic surface was quantified in the presence of different carbon sources (Fig. 3F). Although disruption of *manLMN* resulted in changes to biofilm thicknesses in response to different carbon source supplements, there was little difference in unsupplemented media. Instead, we hypothesised that the difference was caused by the type 1 pilus, as the *rlrA* pilus islet was more highly expressed in RMV7 *tvr*_rare_::Janus, which replicated the thicker biofilm phenotype of RMV7_rare_ (Fig. 2). Disruption of the pilus structural genes (*rrgABC*), or their activator gene (*rlrA*), reduced the surface adherence of RMV7_rare_ to that of RMV7_wt._

However, the same mutation in the RMV7_wt_ background had little effect (Fig. S39). Restoring the pilus in RMV7_rare_ rescued the biofilm thickness phenotype (Fig. 3C). Hence multiple regulatory pathways underlie the individual phenotypic differences between the phase variants.

### Mobile element activation represses transformation by increasing intracellular stress

As additional regulatory pathways were likely to affect competence induction, we tested the hypothesis that the increased activity of PRCI*_dnaN_* in RMV7 *tvr*_domi_::Janus may also inhibit the competence of the host cell, in order to prevent the MGE being deleted through homologous recombination (17). The entire element was removed, either with (RMV7_wt_ PRCI*_dnaN+att_*::Janus) or without (RMV7_wt_ PRCI*_dnaN_*::Janus) the flanking *att* sites. In both cases, a ~5-fold increase in transformation rates was observed (Fig. 4A). This implied the PRCI inhibited the activation of the competence system. To if this was caused by a specific locus within the MGE, four mutations were generated within the PRCI: one removing the regulatory genes; one removing the regulatory genes and IONPJBJN_00496, a gene that encoded a protein similar to DNA damage-inducible protein D (DinD), which inhibits RecA activity in *E. coli* (76); one removing the replication genes; and one removing the integration, regulatory and replication genes. However, none of these mutations had such a large effect on transformation rates as the elimination of the entire element (Fig. 4A-B). This suggested the inhibition of competence induction was not the consequence of a single gene product, but instead the activity of the MGE itself.

**Figure 4.**
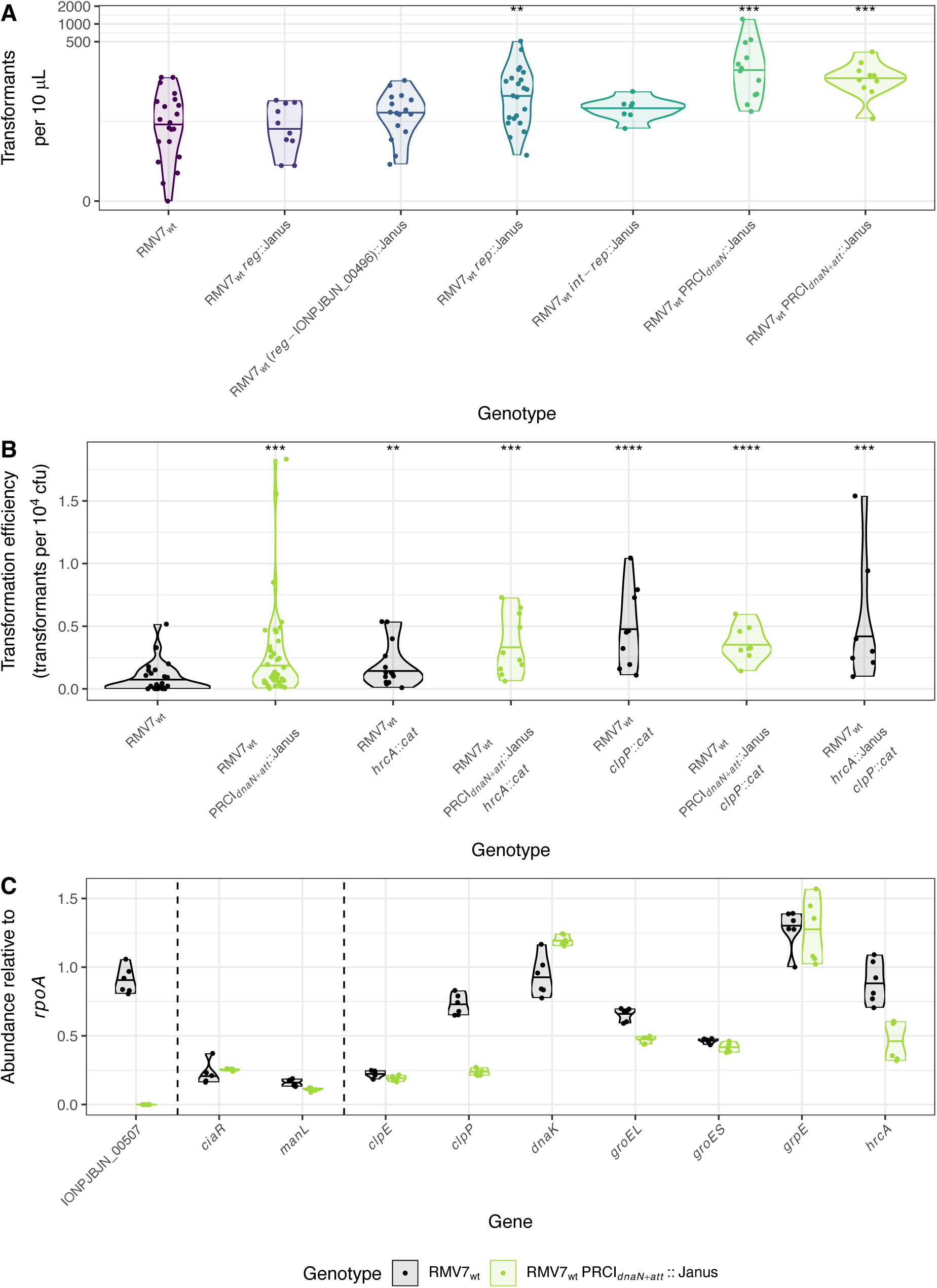
Effect of removing PRCI*_dnaN_* on the transformation efficiency of RMV7_wt_. (A) Violin plots showing the number of transformants observed in assays of RMV7_wt_ mutants in which different parts of PRCI*_dnaN_* were replaced with a Janus cassette. The genotypes are arranged left to right, and coloured black to purple, to represent the increasing proportion of the element that was replaced by the cassette. Mutants removed different combinations of the *dinD*-like gene (IONPJBJN_00496); the regulatory genes (*rep*); the replication genes (*rep*); the integration genes (*int*), and the *att* site. The structure of RMV7 PRCI*_dnaN_* meant that replacing the *int*-*rep* region also deleted the intervening *reg* genes. Each point represents an independent transformation assay. The violin plot summarises the result for each mutant, with a horizontal line indicating the median. Each mutant was compared against the parental RMV7_wt_ genotype using a Wilcoxon rank sum test. Black brackets at the top of the plot indicate significant differences in the number of observed transformants. (B) Violin plot quantifying the effect of PRCI*_dnaN_* and the chaperones HrcA and ClpP on transformation efficiency in RMV7_wt_. The comparison of RMV7_wt_ with a mutant in which the PRCI and its *att* site were removed was independent of the experiments presented in panel A, and more accurately quantified transformation efficiency. This approach was also used to compare the effects of single and double mutations that disrupted *hrcA*, *clpP* and PRCI*_dnaN_*. (C) Violin plots showing the effect of the disruption of PRCI*_dnaN_* on gene expression, quantified as abundance relative to *rpoA* by qRT-PCR. IONPJBJN_00507 is a coding sequence within the PRCI that is absent from RMV7_wt_ PRCI*_dnaN_*_+*att*_::Janus. The six points for each gene correspond to three technical replicate assays on each of two biological replicates. The horizontal line on the violin plot shows the median relative abundance for each gene in each genotype. Across all panels, significance is coded as: *p* < 0.05, *; *p* < 0.01, **; *p* < 10^−3^, ***; *p* < 10^−4^, ****. All *p* values were subject to a Holm-Bonferroni correction within each panel.

A qRT-PCR assay was employed to test whether the deletion of PRCI*_dnaN_* could affect transformation through disrupting the expression of other loci. Neither *ciaR* nor *manL* expression was altered when the PRCI was removed (Fig. 4C), suggesting an alternative pathway was involved. As PRCI*_dnaN_* caused a growth defect in RMV7_domi_ (Fig. S40), it was hypothesised that the element’s activity might drive the higher expression of the stress response proteins in *tvr*_domi_::Janus (Fig. S21). Correspondingly, transcription of both the chaperone regulator HrcA and chaperone-protease ClpP was approximately halved following deletion of PRCI*_dnaN_* in RMV7_wt_ (Fig. 4C). These genes were confirmed to more highly expressed in RMV7_domi_ than RMV7_rare_ by qRT-PCR (Fig. S21, S41). Yet the transcription of multiple other chaperones that were significantly more highly expressed in RMV7_domi_ (Table S3) were unperturbed by the loss of PRCI*_dnaN_*, suggesting the mobile element is one of multiple causes of increased intracellular stress in this variant.

The ClpP protease is known to inhibit the induction of competence (77), suggesting the increased expression of this protein may link PRCI activation with reducing transformation. Correspondingly, disrupting *clpP* significantly increased the transformation efficiency of RMV7_wt_ (Fig. 4B). Removing PRCI*_dnaN_* in this *clpP*^−^ background did not elevate the transformation efficiency further, consistent with the effects of mobile element activation being mediated through this protease. Disruption of *hrcA* caused a smaller, but still significant, rise in transformation efficiency, with limited evidence of a further increase in the double mutants *hrcA*::*cat* PRCI*_dnaN_*::Janus and *hrcA*::Janus *clpP*::*cat* (Fig. 4B). This suggests the inhibition of transformation driven by PRCI*_dnaN_* primarily reflected its stimulation of increased ClpP activity, with HrcA also likely contributing independently. This is consistent with the *hrcA*::Janus *clpP*::*cat* double mutant exhibiting a stronger growth defect than either single mutant, indicating the proteins have non-redundant functions (Fig. S41). Hence increased intracellular stress driven by MGE activity represses competence through at least one chaperone-mediated pathway.

### Activation and repression of competence by the chaperone regulator HrcA

The lower activity of PRCI*_dnaN_* in RMV7_rare_ suggested that ClpP would be less active in this variant. Correspondingly, disruption of *clpP* caused a smaller growth defect in this variant (Fig. S42), and only a three-fold increase in transformation efficiency, as compared to the >100-fold increase observed in RMV7_wt_ (Fig. 5A). Yet transformation efficiency decreased in RMV7_rare_ following the disruption of *hrcA*, contrasting with the same mutation reproducibly causing increased transformation efficiency in RMV7_wt_ (Fig. 4B, 5A). Furthermore, the RMV7_rare_ *hrcA*::Janus *clpP*::*cat* double mutant also exhibited an elevated transformation efficiency, suggesting this effect of HrcA dominated that of ClpP in this phase variant.

**Figure 5.**
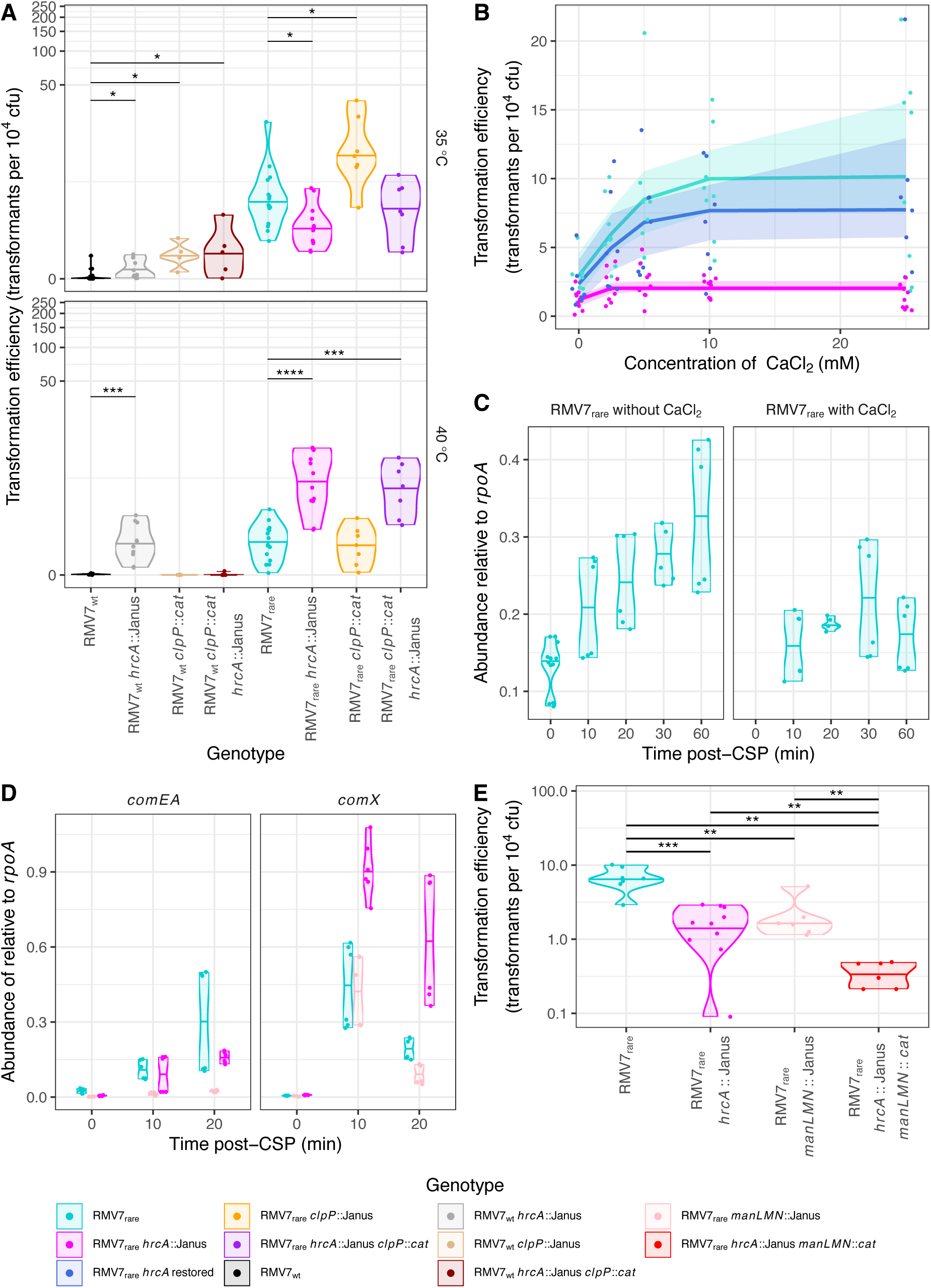
The regulation of transformation by Ca^2+^ and heat shock in RMV7. (A) Violin plots showing the transformation efficiency of RMV7_wt_, RMV7_rare_ and mutant derivatives in which *hrcA* and *clpP* were disrupted during normal growth (35 °C) or during a 40 °C heat shock. Each point corresponds to an independent transformation experiment, and the violin plots have a horizontal line indicating the median transformation efficiency of each mutant at each temperature. (B) Scatterplot showing the dose-dependent effect of CaCl_2_ on transformation efficiency in RMV7_rare_ (cyan) and mutants in which *hrcA* had been disrupted (magenta), and then restored (dark blue). Each point represents an independent transformation assay of one genotype at the indicated CaCl_2_ concentration. The best-fitting dose response logistic models are shown, with the shaded areas corresponding to the 95% confidence intervals. (C) Expression of *hrcA*, measured as transcript abundance relative to *rpoA* by qRT-PCR, following the exposure of cells to CSP. CSP stimulated higher expression of *hrcA* unless co-administered with 12.5 mM CaCl_2_, demonstrating that CaCl_2_ addition affects HrcA regulatory activity. (D) Expression of the early competence gene *comX* and late competence gene *comEA* following the addition of CSP in RMV7_rare_ (cyan), RMV7_rare_ *hrcA*::Janus (magenta) and RMV7_rare_ *manLMN*::Janus (peach), measured relative to the abundance of *rpoA* by qRT-PCR. (E) Combined effects of chaperone and carbon source regulation on transformation efficiency. Results are displayed as in panel A. Wilcoxon rank-sum tests were conducted between all pairs of genotypes. These found both the single mutants, lacking *manLMN* or *hrcA*, were significantly less transformable than the parental genotype. Furthermore, the double mutant was less transformable than either single mutant. Across all panels, significance is coded as: *p* < 0.05, *; *p* < 0.01, **; *p* < 10^−3^, ***; *p* < 10^−4^, ****. All *p* values were subject to a Holm-Bonferroni correction within each panel.

HrcA is unusual in having two conformations (78, 79), only one of which binds the CIRCE DNA motif, enabling autoregulation through repressing the chaperone-encoding gene cluster that includes *hrcA* (80). In *S. pneumoniae*, HrcA binding of CIRCE is reduced at elevated temperatures, relieving its repression of the *dnaK* and *groEL* operons, enabling a heat shock response (81). By contrast, Ca^2+^ ions facilitate HrcA-CIRCE motif binding, inhibiting chaperone expression (79). Both low Ca^2+^ concentrations and extreme temperatures inhibit the induction of competence (82). Hence the divergent effects in the phase variants could reflect the two conformations of HrcA having different effects on the regulation of transformation.

The DNA-binding conformation of HrcA was active in RMV_rare_, as *dnaK* and *groEL* expression was increased in the *hrcA*::Janus genotype (Fig. S43). There was no change in transcription of *clpP*, which is outside the HrcA regulon (Fig. S41). Increasing the proportion of the DNA-binding conformation through supplementation with CaCl_2_ increased the transformation efficiency of RMV7_rare_ in a dose-dependent manner (Fig. 5B). This response was lost in the *hrcA*^−^ genotype, and regained when *hrcA* was restored (Fig. 5B). These effects could be reproduced following gene disruption and restoration in *S. pneumoniae* R6 (Fig. S44). Hence HrcA aids the activation of transformation when adopting the DNA-binding conformation facilitated by Ca^2+^ ions.

As Ca^2+^ increases HrcA’s DNA binding ability, the regulation of transformation seemed likely to occur through altering transcription. Quantifying the expression of *hrcA* post-CSP with and without CaCl_2_ supplementation confirmed the ion concentrations added altered HrcA’s autorepressive activities (Fig. 5C), and decreased expression of the regulon representative *dnaK* (Fig. S45). In contrast to normal growth, the reduction in *hrcA* transcription was associated with slightly increased *clpP* expression, as a potential compensatory mechanism (Fig. S41). However, these changes were primarily manifest 20-60 min post-CSP administration, suggesting the transcriptional effects were too slow to affect competence induction. Correspondingly, neither the early competence gene *comX*, nor the late competence gene *comEA*, showed a decreased response to CSP in the *hrcA*::Janus mutant (Fig. 5D). In contrast, strong suppression of *comEA* transcription was evident in the *manLMN* mutant. This suggests HrcA regulates competence through protein-protein interactions.

As both ClpP and HrcA have roles in thermotolerance, to test whether they caused the reduction in transformation efficiency associated with elevated temperatures, the transformation efficiencies of RMV7_rare_ and RMV7_wt_ *hrcA*^−^ and *clpP*^−^ mutants were compared at 40 °C (Fig. 5A). This heat shock caused a growth defect (Fig. S42), and decreased transformation efficiency, in both RMV7_rare_ and RMV7_wt_. The disruption of *clpP* had no significant effect on transformation efficiency in either variant under these conditions, whereas the disruption of *hrcA* significantly increased transformation efficiency in both phase variants after the heat shock (Fig. 5A). Restoration of *hrcA* reversed the phenotypic change caused by the gene disruption (Fig. S46). This is consistent with HrcA repressing the induction of competence when intracellular stresses shift the protein away from its DNA- binding conformation. This could also explain the phenotypes of the double mutants. The more stressed RMV7_wt_ *hrcA*::Janus *clpP*::*cat* genotype was not transformable, as it exhibited a severe growth defect. By contrast, the RMV7_rare_ *hrcA*::Janus *clpP::cat* double mutant transformed and grew at similar rates to the *hrcA*::Janus single mutant, suggesting a lower dependence on ClpP in the absence of a highly-active PRCI (Fig. S41). Hence HrcA is a pleiotropic regulator that can activate competence in healthy cells, but represses it in response to stress.

### Independent pathways limit the competent cell subpopulation

To test whether ManLMN and HrcA separately contributed to the difference between the variants, the transformation efficiency of the RMV7_rare_ *hrcA*::Janus *manLMN*::*cat* double mutant was compared with that of the progenitor genotype, and the single mutants (Fig. 5D). This demonstrated an approximately five-fold decrease in transformation for each single mutant, and a ~25-fold reduction for the double mutant. This is consistent with HrcA and ManLMN both regulating competence independently.

Experiments with two unlinked resistance markers were used to test whether the difference in competence between variants reflected a uniform reduction in DNA import across cells, or an alteration in the fraction of cells in which competence was induced. The excess of double mutants, relative to the expected frequency inferred from the single mutants (Fig. S47), demonstrated the latter explanation accounted for the distinct behaviours of the variants.

Most bacteria remained recalcitrant to CSP in both variants: with GlcNAc supplementation, it was estimated that 1-2% of the RMV7_rare_ population became competent for transformation, whereas 0.5% of the RMV7_wt_ population did under the same conditions (Fig. S47). Hence HrcA and ManLMN independently changed the probability of an individual cell entering the competent state.

## Discussion

This analysis highlights important challenges in the functional genomic characterisation of clinical pathogen isolates. The RMV7_domi_ and RMV7_rare_ phase variants differed by few polymorphisms, and have the same gene content, with the key differences corresponding reversible DNA excision-reintegration dynamics and alterations to methylation. Yet they exhibited distinct phenotypes that confounded the interpretation of multiple genetic experiments. These differences were attributed to the methylation patterns driven by the *Spn*IV system, as the transformation efficiency of RMV7_wt_ increased when the *tvr* locus was removed, and fell when the *tvr*_domi_::Janus locus was reinstated. Furthermore, the phenotype could not be attributed to mutations outside of the the *tvr* locus. Yet the mechanism linking the epigenetic cause with the phenotypic consequences was difficult to establish, as the effects of DNA methylation were not primarily manifest at genes with modified bases in their promoters. This is consistent with the effects of *Spn*III methylation at the capsule polysaccharide synthesis locus, despite the lack of nearby modification sites (27, 30), and genome-wide analyses of the effects of methylation in other species (48). Instead, it is likely that afflicted genes have an intrinsic sensitivity to intracellular perturbations. For instance, the contribution of the type 1 pilus to biofilm formation were only detectable in RMV7_rare_, while a previous study found the effects of the pilus on the same phenotype differed between a wild-type bacterium and an unencapsulated derivative (83). Hence the substantial impact of methylation variation on competence induction suggests a sensitivity to small genetic, epigenetic and physiological changes that likely underlies its heterogeneity across populations (9, 10, 84), and over the history of individual strains (8, 85).

This susceptibility to variation is likely symptomatic of the many pathways that regulate this phenotype (Fig. 6). One of the important signals identified in this analysis was the availability of GlcNAc, likely the most abundant non-glucose carbon source in the nasopharyngeal mucosa, reaching concentrations similar to the supplements in this work (86). GlcNAc can be liberated from host mucins by pneumococcal glycosylases (87), and used as a carbon and nitrogen source either for growth or metabolism, making it a highly informative signal of nutrient availability and cell physiology (88). Hence GlcNAc-6-phosphate is a regulatory molecule recognised by proteins in some species (89), including *V. cholerae* (90), in which GlcNAc also modulates the induction of competence by a quorum sensing signal (Fig. 6). This analysis identified similarity with components of the *V. cholerae* GlcNAc-signalling pathways in pneumococci, including TfoX, found in many bacterial phyla (Fig. S48), and YjbK, which belongs to a recently-defined subset of CYTH proteins (91, 92) of unknown function in gram-positive bacteria (74, 93). Despite their highly-conserved nature, the corresponding genes were not substantially upregulated by CSP (Fig. S49). Hence these proteins are likely to regulate multiple aspects of pneumococcal physiology, rather than being specific regulators of transformation.

**Figure 6.**
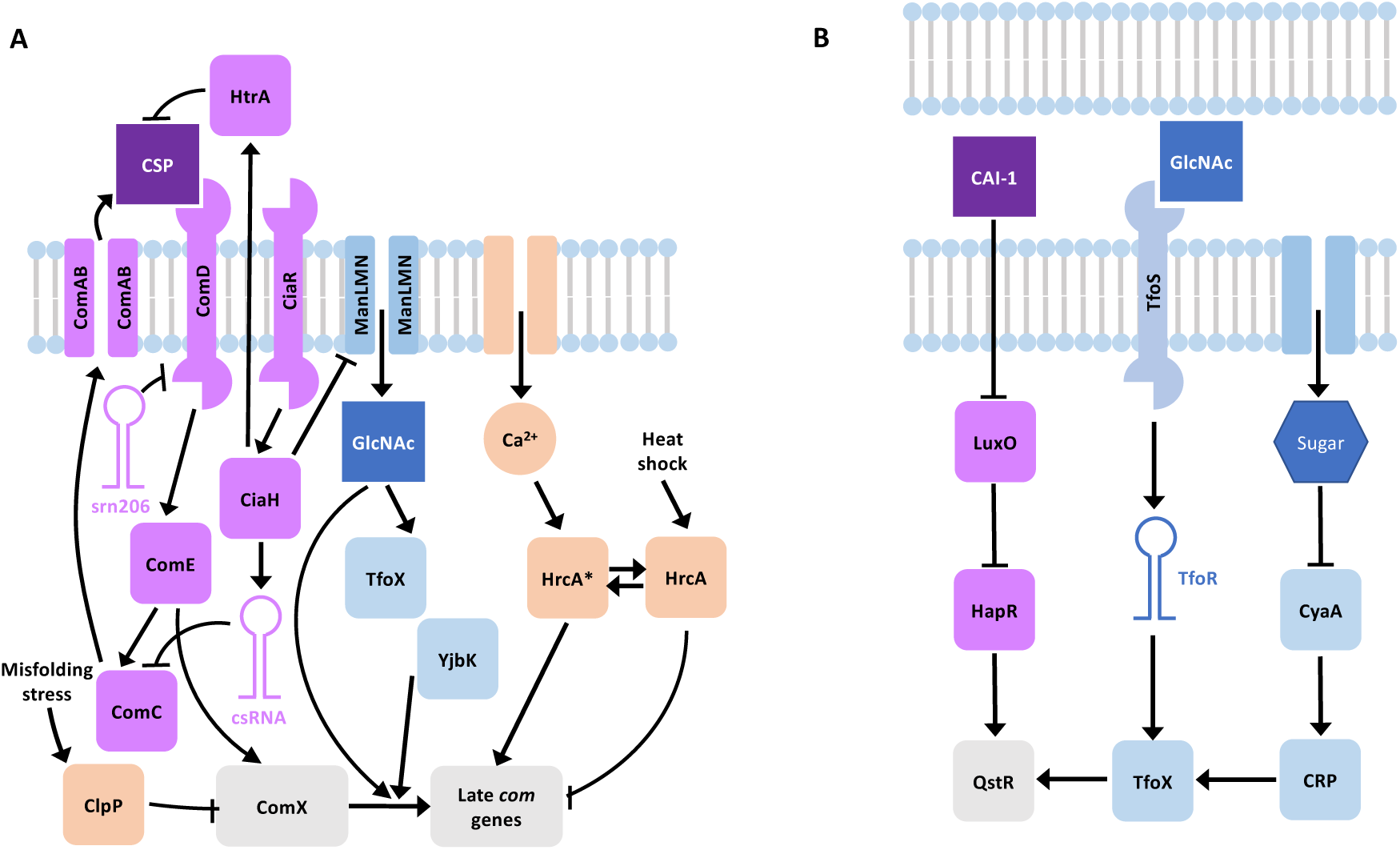
Comparison of the regulation of competence in (A) *S. pneumoniae*, from this work, and (B) *V. cholerae*, summarized from (140). In each, competence is regulated by a quorum-sensing system: CSP in *S. pneumoniae*, and the cholera autoinducer 1 (CAI-1) in *V. cholerae*. The production of CSP is known to be inhibited by the non-coding csRNA, and the HtrA protease derades the signal. Similarly, the srn206 non-coding RNA represses the ComD receptor of CSP (141). CAI-1 operates through inhibiting LuxO, thereby activating the HapR protein, which indicates a high-cell density environment. HapR activates competence through the Quorum-Sensing and TfoX-dependent Regulator (QstR). This regulator also senses the activation of TfoX in response to GlcNAc being sensed by the transmembrane regulator TfoS, via the TfoR small RNA. TfoX activity is also regulated by the Catabolite Regulatory Protein (CRP), which is activated by 3’,5’-cAMP, generated by the CyaA adenylate cyclase under carbon source starvation conditions. Hence there are parallels with the TfoX orthologue, and adenylate cyclase-like protein YjbK, responding to GlcNAc in *S. pneumoniae* RMV7. GlcNAc also appears to promote competence through a TfoX/YjbK-independent route, based on the behaviour of RMV7_wt_. There are no known parallels in *V. cholerae* for the regulation of competence by the chaperones ClpP and HrcA. ClpP represses the induction of competence in response to stresses, such as MGE replication, and has been previously shown to degrade ComX (142). One of HrcA’s two conformations appears to have a similar role, repressing competence in response to heat shock, albeit through a different mechanism. The other active conformation of HrcA, denoted HrcA*, appears to activate competence in response to Ca^2+^, again through an unknown mechanism.

The effects of TfoX and YjbK were dependent on the primary glucose transporter and central metabolic regulator ManLMN (94). As ManLMN is the only effective route by which GlcNAc can be imported (64), signalling by this molecule is limited by the competence regulator CiaRH, but not subject to carbon catabolite repression. Orthologues of ManLMN serve as the main glucose transporter across many Firmicutes, including other streptococci, *Lactococcus lactis* and *Listeria monocytogenes* (64), and the transporter has been associated with regulation of biofilm formation and transformability in *Streptococcus mutans* (95) suggesting its signalling role is likely to be common among Firmicutes. Hence the heterogeneity of pneumococcal competence induction partly reflects highly-conserved metabolic signalling networks intervening in competence-specific pathways.

HrcA is another widely-conserved key regulator of pneumococcal physiology that this analysis found to regulate competence induction. This protein is sensitive to physiologically-relevant concentrations of CaCl_2_ (96, 97), the most common ionic compound in the nasopharyngeal mucosa (86). Hence rather than Ca^2+^ aiding the translocation of DNA molecules across the plasma membrane, as suggested previously (98), HrcA appears to mediate a rare example of Ca^2+^ signalling in bacteria (79, 99). Another unusual aspect of HrcA’s activity is that its two conformations appear to have opposing effects on competence activation, resulting in the regulator’s phenotype depending on the epigenetic context of the cell. Hence HrcA is able to modulate competence induction through integrating information on intracellular stress and extracellular ion concentrations. The mechanism by which this was achieved is not clear, but is more likely to involve protein-protein interactions than DNA binding, given the slow transcriptional responses to Ca^2+^ administration, and the lack of change in competence gene expression in an *hrcA*^−^ genotype. This contrasts with the independent effects of ManLMN, which limited the induction of late competence genes by ComX, meaning general regulators of cell biology affect multiple steps of the competence regulatory cascade (Fig. 6).

The structure of this regulatory network can help explain the paradox of two common, but contrasting, aspects of competence regulation (100): quorum sensing, which drives coordinated responses, and bet hedging, which underlies population-wide heterogeneity. In isolated cells, bet hedging could result from intrinsic noise, the variation in gene activity reflecting the inherent stochasticity of transcription and translation (101). Yet competent pneumococci increase their production of CSP, propagating induction to neighbouring cells (102), tending to homogenise the population-wide response. Maintaining heterogeneity therefore requires cells within a clonally-related population, adapted to the stable niche of the nasopharynx, stochastically differ in their susceptibility to the quorum sensing signal.

The examples of HrcA and ManLMN demonstrate how this is achieved by pneumococci. Firstly, both leverage variation in cell clusters’ microenvironments and intracellular physiology (101) as sources of stochasticity, or extrinsic noise (103). Secondly, both mechanisms act on steps of the regulatory cascade that do not affect CSP production, meaning their effects occur independently within cells, and are not propagated intercellularly. Thirdly, both act on independent steps of the cascade, rather than being integrated into a common mechanism, enabling them to have uncorrelated, independent effects on competence induction. Fourthly, although temperature, and the availability of Ca^2+^ and GlcNAc, substantially altered the probability of competence induction, none alone had a large enough effect to risk the entire population becoming competent, maintaining heterogeneity in the population. Hence bet hedging can emerge as an apparently random output of combining multiple noisy signals, with the phase-variable *Spn*III and *Spn*IV restriction-modification systems further potentiating such variation (27, 57). The function of CSP is therefore not to homogenise the population, but to coordinate the induction of this transient state in a subset of pneumococci.

This complex regulation in *S. pneumoniae* is similar to that in *Bacillus subtilis* (104), as well as *Vibrio cholerae* (105) and some other gram-negative bacteria (106). These distantly-related species all have strongly-inducing quorum sensing signals that rapidly induce a transient competence state in a subset of bacteria, with coordinated release of DNA from conspecific cells through fratricide or cannibalism (107–109). In each case, the intercellular signalling is modulated by multiple extracellular stimuli, although the signals themselves vary between the bacteria, likely driven by their divergent ecologies. Hence *V. cholerae* responds to chitin (66), while *B. subtilis* induces competence under starvation conditions (108), whereas this work demonstrates that pneumococcal competence is favoured in healthy bacteria, replete with host-derived nitrogen, carbon and ion sources. Instead, it is the complex structure of the regulatory network that is shared, enabling environmental variation and stochasticity of signalling to modulate responses to quorum sensing. Hence some naturally transformable bacteria appear to have convergently evolved noisy regulatory systems that stochastically segregate populations into donors and recipients, thereby enabling the efficient transfer of DNA during a transient period of competence.

Other naturally transformable bacteria are either constitutively competent, in the case of some *Neisseria* species (110), or do not employ quorum sensing, as appears to be the case for *H. influenzae* (111). However, the import of DNA is constrained to molecules containing DNA uptake sequences (DUSs) in these bacteria (110, 112). Therefore it has been proposed that the purpose of the transient nature of competence by quorum sensing is to synchronise the release and acquisition of DNA from conspecific bacteria, thereby serving as an alternative mechanism to DUSs for ensuring imported genetic material comes from close relatives (113). Hence it is highly unlikely the competence system primarily functions to acquire nucleic acids as a source of nutrients (114–116), as both common types of competence system limit their imported DNA to sequences that are sufficiently closely-related to be integrated through homologous recombination. Rather, in species regulating competence using quorum sensing, multiple signals are likely used to ensure both the coordination of induction, and the emergence of population-level heterogeneity. Hence the variable nature of species-wide transformation efficiency represents the delicate balance between chaos and order necessary for the synchronised bet hedging that characterises competence in many bacterial species.

## Methods and materials

### Cell culture

Genotypes used in this study are described in Table S5. Unless otherwise stated, encapsulated *S. pneumoniae* were cultured statically at 35 °C in 5% CO_2_. Culturing on solid media used Todd-Hewitt broth supplemented with 0.5% yeast extract and 1.5% agar (Sigma-Aldrich). Media were supplemented antibiotics for selection of mutated genotypes: rifampicin (Fisher Scientific) at 4 μg mL^−1^; kanamycin (Sigma-Aldrich) at 400 μg mL^−1^, or chloramphenicol (Sigma-Aldrich) at 4 μg mL^−1^. Phase contrast microscopy of colonies used a Leica DFC3000 G microscope.

Unless otherwise specified, culturing in liquid media used 10 mL of a mixture of a 2:3 ratio of Todd-Hewitt broth (Sigma-Aldrich) with 0.5% yeast extract (Sigma-Aldrich), and Brain-Heart Infusion media (Sigma-Aldrich); this referred to as “mixed” media. Transformation experiments with *S. pneumoniae* R6 derivatives used a chemically-defined medium, consisting of disodium β-glycerophosphate (20 g L^−1^; Sigma-Aldrich), sodium pyruvate (0.1 g L^−1^; Fluorochem), choline (0.001 g L^−1^; Alfa Aesar), cysteine (0.4 g L^−1^; Tokyo Chemical Industry UK), glucose (3.8 mM; Sigma-Aldrich) and galactose (12 mM; Sigma-Aldrich). Carbon source supplements were added to liquid media at a final concentration of 30 mM, unless otherwise specified.

### Growth curves and surface adhesion assays

To measure growth curves, 2×10^4^ cells from titrated frozen stocks were grown in mixed liquid media in 96-well microtiter plates at 35°C with 5% CO_2_ for 20 h. Measurements of OD_600_ were taken at 30 min intervals over 16 hours using a FLUOstar Omega microplate reader (BMG LABTECH). Three replicate wells were assayed for each tested genotype in each experiment. The R package growthcurver was used for the inference of carrying capacity, *K*, and replication rate, *r* (117). For measuring adhesion to abiotic surfaces, at the end of the growth curve incubation, the microtitre plate was submerged in water and dried for 10 min. Then 125 μl of a freshly-diluted 0.1% crystal violet solution (Scientific Laboratory Supplies) was added to each well, followed by incubation for 30 minutes at room temperature. Each well was then washed by repeatedly submerging the plate in water to remove excess crystal violet. The plate was incubated at room temperature in an inverted position for four hours. Subsequently, 125 μl of 30% acetic acid (Honeywell) was added to each well. Adherence was quantified as OD_550_ across replicate wells, measured by a FLUOstar Omega plate reader.

### Mutagenesis

Disruption of genes for directed mutagenesis required the PCR amplification of 0.8-1 kb regions flanking the gene of interest, using the oligonucleotide sequences listed in Table S6. For each gene, the upstream region was amplified with the oligonucleotides labelled with the gene name and the suffixes “Up_For” and “Up_Rev_ApaI”, with the latter adding an *Apa*I site to the 3’ of the amplicon. The downstream region was amplified with the oligonucleotides labelled with the gene name and the suffixes “Down_For_BamHI” and “Down_Rev”, with the former adding an *Bam*HI site to the 5’ of the amplicon. The corresponding antibiotic resistance markers were amplified with oligonucleotides that added flanking *Bam*HI and *Apa*I restriction enzyme sites: Janus_For_ApaI and Janus_Rev_BamHI for the Janus cassette, or Cat_For_ApaI and Cat_Rev_ApaI for the *cat* chloramphenicol resistance marker (35). PCR products were digested with the appropriate restriction enzymes (Promega) at 35 °C for 2-4 hours, and then ligated to the appropriate antibiotic marker using T4 DNA ligase (Invitrogen). Ligation mixtures were used as templates to amplify correctly-ligated constructs through PCR using the “Up_For” and “Down_Rev” oligonucleotides, which generated the amplicons used for mutagenesis through transformation of competent pneumococcal cells.

Restoration of genes disrupted using the Janus cassette depended on all genotypes in which mutations were generated originally being resistant to streptomycin (Table S5), owing to a mutation in *rpsL* (118). Mutants were isolated in which the Janus cassette restored susceptibility to streptomycin. This enabled the Janus cassette to be removed through transformation with PCR amplicons, encoding the intact gene, generated from the original genotypes with the “Up_For” and “Down_Rev” oligonucleotides, using selection for streptomycin-resistant cells. As intragenomic recombination generates false positive streptomycin-resistant genotypes independently of transformation, identifying true positive recombinants in which genes were restored through eliminating the Janus cassette was only feasible in the more transformable RMV7_rare_ variant. PCR amplification was used to ensure the complete removal of the cassette.

The exception to this approach was the construction of RMV7 *tvr*_rare_::Janus and RMV7 *tvr*_domi_::Janus. Both genotypes derived from the RMV7_wt_ *tvr*::*cat* mutant, which was transformed with DNA from RMV7_domi_ or RMV7 *tvr*_rare_ *tvrR*::Janus, followed by selection on plates supplemented with kanamycin. The RMV7 *tvr*_rare_ *tvrR*::Janus genotype was the original locked phase variant from which RMV7_rare_ was derived (35), through removal of the Janus cassette. Using RMV7 *tvr*_rare_::Janus as the donor in this transformation allowed kanamycin selection to be used to isolate transformed genotypes that had acquired the modified *tvr* locus. Notably, the original isolation of different phase variants through disruption of *tvrR* used different constructs, and therefore RMV7_domi_ was locked through the replacement of 681 bp from the 3’ end of *tvrR*, whereas RMV7 *tvr*_rare_ *tvrR*::Janus was locked through the replacement of 267 bp from the 5’ end of the gene (Fig. 1). Hence the comparison of RNA-seq data across the two variants identified differential levels of sequence read mapping to *tvrR* (Fig. 2).

### Transformation assays

One milliliter of bacterial culture was collected at an optical density of 600 nm (OD_600_) between 0.15 and 0.25. Cells were then incubated for 2 hours at 35 °C with 5 μl of 500 mM CaCl_2_ (Sigma-Aldrich), 250 ng of competence stimulating peptide 1 (CSP-1; Cambridge Bioscience Ltd) and 100 ng of the purified *rpoB* gene, containing a base substitution that conferred resistance to rifampicin (118). In experiments using carbon source supplements, these were added at a final concentration of 33 mM. After two hours of incubation at 35 °C, a volume of between 1 and 200 μl of the transformed culture was spread on an agar plate supplemented with rifampicin. For precise quantification of transformation frequencies, 1 μl of a 10^3^-fold dilution of the same culture was spread on a non-selective plate in parallel. Colonies were counted after 24 hours of incubation at 35 °C with 5% CO_2_.

A logistic curve was used to analyse the relationship between transformation efficiency (in transformants per 10^4^ colony forming units), *t*, and the concentration of the CaCl_2_, *x*. The fitted function was:

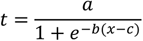

The values of the variables *a*, *b* and *c* were estimated using the Levenberg-Marquardt nonlinear least-squares algorithm in the minpack R package (119), using the starting values of 10 (increased to 500 for *S. pneumoniae* R6), 1 and 0.5, respectively. The confidence intervals were calculated through refitting the function to 999 bootstrapped samples using the car R package (120).

### Estimation of the frequency of competence induction

The calculation of the proportion of cells in which competence was induced required the transformation of cultures with two independent selectable markers. The frequency of transformation with each single marker, *f*, can be expressed as the product of the proportion of pneumococci that are competent for transformation, τ, and the probability of acquisition of each marker: *p*_rif_ for rifampicin, and *p*_kan_ for kanamycin. Hence the product of these two transformation efficiencies, *P*_double_, is:

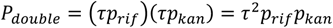

The observed frequency of double mutants, *O*_double_, can be expressed as the product of the probability a cell is competent for transformation, *p*_rif_ and *p*_kan_:

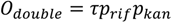

Therefore the proportion of bacteria that are competent for transformation within a culture can be calculated as:

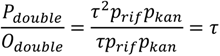

Note that if all cells in the culture are transformable, then *P*_double_ = *O*_double_, whereas an excess of double mutants over the frequency expected from the single mutant frequencies is evidence that only a subset of the population is competent for transformation.

### Preparation of RNA samples and quantitative PCR

Three replicate cultures of RMV7 *tvr*_domi_::Janus and RMV7 *tvr*_rare_::Janus were grown in 25 mL of mixed liquid media until they reached an OD_600_ of 0.15. A 5 mL sample of bacterial cells was collected and 50 μL 100 ng mL^−1^ CSP1 was added to the remaining culture. Further 5 mL samples were taken from each culture 10 and 20 min post-CSP addition. Each sample was immediately treated with 10 mL RNAprotect (Qiagen) and incubated at room temperature for 5 min. Cell were then pelleted by centrifugation at 3,220 *g* for 10 min. RNA was extracted from the washed pellets using the SV Total RNA Isolation System (Promega) according to the manufacturer’s instructions. The extracted RNA was used for RNA sequencing or qRT-PCR.

All qRT-PCR experiments were conducted as described previously (35), using 0.2 μg of RNA to generate cDNA with the First-Strand III cDNA synthesis kit (Invitrogen). Reactions used the PowerUp™ SYBR™ Green Master Mix (Thermo Fisher) and the QuantStudio™ 7 Flex Real-Time PCR System (Applied Biosystems). All experiments use at least two independent biological replicates, each of which was analysed with three technical replicates.

Gene expression was quantified used standard curves. Genomic DNA was diluted to copy numbers of 3×10^3^, 3×10^4^, 3×10^5^ and 3×10^6^, and cycle threshold (Ct) values were calculated for *rpoA* and the gene of interest for each. The standard curve was used to convert Ct values into absolute copy numbers. This allowed the quantification of expression as a relative abundance of each gene of interest relative to *rpoA*.

To quantify the dynamics of *tvr* loci during culture, three independent passages of RMV7_wt_ in liquid media were each initiated through inoculation with a single colony. Each subsequent 24 h passage was inoculated with 10 μL of the previous culture. Genomic DNA was extracted from the initial culture, and after four and eight days of passage, using the Wizard Genomic DNA purification kit (Promega). Quantitative PCR was used to measure the frequency of each distinguishable allele of the *tvr* locus, using triplicate technical replicates on each sample, as described previously (121). The absolute copy numbers of each allele were then estimated using standard curves, thereby enabling the *tvr*_domi_:*tvr*_rare_ ratio to be calculated.

### Generation and analysis of RNA-seq data

RNA samples were quantified using an Agilent Bioanalyser RNA Nano Chip (Agilent Technologies), and treated with the RiboZero® rRNA Removal Kit for Bacteria (Illumina) to deplete rRNA. The samples were then cleaned with Agencourt RNAClean Beads (Beckman Coulter). Sequencing libraries were generated with the NEBNext® Ultra II Directional Library Prep Kit for Illumina (New England BioLabs), modified to use oligonucleotide sequences appropriate for the sequencing pipelines of the Wellcome Sanger Institute. The library was amplified through nine PCR cycles using the Kapa HiFi HotStart Ready Mix (Roche) to generate sufficient material for sequencing. All eighteen samples were sequenced as a multiplexed library on a single lane of a HiSeq 4000 sequencing system (Illumina), generating 200 nt paired end reads.

The set of genes used for expression analysis were the 2,088 protein coding sequences annotated on the *S. pneumoniae* RMV7_domi_ genome (accession code OV904788), and the 81 non-coding RNAs predicted by infernal version 1.1.2 (122) using the Rfam database (123). RNA-seq reads were aligned to these sequences using kallisto version 0.46.2 (124), using default settings and 100 bootstraps. Differential gene expression analysis used sleuth version 0.30 (125). Wald tests were conducted to compare the pre-CSP samples for RMV7 *tvr*_domi_::Janus and RMV7 *tvr*_rare_::Janus, and to compare the 10 min and 20 min post-CSP samples for each genotype to the corresponding pre-CSP samples. Visualisation and plotting of data used the genoplotR (126), circlize (127), cowplot (128), ggpubr (129) and tidyverse (130) packages.

### Quantification of 3’,5’-cAMP production

Quantification of 3’,5’-cAMP used the Cyclic AMP XP^®^ assay kit (Cell Signalling Technology). Samples were harvested in the exponential (OD_600_ of 0.2) and stationary (OD_600nm_ of 0.5) phases of statically-grown *S. pneumoniae* cultures. *E. coli* DH5⍺ was grown in 25 mL Luria-Bertani media (Sigma-Aldrich) at 37 °C, shaken at 250 revolutions per minute, and samples were harvested in the exponential (OD_600_ of 0.3) and stationary (OD_600nm_ of 1.0) phases. Cells were pelleted through centrifugation at 3,220 *g* for 10 min. The cell pellets were re-suspended in 850 μL of the kit’s lysis buffer and 150 μl of lysozyme (10 mg mL^−1^), followed by incubation at 35 °C for 20 minutes. Cells were pelleted by centrifugation at 9,500 *g* for 5 minutes. The supernatants were collected, and the cell pellets were resuspended in 400 μl phosphate-buffered saline. The protein concentrations of all the collected samples were adjusted to 400 μg mL^−1^ using the Qubit protein broad range assay kits (Qiagen). The Cyclic AMP XP^®^ kit ELISA plates were prepared according to the manufacturer’s protocols, and a 50 μl sample of supernatant from each tested culture was loaded in each well. The OD_450_ was measured using the FLUOstar Omega plate reader. Concentrations of 3’,5’-cAMP were calculated using a standard curve, according to the manufacturer’s instructions.

### Analysis of motif distribution and sequence diversity

The RMV7_domi_ and RMV7_rare_ sequences were aligned with nucmer (131). The overall distribution of methylation motifs was calculated using DistAMo (132). The calculation of distances between coding sequences and motifs used biopython (133).

The *S. pneumoniae* RMV7 TfoX and YjbK proteins were aligned to orthologues from *H. influenzae* Rd and *V. cholerae* ATCC 39315 using Muscle (134). The predicted structures were retrieved from the AlphaFoldDB (135).

Proteins containing the TfoX N-terminal domain were identified using EMBL SMART (136). These amino acid sequences were aligned with MAFFT (137), and a phylogeny generated with Fasttree2 (138) using default settings. Proteins were assigned to bacterial Families using the NCBI Taxonomy (139).

## Acknowledgements

MJK, AVI and NJC were supported by the UK Medical Research Council and Department for International Development (grant nos MR/R015600/1 and MR/T016434/1), and a Sir Henry Dale Fellowship, jointly funded by Wellcome and the Royal Society (grant no. 104169/Z/14/A). MJK and MRO were supported by the BBSRC (grant no. BB/N002903/1). SDB was supported by Wellcome (grant no 206194). We thank the Bespoke team at the Wellcome Sanger Institute for generating the sequencing libraries.

## Data availability

The genome sequence and annotation of *S. pneumoniae* RMV7_domi_ is available from the European Nucleotide Archive (ENA) with the accession code OV904788. The RNA-seq data are available from the ENA with the accession codes listed in Table S2. The expression values and statistical tests calculated for all analysed genes in the RNA-seq analysis are available from FigShare, alongside the raw gel images, micrographs and the results of qRT-PCR and microbiological experiments (https://figshare.com/projects/Diverse_regulatory_pathways_modulate_bet_hedging_of_competence_induction_in_epigenetically-differentiated_phase_variants_of_Streptococcus_pneumoniae/171060), as detailed in Table S7.

