## Supplementary figures and table for "Diverse regulatory pathways modulate “bet hedging” of competence induction in epigenetically-differentiated phase variants of *Streptococcus pneumoniae*"

**Supplementary materials**

Figures S1–S49
Tables S1-S7 (legend only for Table S3)

### Supplementary Figures

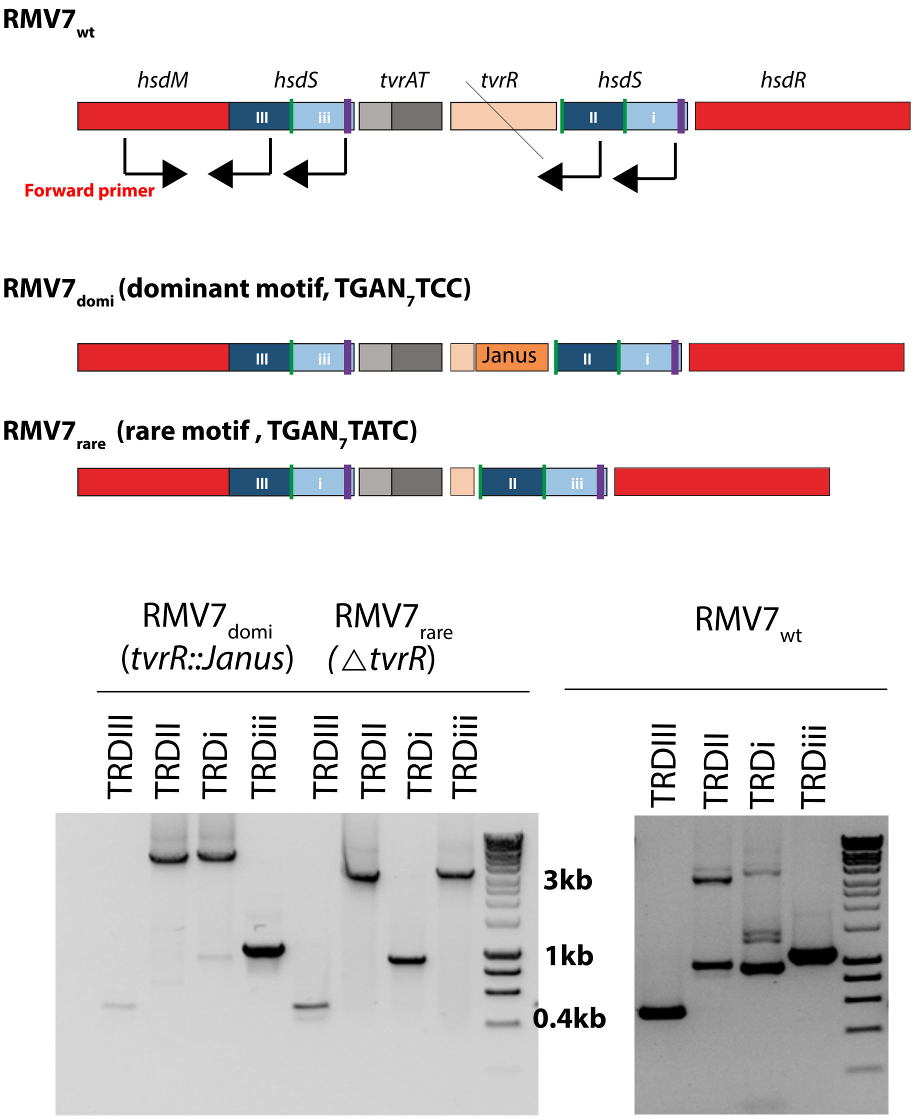

**Figure S1** Establishing the differing arrangements of the *tvr* loci in the RMV7 variants. (A) Schematic showing the binding site of oligonucleotides used for PCR amplification experiments. The forward primer binds an invariant site within *hsdM*. The four different reverse primers each bind a site specific to an individual TRD-encoding sequence, which are rearranged through the integration-excision activity of TvrR. (B) Separate PCR amplification experiments were conducted with the same conserved forward primer, and the four reverse primers, on $RMV7_{wt}$, $RMV7_{domi}$ and $RMV7_{rare}$. In $RMV7_{domi}$, the sequence encoding TRDiii forms part of the active *hsdS* gene, adjacent to *hsdM*, whereas the sequences encoding TRDII and TRDi are in the inactive, distal position. In $RMV7_{rare}$, the sequence encoding TRDi forms part of the active *hsdS* gene, adjacent to *hsdM*, whereas the sequences encoding TRDII and TRDiii are in the inactive, distal position. In $RMV7_{wt}$, which has an active TvrR, both arrangements are evident, as are alternative alleles in which the sequence encoding TRDII is proximal to *hsdM*. The size of products is shown relative to DNA Hyperladder 1kb (Bioline).

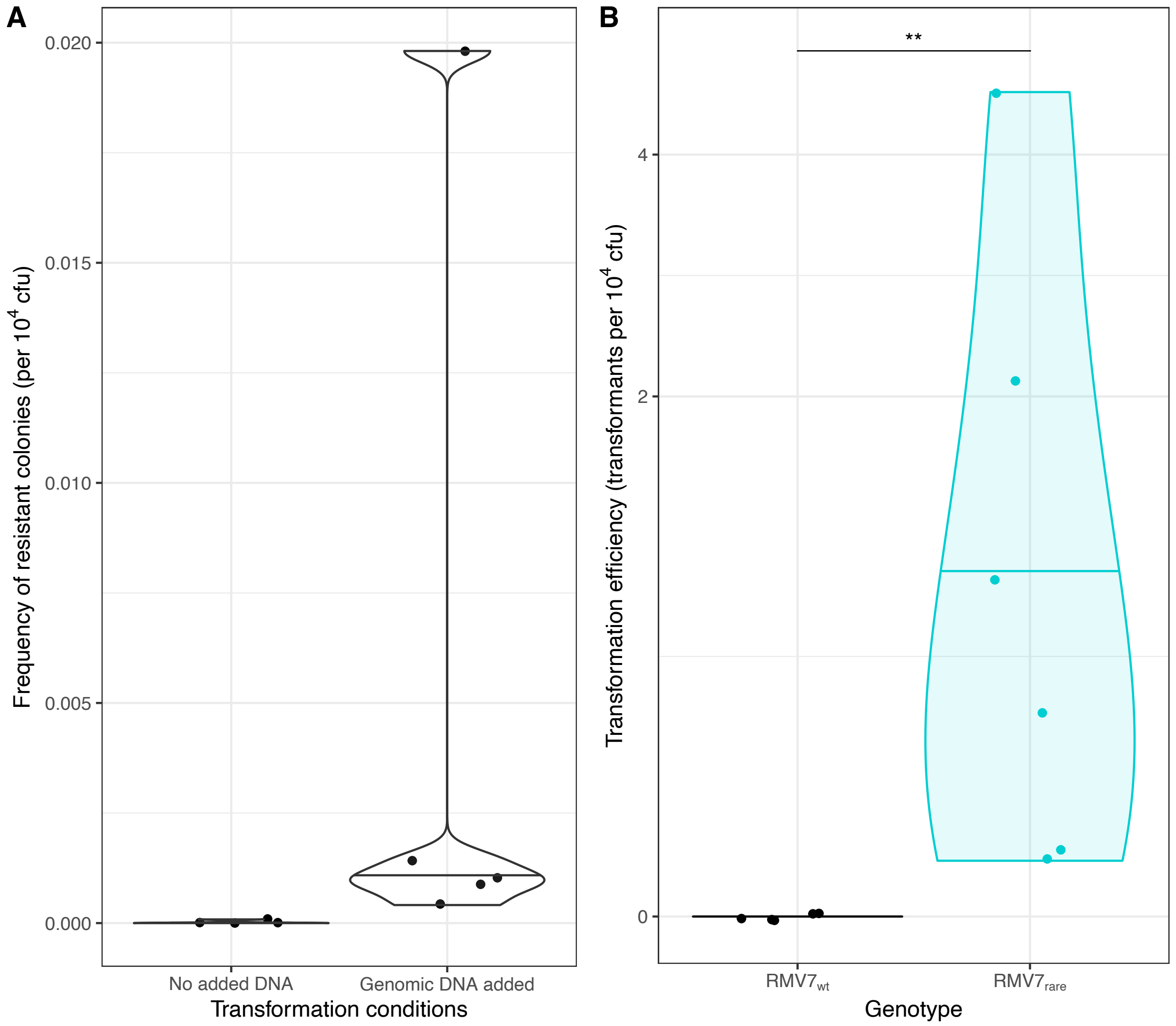

**Figure S2** Validation of differences in transformation between RMV7$_{wt}$ and RMV7$_{rare}$. (A) Frequency of spontaneous rifampicin resistant mutants in RMV7$_{wt}$ observed after performing the transformation protocol, including the addition of CSP, in the presence and absence of exogenous DNA. The frequency of spontaneous rifampicin resistant RMV7$_{wt}$ bacteria was very low in these experiments. This confirmed the majority of resistant bacteria observed in transformation experiments resulted from the uptake of exogenous DNA. (B) Both RMV7$_{wt}$ and RMV7$_{rare}$ were transformed, and the cells were plated on selective mediate after 20 h, rather than 2 h. This tested whether the induction of competence was slower, rather than lower, in RMV7$_{wt}$. A significant difference between RMV7$_{wt}$ and RMV7$_{rare}$ was still observed, confirming the variation recorded in other experiments was not the consequence of delayed competence induction in RMV7$_{wt}$. Significance between results is coded as: $p < 0.05$, *; $p < 0.01$, **; $p < 10^{-3}$, ***; $p < 10^{-4}$, ****.

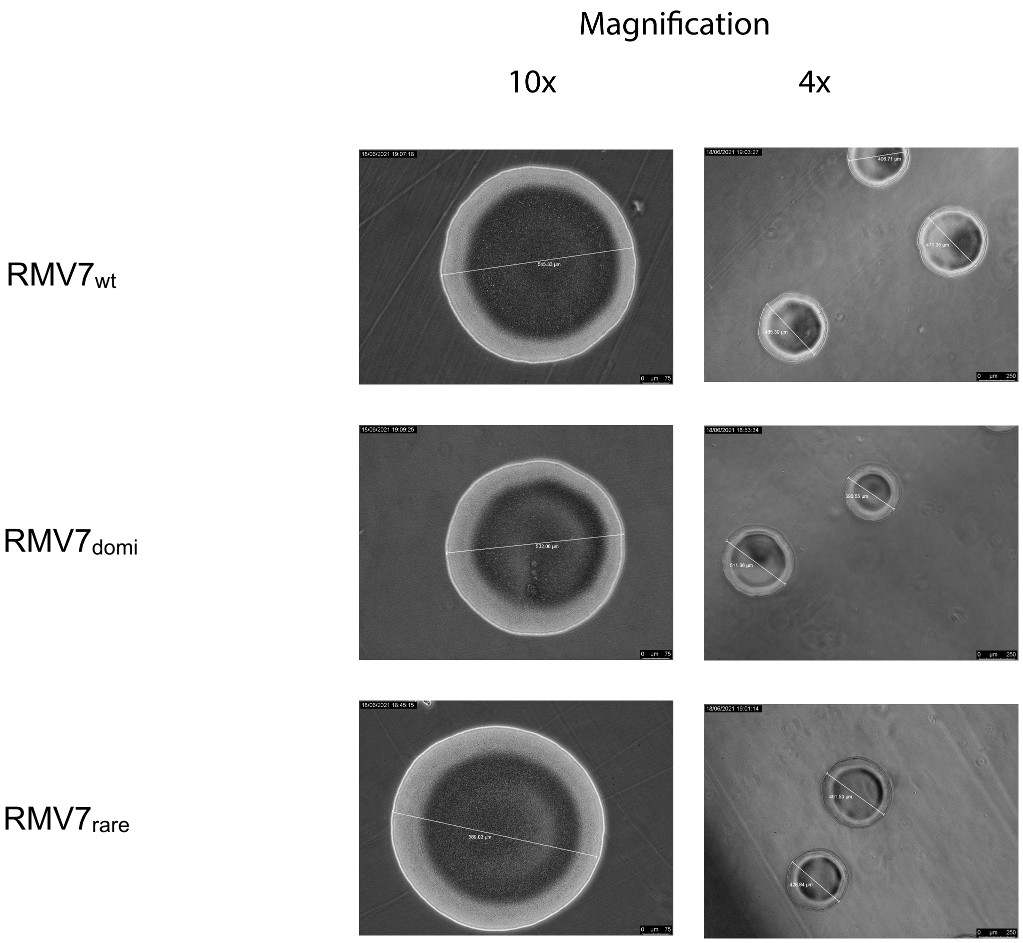

**Figure S3** Morphology of RMV7 variant colonies. Phase contrast microscopy was used to visualize colonies of RMV7$_{wt}$, RMV7$_{domi}$ and RMV7$_{rare}$. All three had a consistent size: the three scale bars spanning the colonies shown at 10x magnification have lengths of 545, 502 and 589 µm, from top to bottom. This is consistent with all genotypes expressing a similar thickness of capsule. This distinguishes them from transparent and opaque phase variants, which exhibit distinctive colony morphologies.

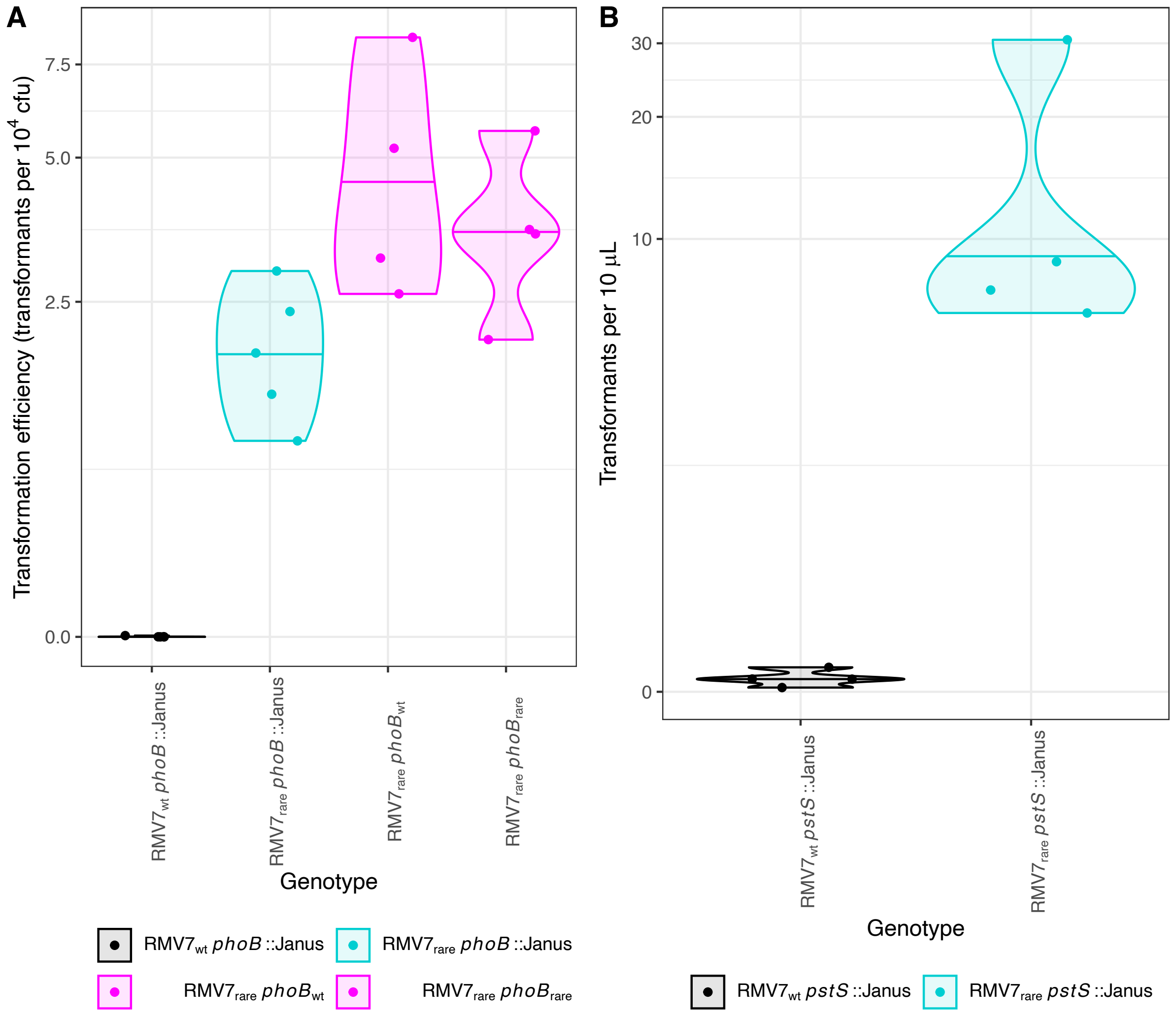

**Figure S4** Testing the effect of mutations outside the *tvr* locus on variant transformation efficiency. (A) Violin plot showing the transformation efficiency of *phoB* knock out and knock in mutants of $RMV7_{wt}$ and $RMV7_{rare}$. Each point represents an independent transformation experiment. The violin plots summarise the transformation efficiency of each genotype, with a horizontal line representing the median transformation efficiency. The maintenance of a ~100-fold difference in transformation efficiency between $RMV7_{wt}$ and $RMV7_{rare}$ in the absence of *phoB* demonstrates the polymorphisms detected within this regulatory gene are unlikely to cause the phenotypic differences between $RMV7_{wt}$ and $RMV7_{rare}$. This is confirmed by the similarity of the transformation efficiency of $RMV7_{rare}$ carrying the knocked in *phoB*$_{wt}$ and *phoB*$_{rare}$ alleles. (B) Violin plot showing the transformation efficiency of *pstS* knock out mutants of $RMV7_{wt}$ and $RMV7_{rare}$. The loss of the *pstS* gene, which contains a premature stop codon in $RMV7_{rare}$, did not affect the difference in transformation efficiency between the variants.

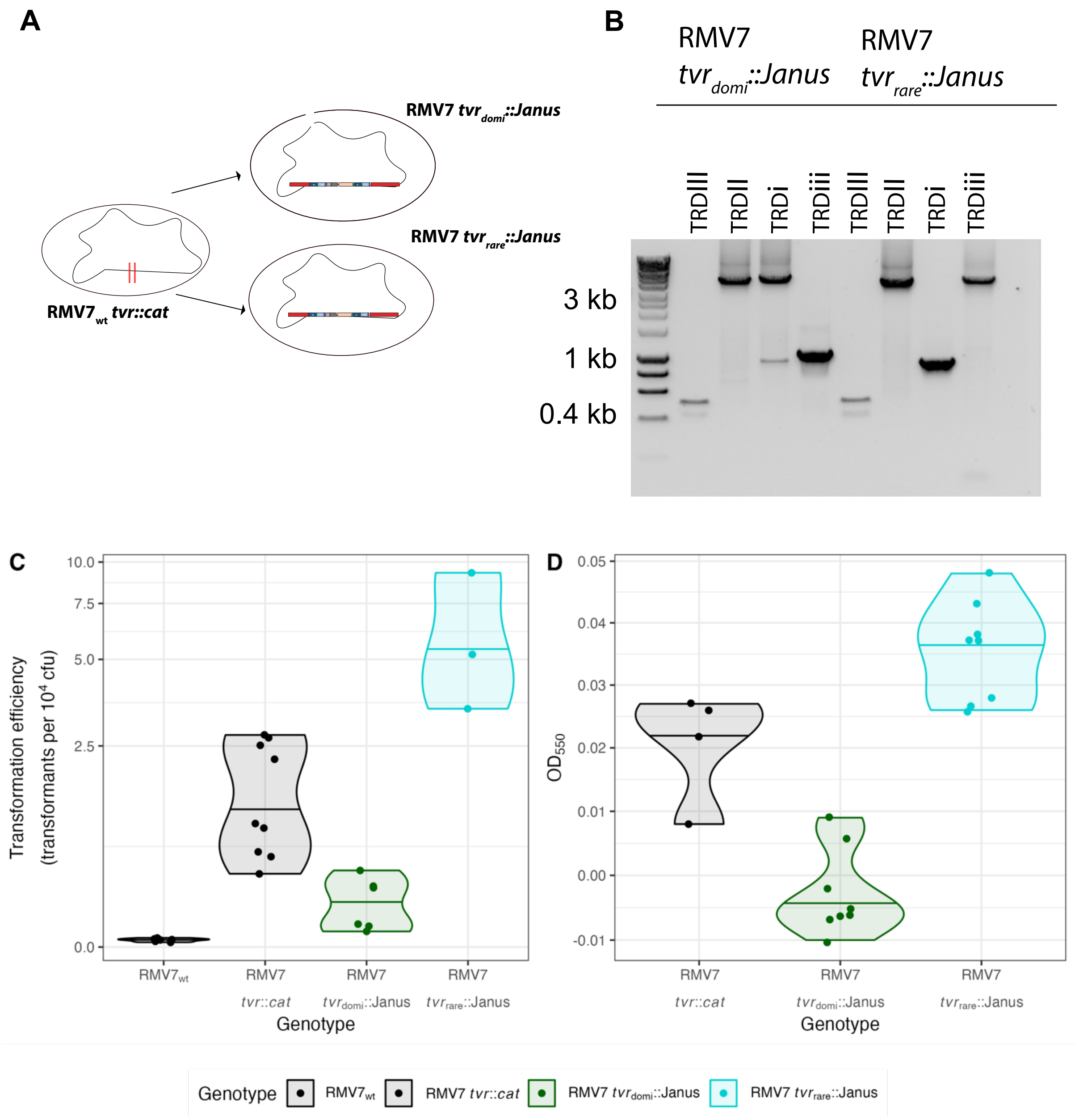

**Figure S5** Design of the RNA-seq and qRT-PCR experiments. (A) The *tvr* loci of RMV7$_{\text{domi}}$, and a version of RMV7$_{\text{rare}}$ that retained a Janus cassette inserted into *tvrR*, were introduced into a common RMV7$_{\text{wt}}$ *tvr::cat* recipient. (B) The agarose gel shows the PCR amplicons generated with the primers described in Fig. S1, which demonstrates the intact *tvr* loci have been integrated into the recipients to generate RMV7 *tvr*$_{\text{domi}}$::Janus and RMV7 *tvr*$_{\text{rare}}$::Janus. The amplicon sizes are measured relative to DNA Hyperladder 1kb. (C) Violin plot showing the transformation efficiencies of genotypes from which RNA was extracted for RNA-seq analysis. These results demonstrated the insertion of the *tvr*$_{\text{domi}}$::Janus and *tvr*$_{\text{rare}}$::Janus loci replicated the divergence in transformation between RMV7$_{\text{domi}}$ and RMV7$_{\text{rare}}$. (D) Violin plot showing the thicknesses of biofilms formed by the genotypes from which RNA was extracted for RNA-seq analysis. These results demonstrated the insertion of the *tvr*$_{\text{domi}}$::Janus and *tvr*$_{\text{rare}}$::Janus loci replicated the divergence in biofilm thicknesses between RMV7$_{\text{domi}}$ and RMV7$_{\text{rare}}$.

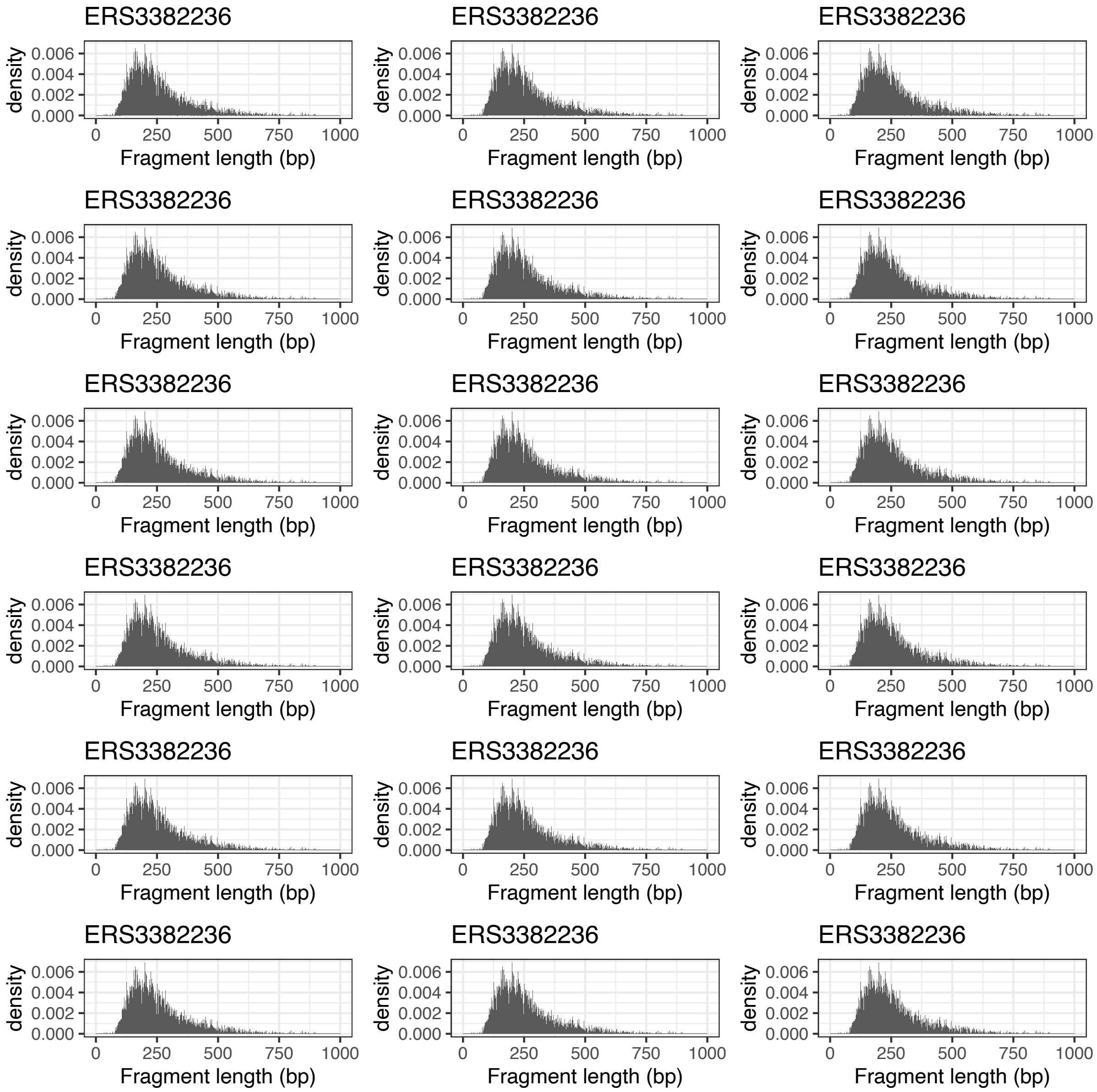

**Figure S6** Density plots showing the inferred distribution of fragment sizes from mapping by Kallisto. The consistency of these distributions between samples, each labelled with its accession code (Table S2), suggests there should not be any bias introduced by differences in sequencing library preparation.

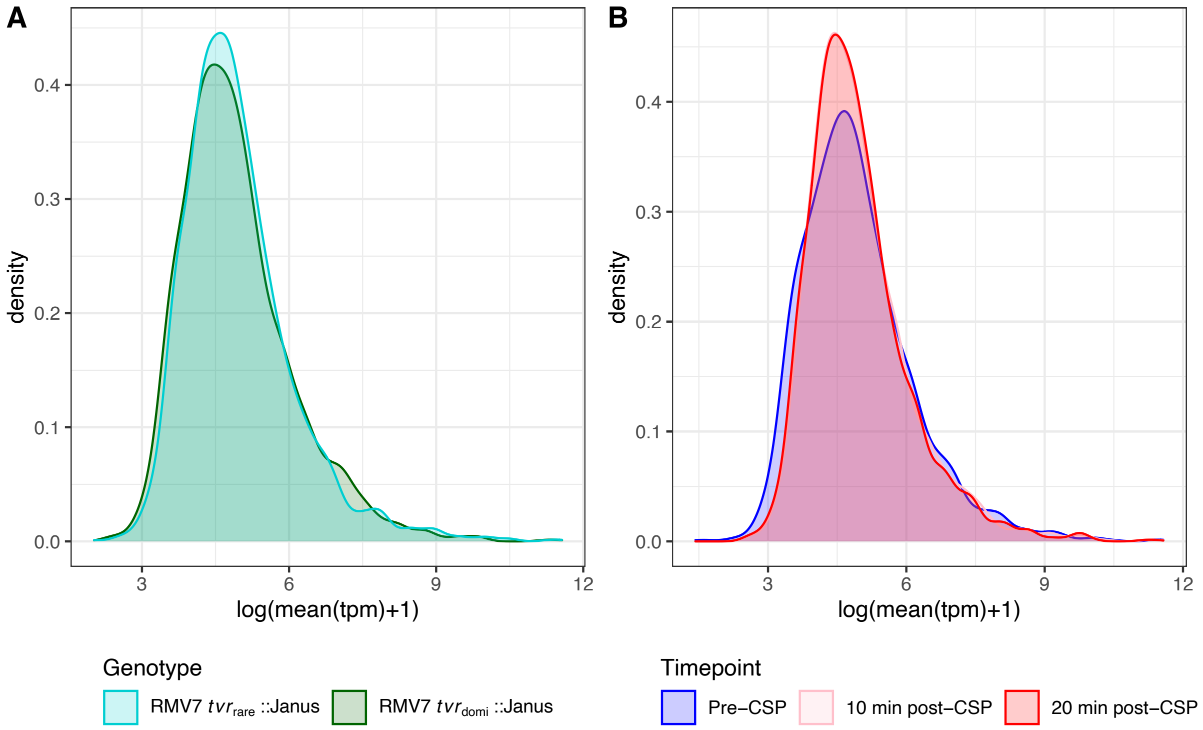

**Figure S7** Density plots showing the distribution of mean transcripts per million (tpm) values across coding sequences for samples in different groups. (A) This density plot shows the mean tpm distribution is highly similar between samples originating between the two genotypes. (B) This density plot shows the mean tpm distribution changes slightly following the addition of CSP, which may represent the altered transcriptional patterns within the cell. However, the general shape of the distributions is similar, suggesting technical biases are unlikely to cause any observed differences between samples.

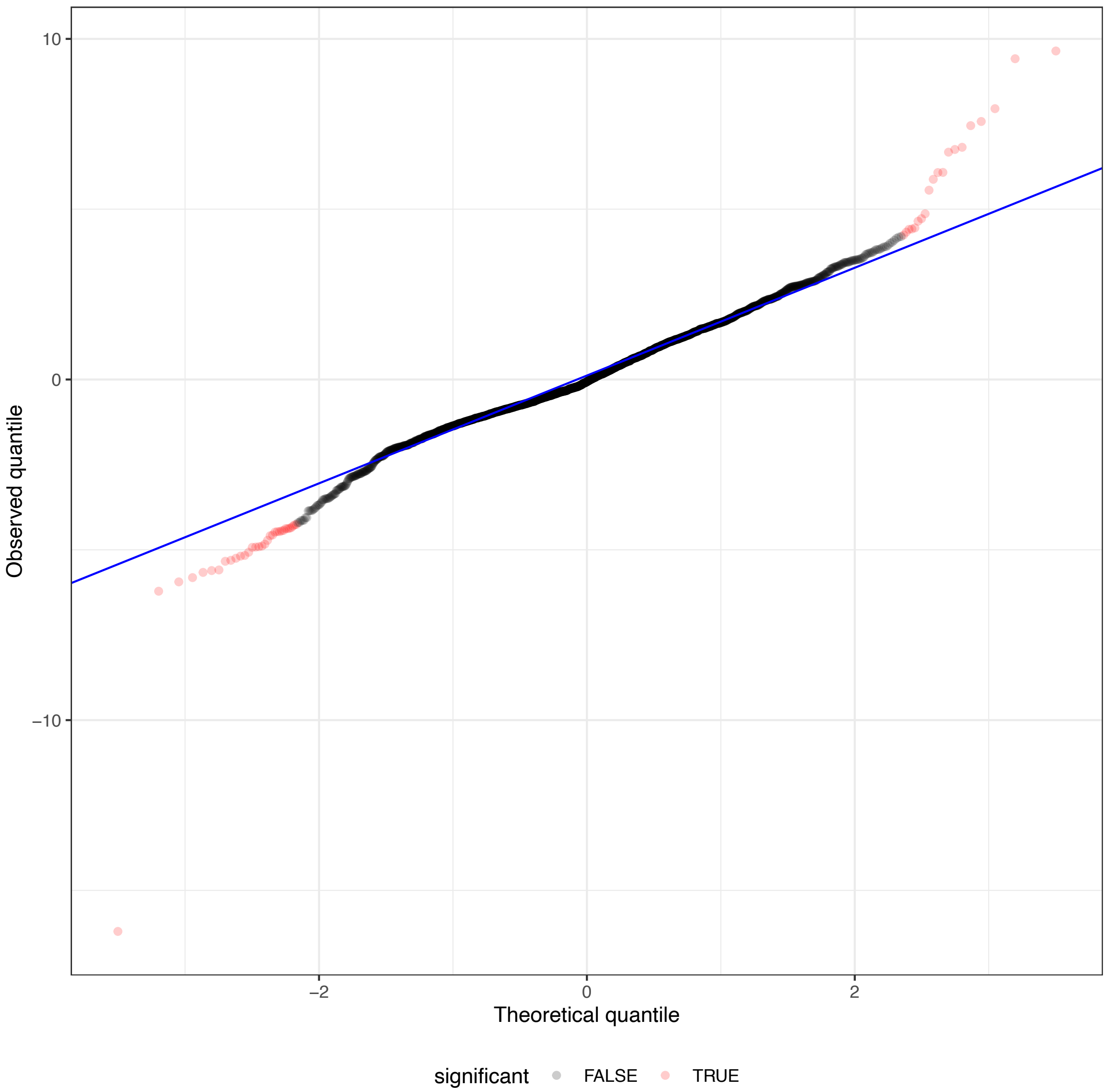

**Figure S8** Q-Q plot comparing the theoretical and observed distributions of the Wald test statistic across genes for the contrast of transcriptional patterns between RMV7 $tvr_{domi}$::Janus and RMV7 $tvr_{rare}$::Janus prior to the addition of CSP. The blue line shows the relationship expected under the null hypothesis of no difference in expression patterns. Each point represents a coding sequence. Points are coloured red if the null hypothesis can be rejected at a false discovery rate of $10^{-3}$, following a Benjamini-Hochberg correction for multiple testing. The Q-Q plot shows this threshold captures the major differences between the two genotypes.

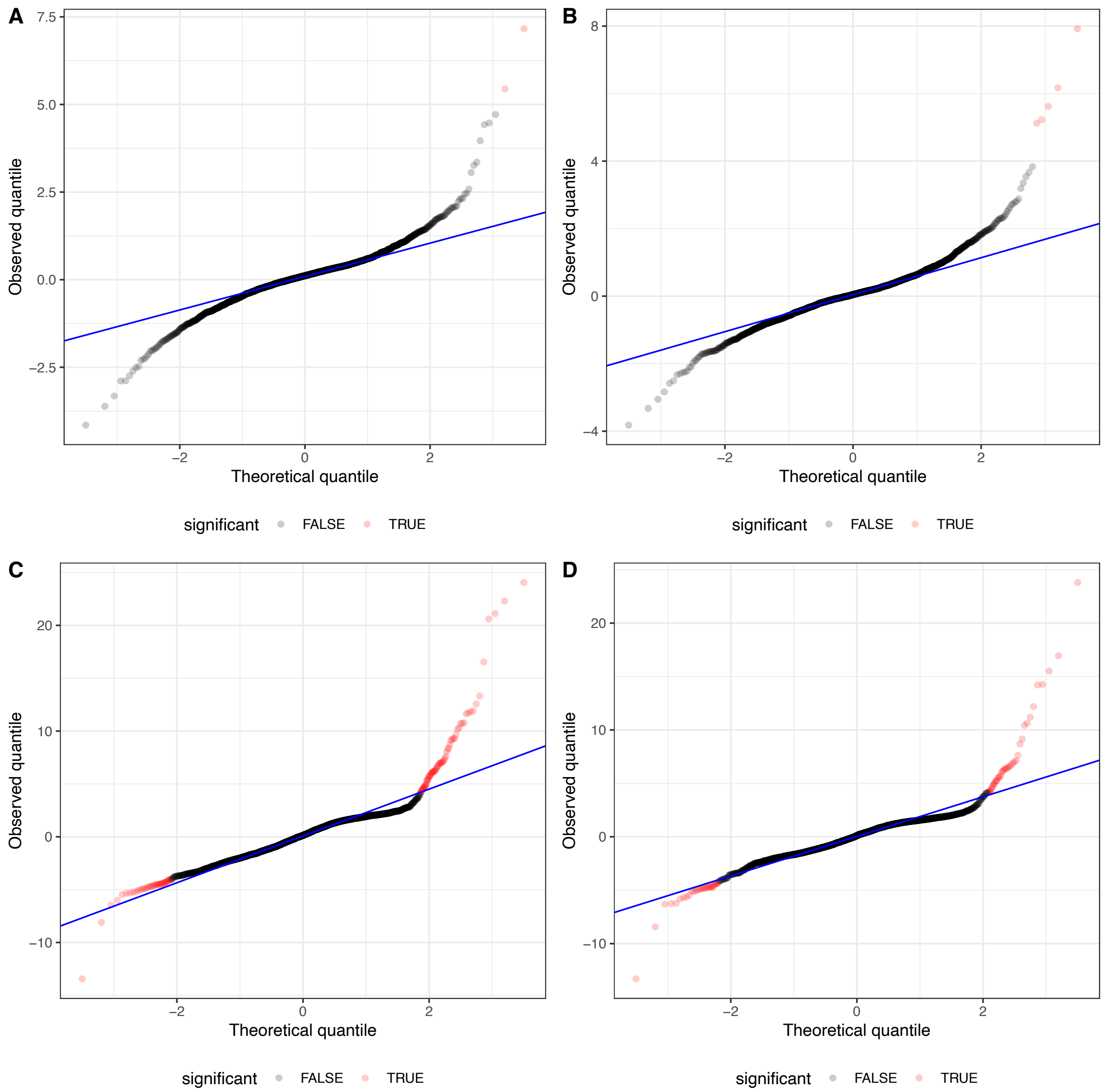

**Figure S9** Q-Q plots comparing the theoretical and observed distributions of the Wald test statistic across genes for the comparison of transcriptional patterns following the addition of CSP. The plots are displayed as in Fig. S8. The panels show contrasts of (A) the 0 minute and 10 minute timepoints for RMV7 $tvr_{domi}$::Janus, and (B) the 0 minute and 20 minute timepoints for RMV7 $tvr_{domi}$::Janus; and (C) the 0 minute and 10 minute timepoints for RMV7 $tvr_{rare}$::Janus, and (D) the 0 minute and 20 minute timepoints for RMV7 $tvr_{rare}$::Janus.

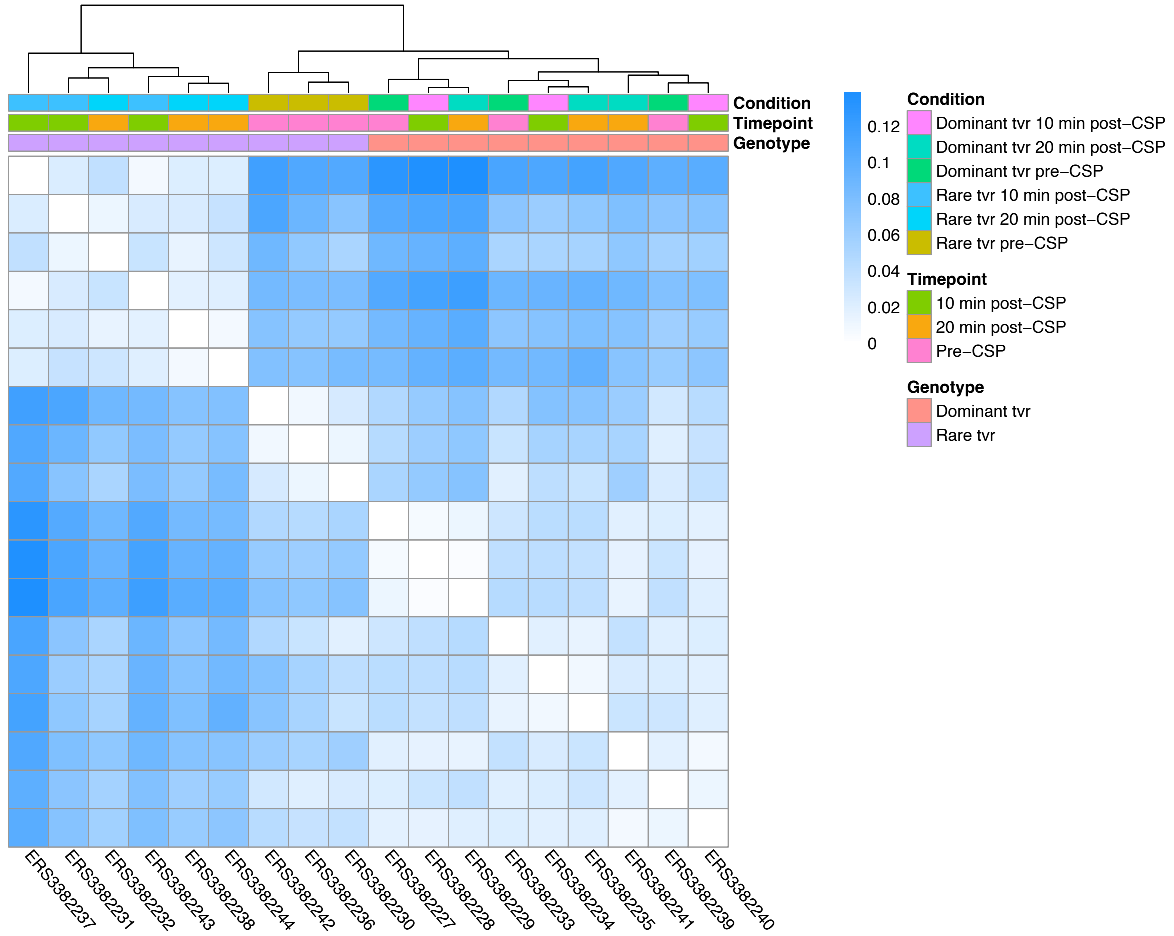

**Figure S10** Clustering of RNA-seq samples by transcriptional patterns. The heatmap represents the dissimilarity between samples. Each column or row corresponds to a sample (Table S2), ordered by their similarity to one another, as illustrated by the dendrogram at the top of the figure. The rows beneath the dendrogram annotate the characteristics of each sample. The fill in each cell shows the pairwise Jensen-Shannon divergence. The matrix is symmetrical, with samples ordered identically across rows and columns, and dissimilarities of zero along the diagonal. These show the samples diverging most strongly from the rest are the post-CSP samples from the RMV7 *tvr*$_{rare}$::Janus genotype.

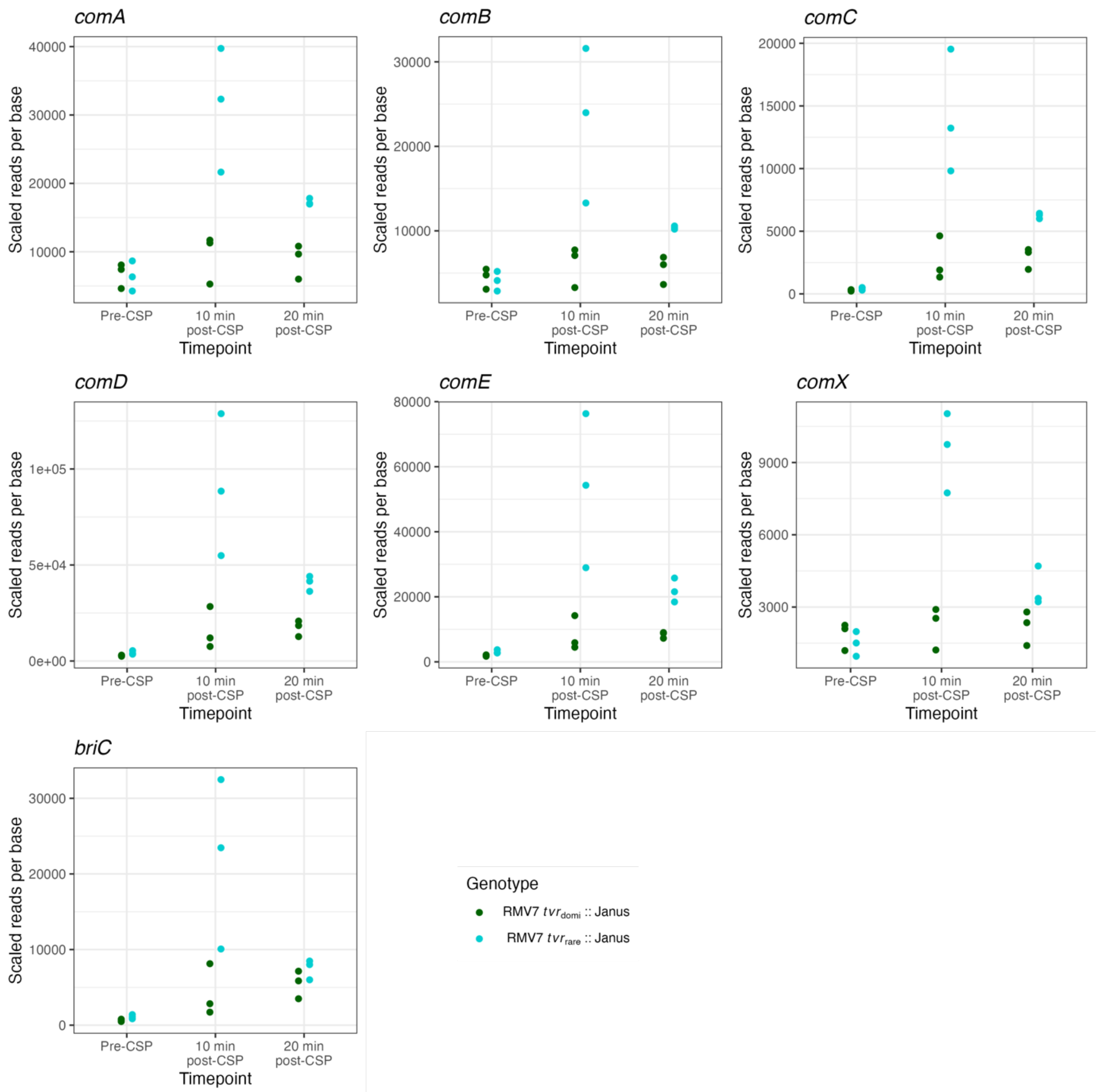

**Figure S11** Quantification of the expression of early competence genes using RNA-seq data. Each plot shows the transcription of a different gene, in scaled reads per base, across the three assayed timepoints in the two variants. There are three biological replicates for each measurement. All these genes show a strong post-CSP upregulation in RMV7$_{rare}$, with a weaker response in RMV7$_{domi}$.

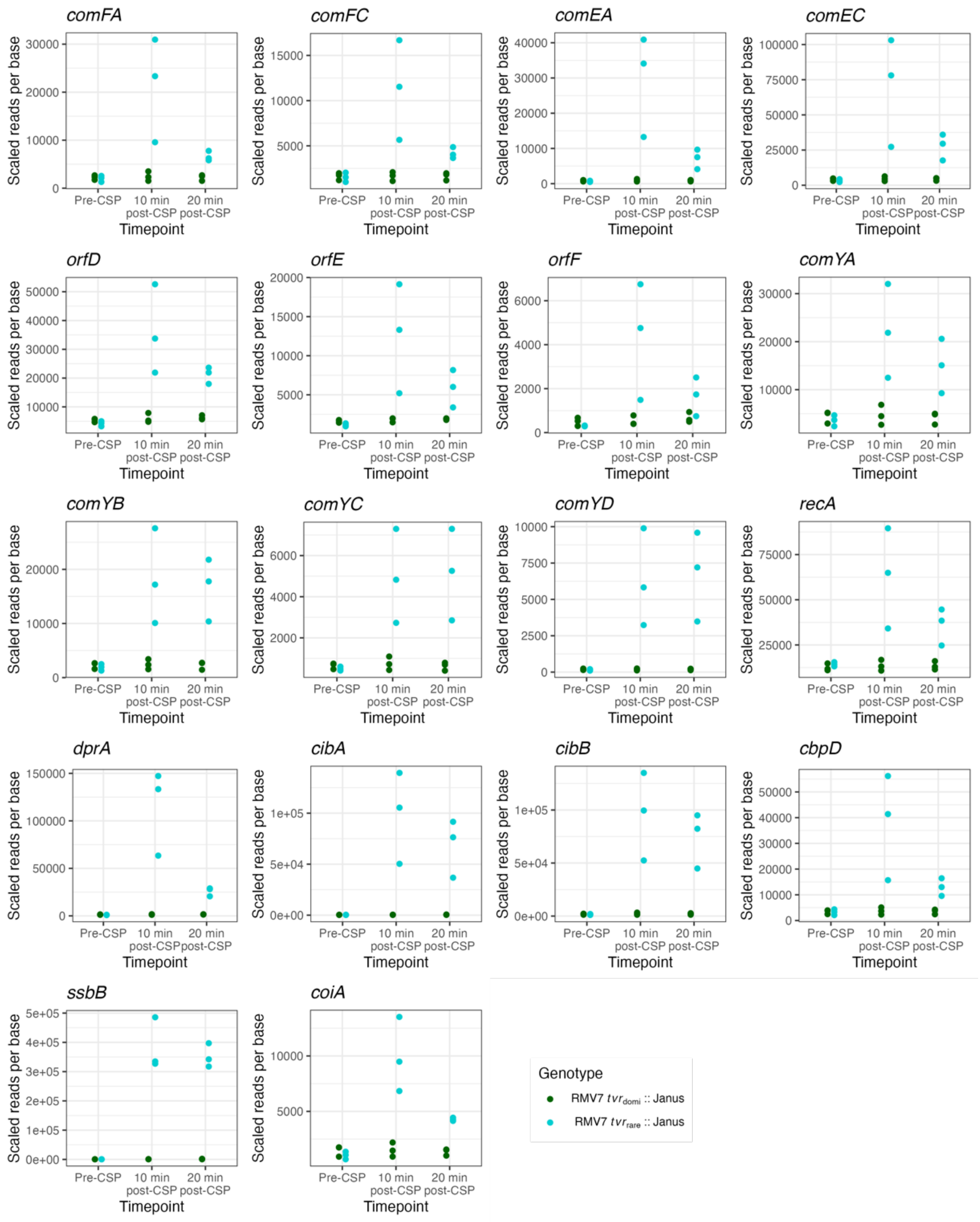

**Figure S12** Quantification of the expression of late competence genes using RNA-seq data. Data are displayed as in Fig. S11. A much stronger induction by CSP is observed in $RMV7_{rare}$ than $RMV7_{domi}$.

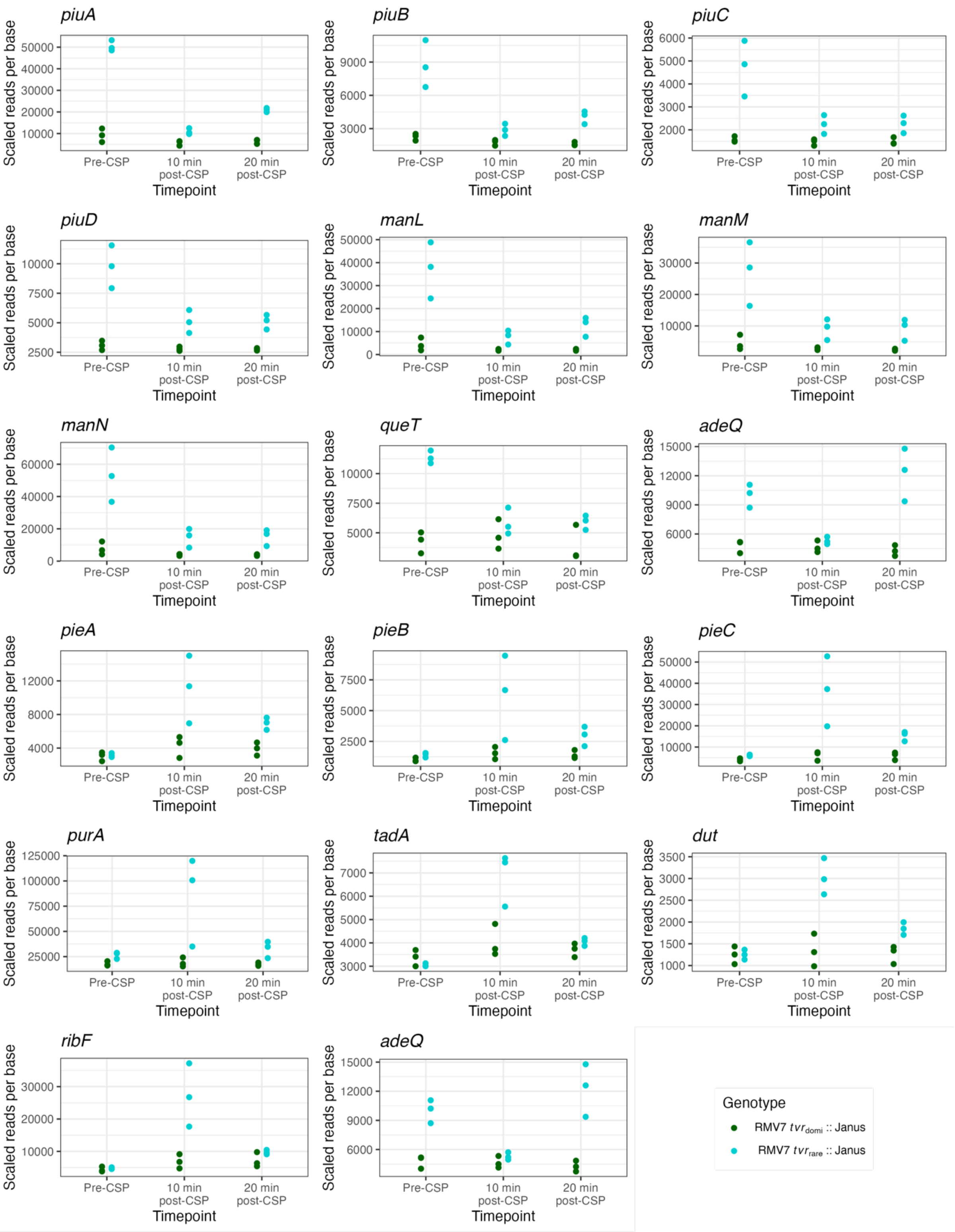

**Figure S13** Quantification of the expression of transporter and nucleotide metabolism genes using RNA-seq data. Data are displayed as in Fig. S11.

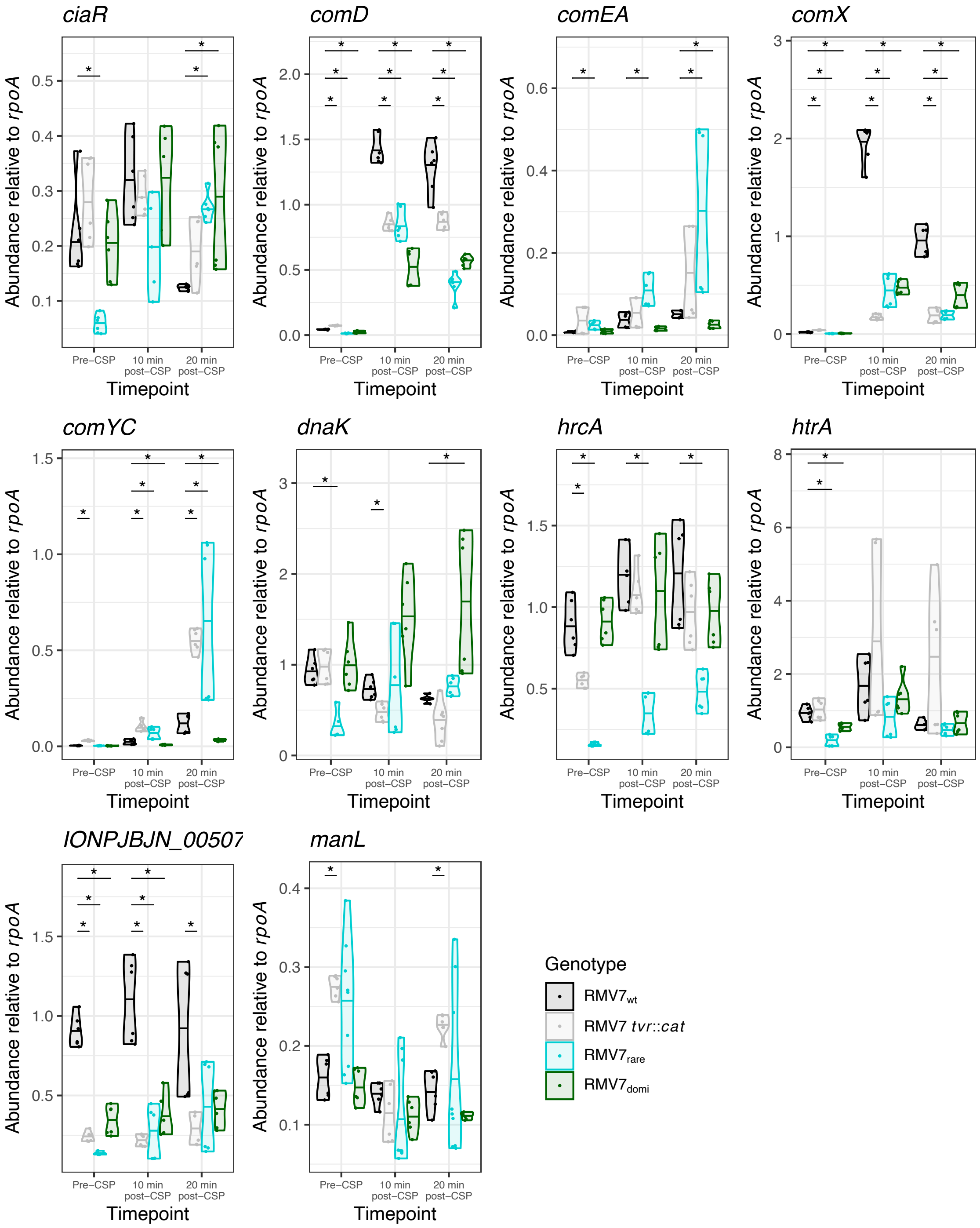

**Figure S14** Quantification of gene expression using qRT-PCR. Samples were collected from RMV7$_{wt}$, RMV7$_{wt}$ *tvr*::*cat*, RMV7$_{domi}$ and RMV7$_{rare}$ prior to the addition of CSP, and both 10 and 20 minutes after the addition of CSP, to mirror the RNA-seq experiment. Each plot corresponds to a specific gene. For each gene, datapoints were combined across three technical replicate measurements of each of two biological replicates. Points are plotted by the timepoint at which they were sampled and coloured by the genotype of the cells from which they were collected. Expression was quantified as the abundance of transcripts relative to *rpoA* RNA. For each gene at each timepoint, Wilcoxon rank sum tests were used to compare each genotype to RMV7$_{wt}$. A Holm-Bonferroni correction for multiple testing was applied within each panel. Significance between results is coded as: $p < 0.05$, *; $p < 0.01$, **; $p < 10^{-3}$, ***; $p < 10^{-4}$, ****.

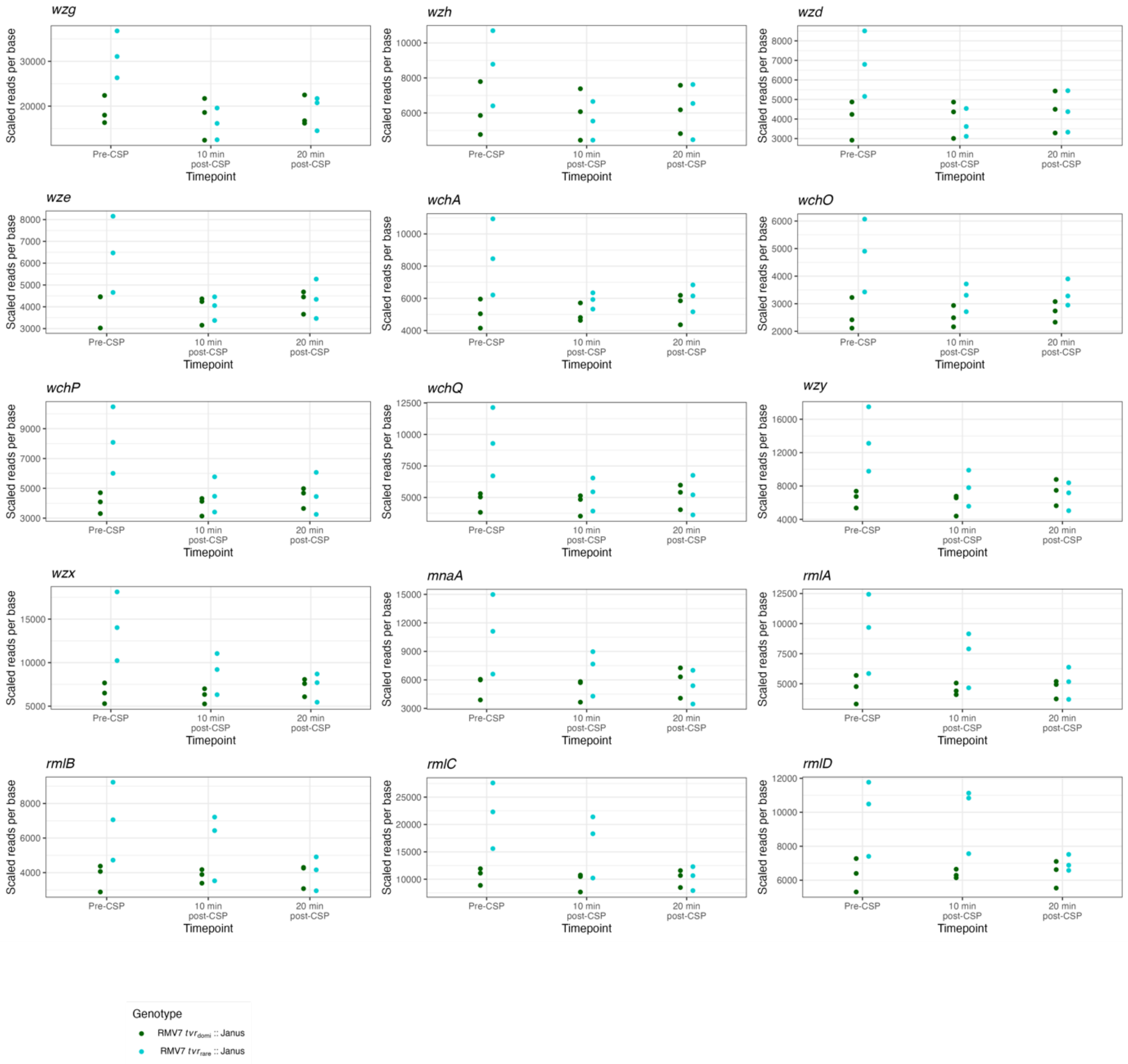

**Figure S15** Quantification of the expression of *cps* locus genes using RNA-seq data. Data are displayed as in Fig. S11.

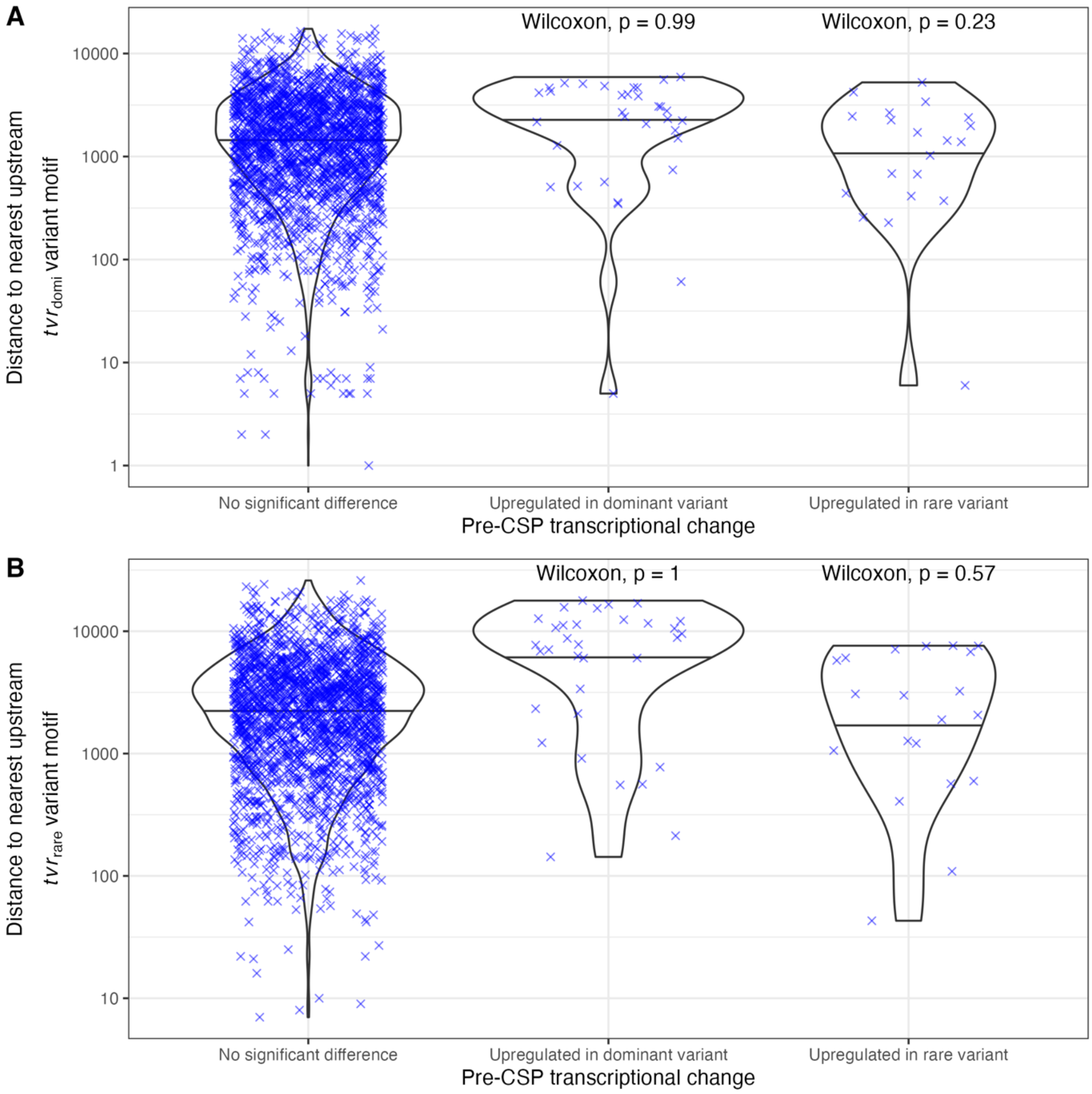

**Figure S16** Violin plots showing the relationship between *Spn*IV methylation sites and transcriptional variation. Each point represents the distance between a protein coding sequence's translational start site, and the nearest upstream *Spn*IV methylation site based on (A) $tvr_{domi}$ motifs and (B) $tvr_{rare}$ motifs. Coding sequences (CDSs) were classified based on whether they exhibited a significant difference in transcription between RMV7 $tvr_{domi}$::Janus and RMV7 $tvr_{rare}$::Janus prior to the addition of CSP. A one-tailed Wilcoxon rank sum test was used to test the hypothesis that CDSs that differed in expression significantly between the two phase variants pre-CSP would be closer to upstream variable methylation sites than the equivalent distances for CDSs that were not associated with strong changes in transcription. No evidence was found of differential expression between the variants being associated with greater proximity of *Spn*IV methylation sites to a coding sequence's start codon.

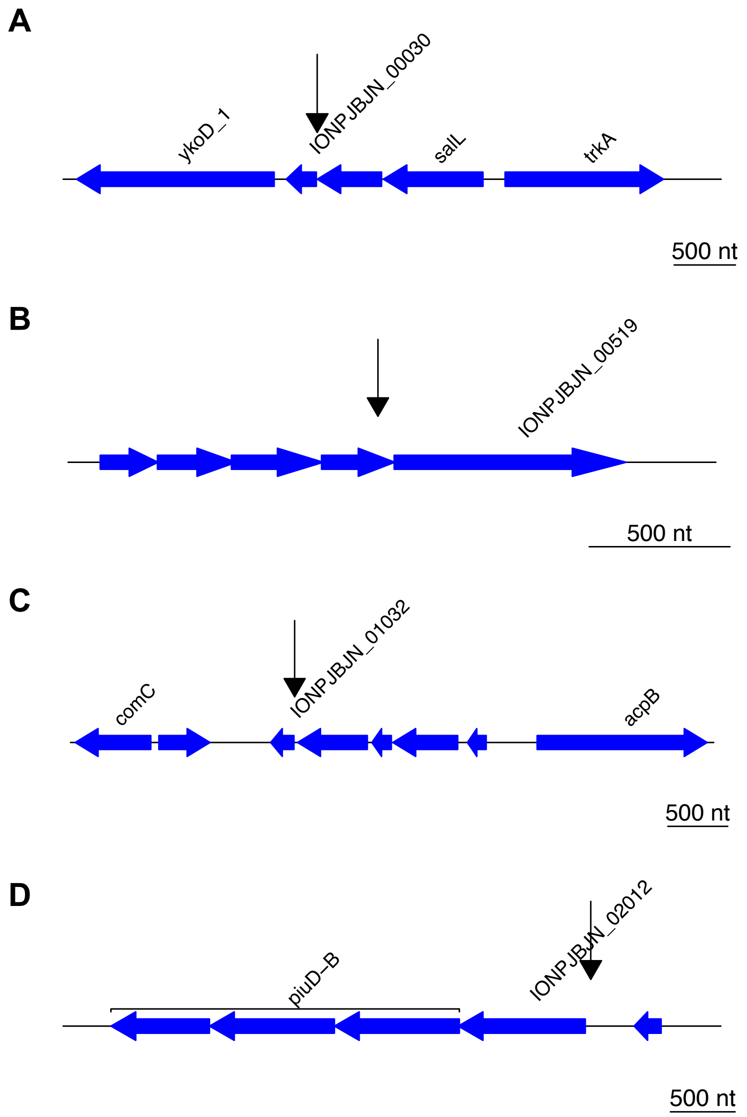

**Figure S17** Spatial relationship between methylation sites and differentially expressed genes. Four genes that significantly differed in levels of transcription between the pre-CSP RMV7 *tvr*$_{\text{domi}}$::Janus and RMV7 *tvr*$_{\text{rare}}$::Janus samples had *Spn*IV methylation sites within 100 nt of their start codon (Fig. S16). In three cases, this corresponded to *tvr*$_{\text{domi}}$ methylation sites shortly upstream of genes within operons: (A) IONPJBJN_00030, which is separated from the upstream gene by only six bases; (B) IONPJBJN_00519, which is within PRCI$_{dnaN}$; and (C) IONPJBJN_01032, which is the final gene (*csbD*) in an operon that appears to be part of the MgrA regulon (Fig. S21). (D) A single gene, *piuA* (IONPJBJN_02012), is the first in an operon, and has a *tvr*$_{\text{rare}}$ motif within its putative promoter sequence. This operon is upregulated in RMV7$_{\text{rare}}$, and this may represent a direct link between methylation and changes in gene expression.

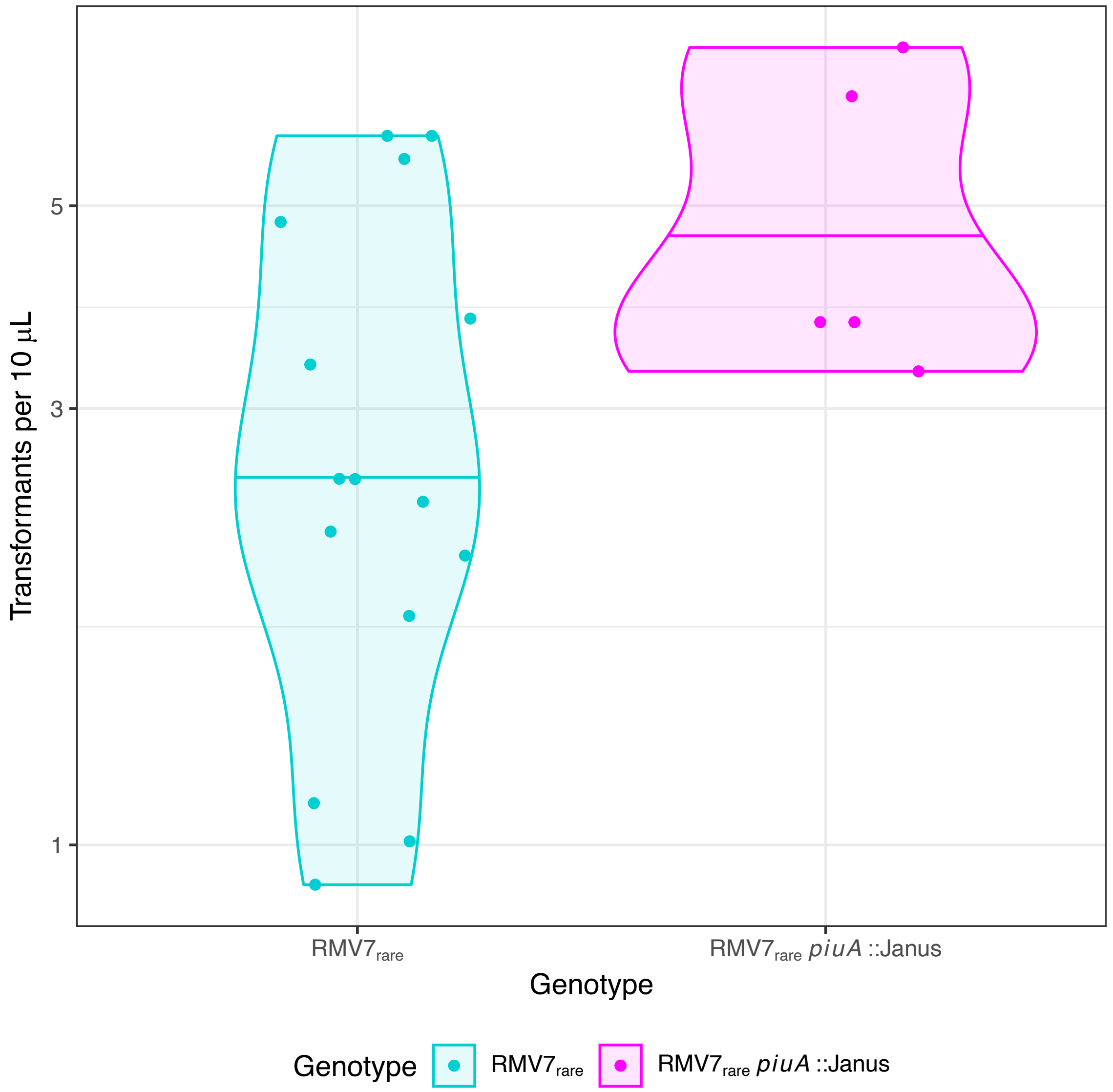

**Figure S18** Violin plot comparing the transformation efficiency of RMV$_{rare}$ and a mutant in which *piuA* had been disrupted by a Janus cassette. Each individual point represents an independent transformation experiment. The horizontal line within the violins shows the median for each genotype. The disruption of *piuA* did not detectably reduce the transformability of RMV7$_{rare}$, suggesting the methylation site within its promoter and the associated upregulation in RMV7$_{rare}$ did not cause the observed phenotypic differences between the variants.

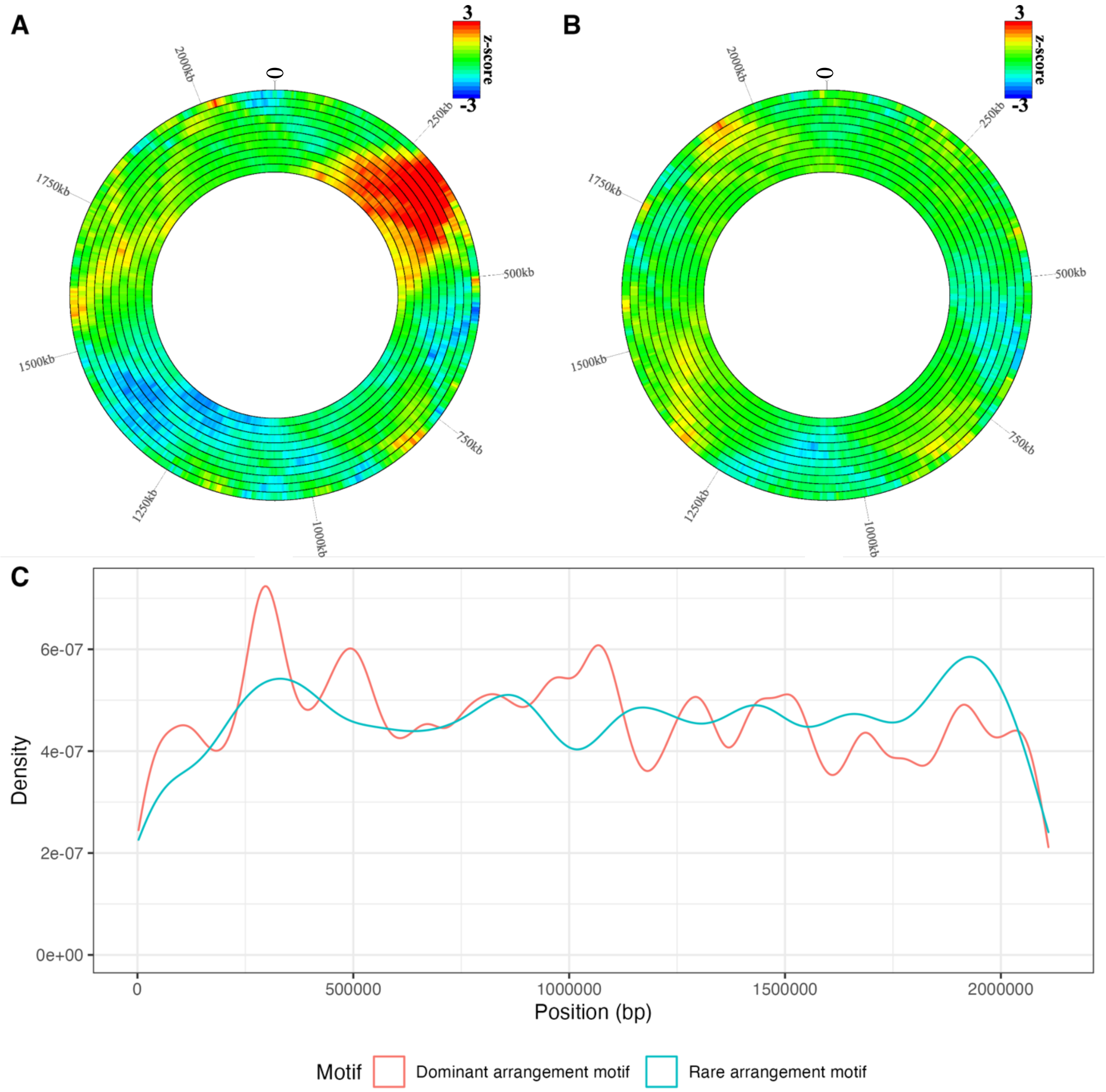

**Figure S19** Distribution of methylation motifs across the chromosome. The distribution of the (A) $\textit{tvr}_{\mathrm{domi}}$ (TGAN$_7$TCC) and (B) $\textit{tvr}_{\mathrm{rare}}$ (TGAN$_7$TATC) motifs across both strands of the RMV7$_{\mathrm{domi}}$ genome were analysed with DistAMo. The ten rings represent different sizes of sliding windows, increasing from 50 kb in the outermost ring, to 500 kb in the innermost ring, in 50 kb increments. The density of motifs is quantified as a z-score, as indicated by the key. These values are shown as a heatmap across the chromosome of RMV$_{\mathrm{domi}}$. The red regions correspond to a local over-representation of motifs, whereas the blue regions correspond to a local under-representation. (C) Line graph comparing the density of $\textit{tvr}_{\mathrm{domi}}$ and $\textit{tvr}_{\mathrm{rare}}$ motifs across the RMV7$_{\mathrm{domi}}$ genome.

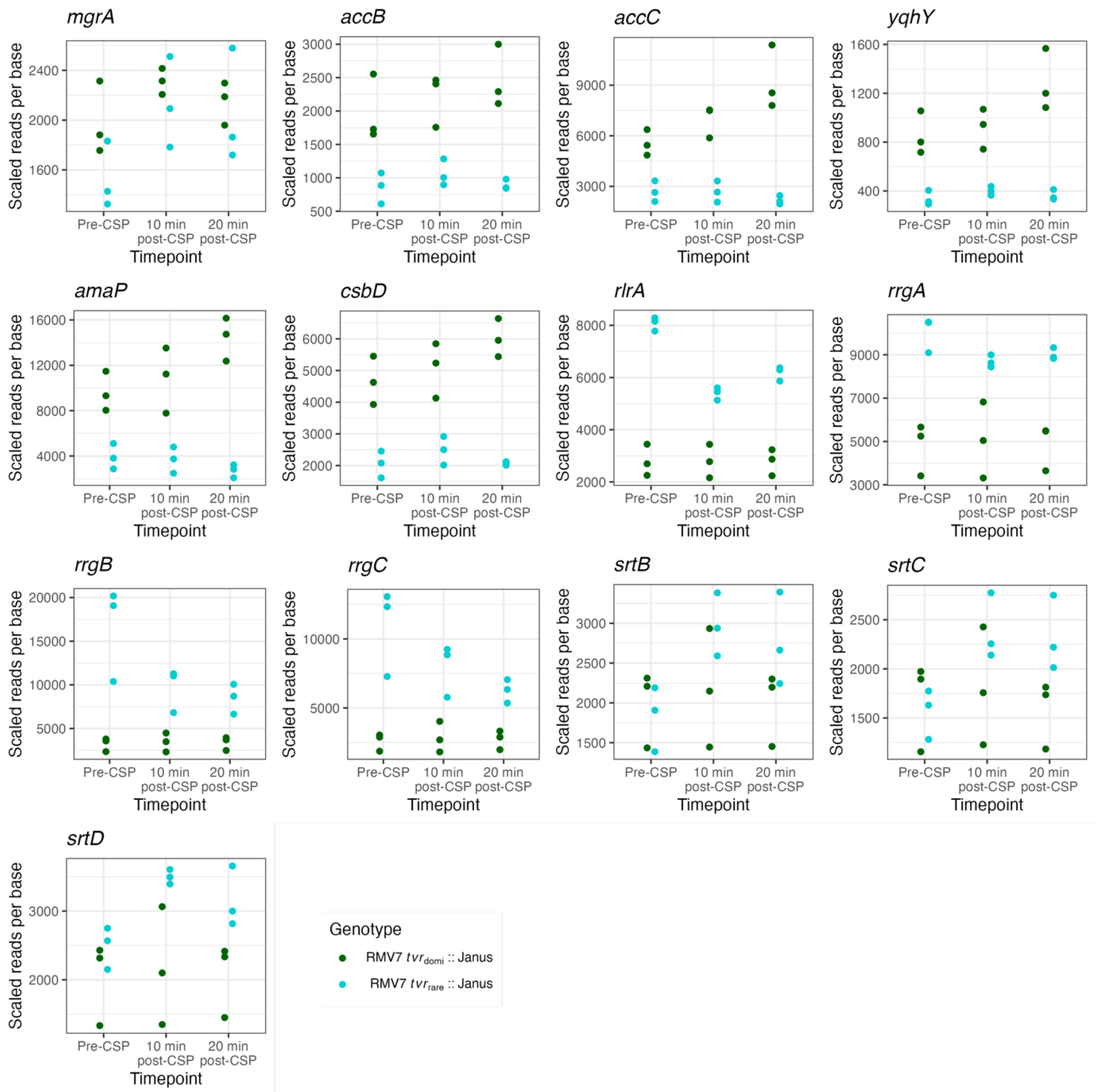

**Figure S20** Quantification of the expression of *mgrA* regulon genes using RNA-seq data. Data are displayed as in Fig. S11.

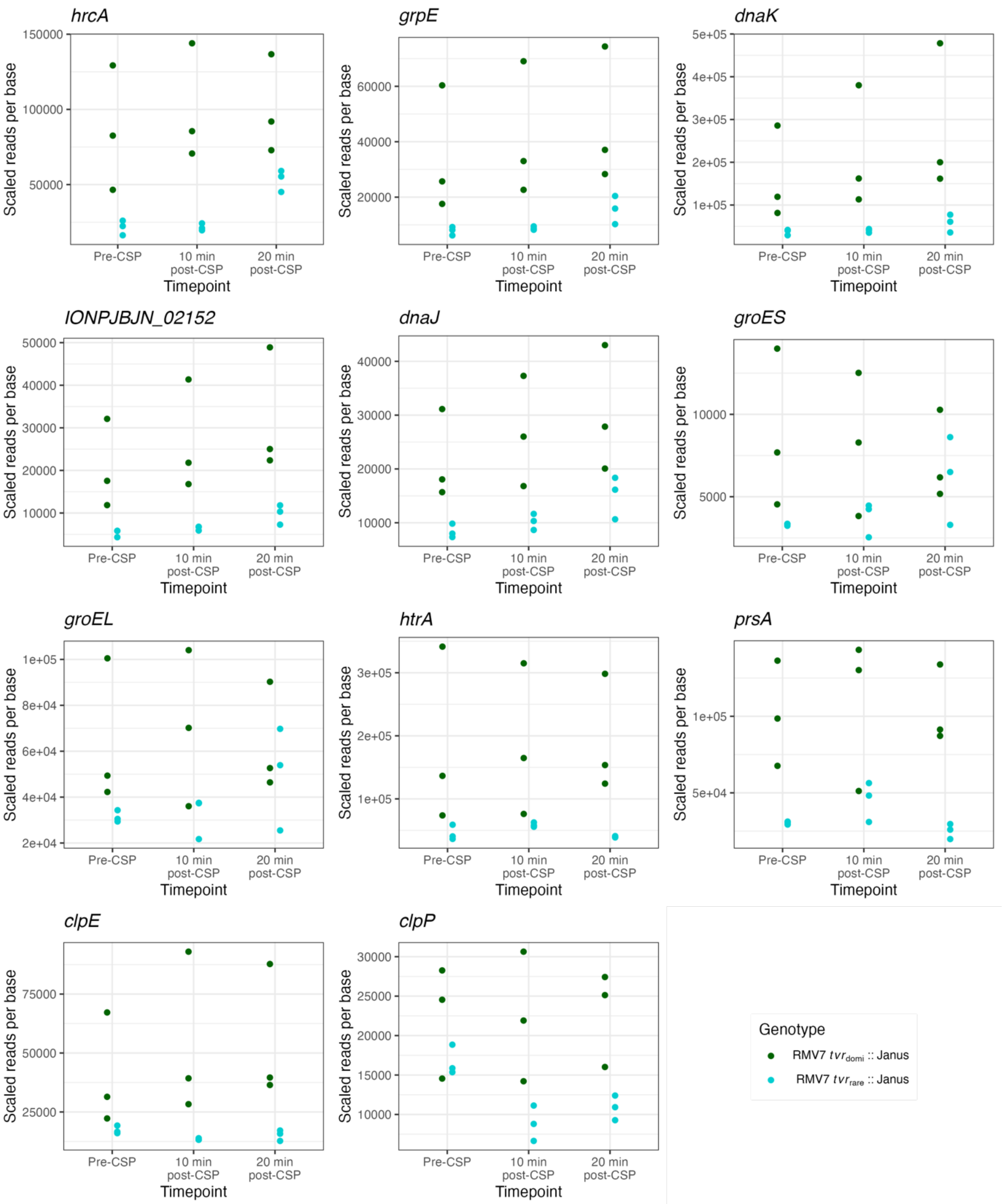

**Figure S21** Quantification of the expression of stress response and chaperone genes using RNA-seq data. Data are displayed as in Fig. S11.

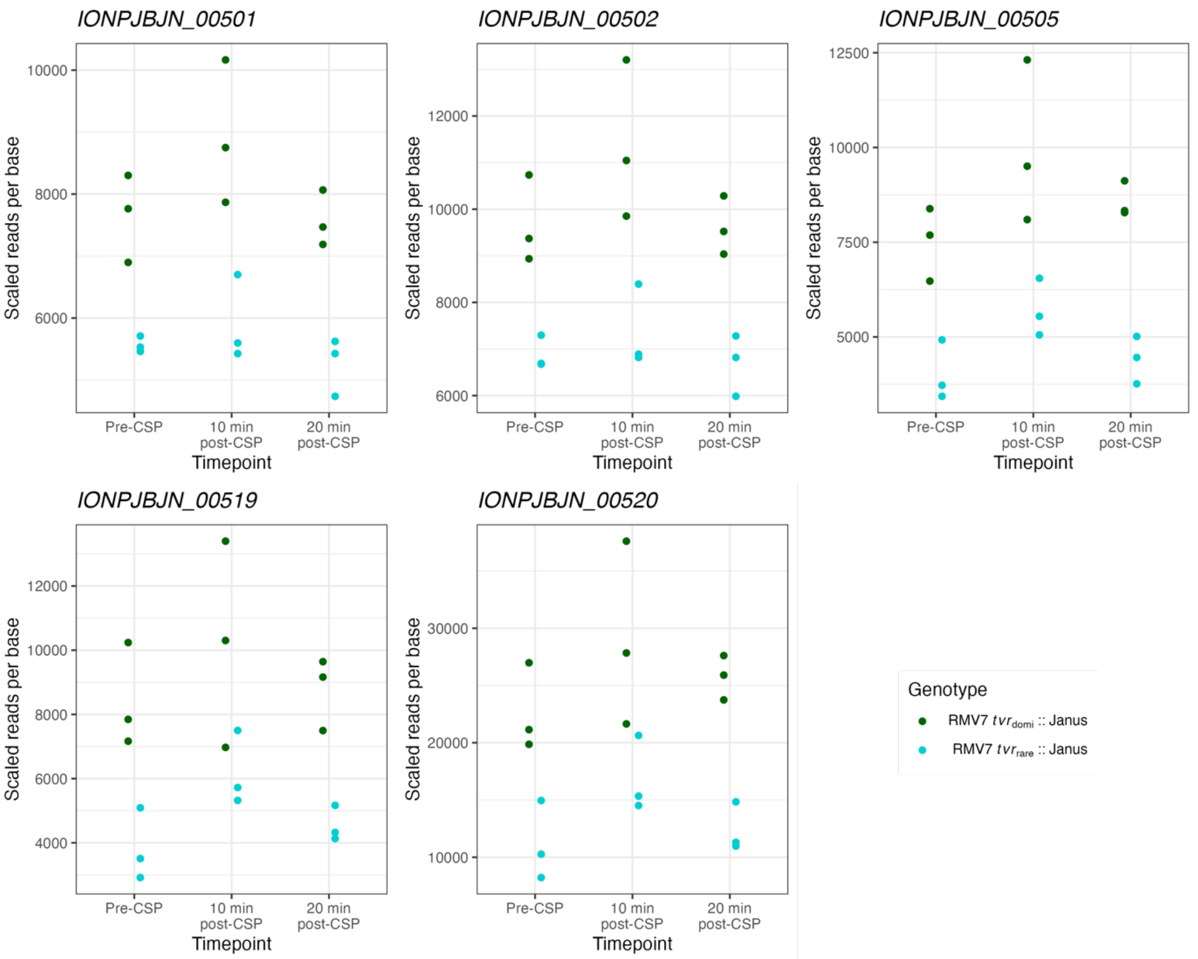

**Figure S22** Quantification of the expression of large PRCI$_{dnaN}$ genes using RNA-seq data. These were all more active in RMV7$_{domi}$. Data are displayed as in Fig. S11.

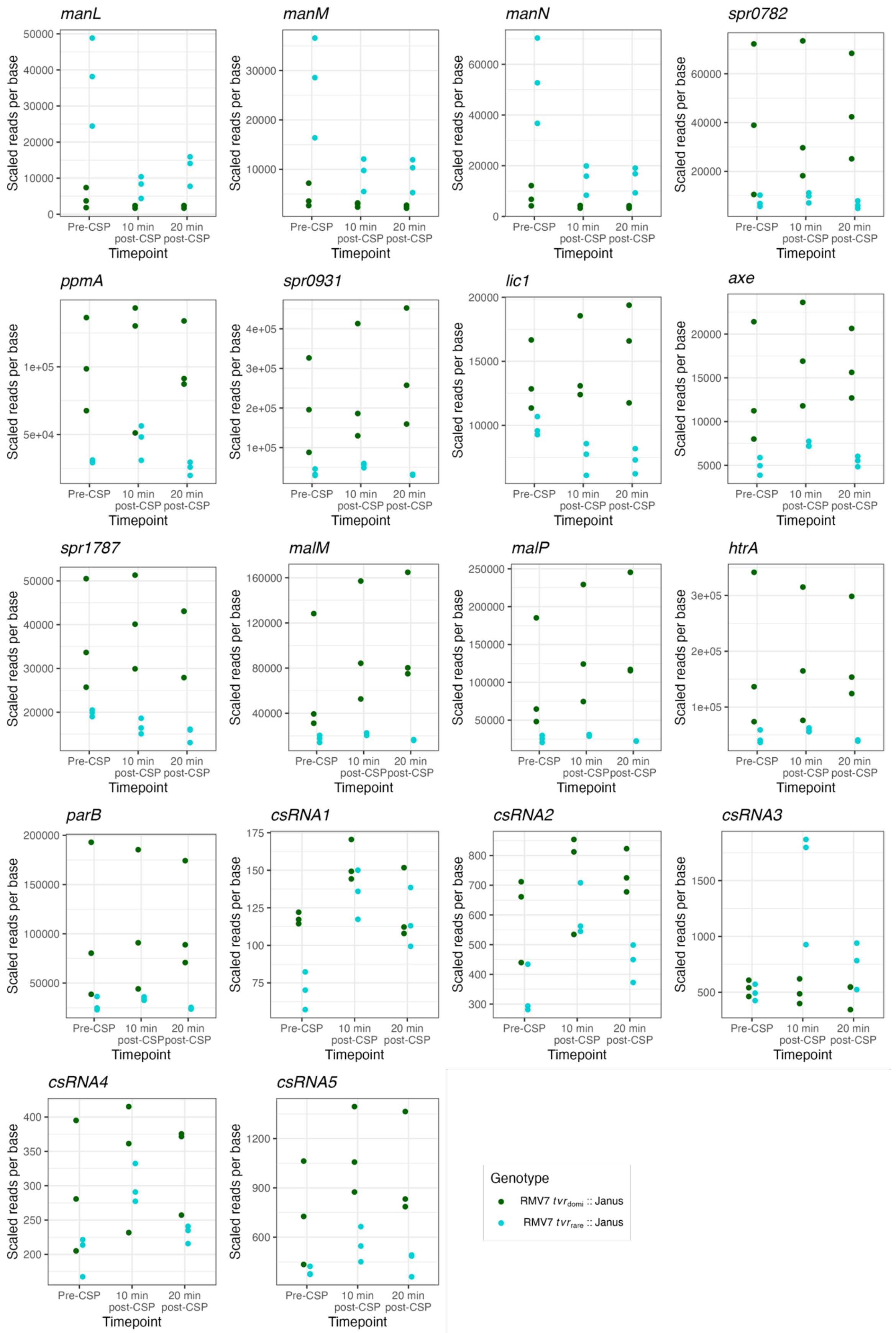

**Figure S23** Quantification of the expression of *ciaRH* regulon genes using RNA-seq data. Data are displayed as in Fig. S11.

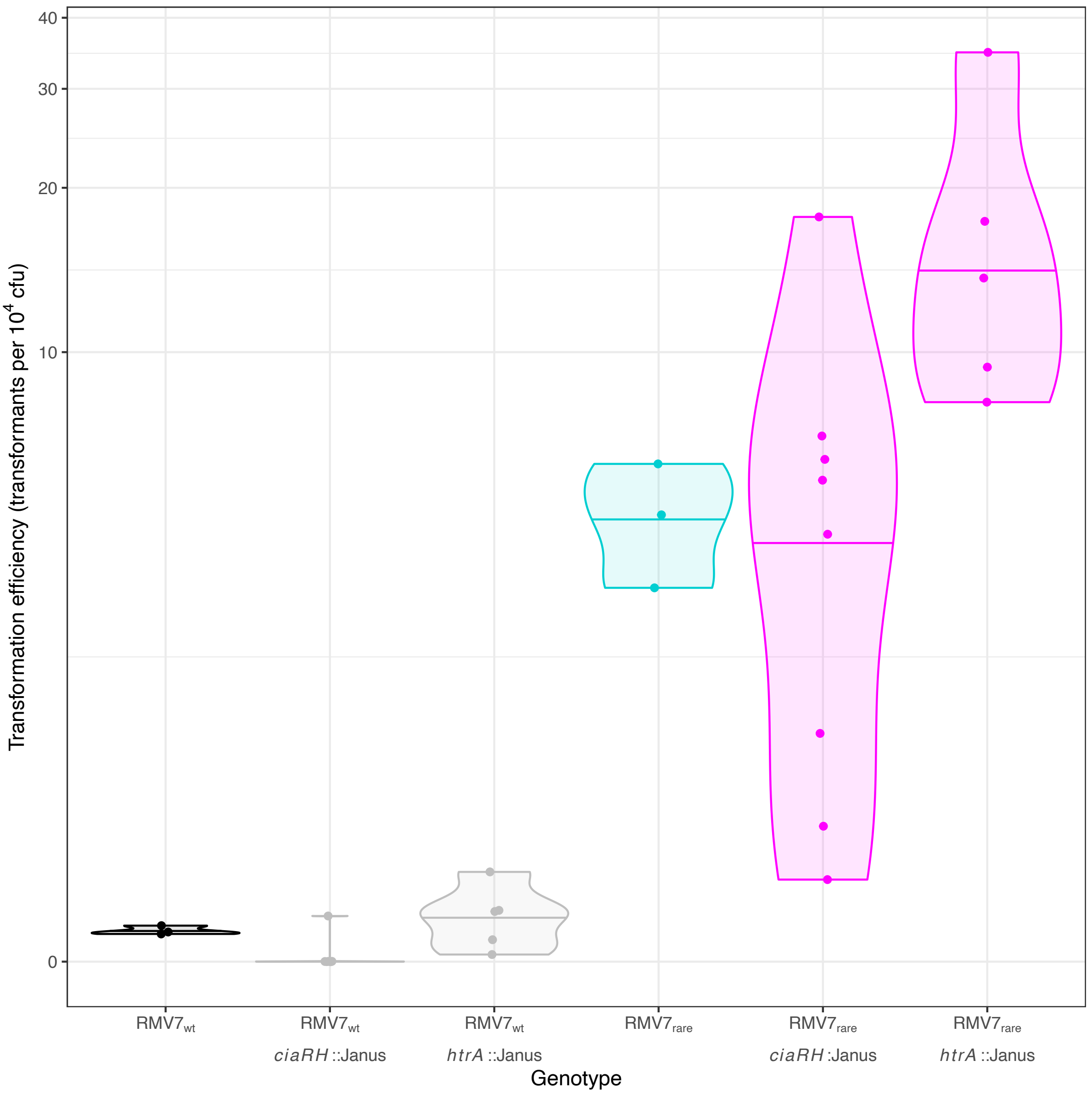

**Figure S24** Violin plot comparing the transformation efficiency of mutants in which *ciaRH* or *htrA* was disrupted in the RMV7$_{wt}$ and RMV7$_{rare}$ backgrounds. Each individual point represents an independent transformation experiment. The horizontal line within the violins shows the median for each genotype. Disruption of *ciaRH* had limited impact on the transformation efficiency of either genotype. However, disruption of *htrA* increased the transformation efficiency of the already highly transformable RMV7$_{rare}$.

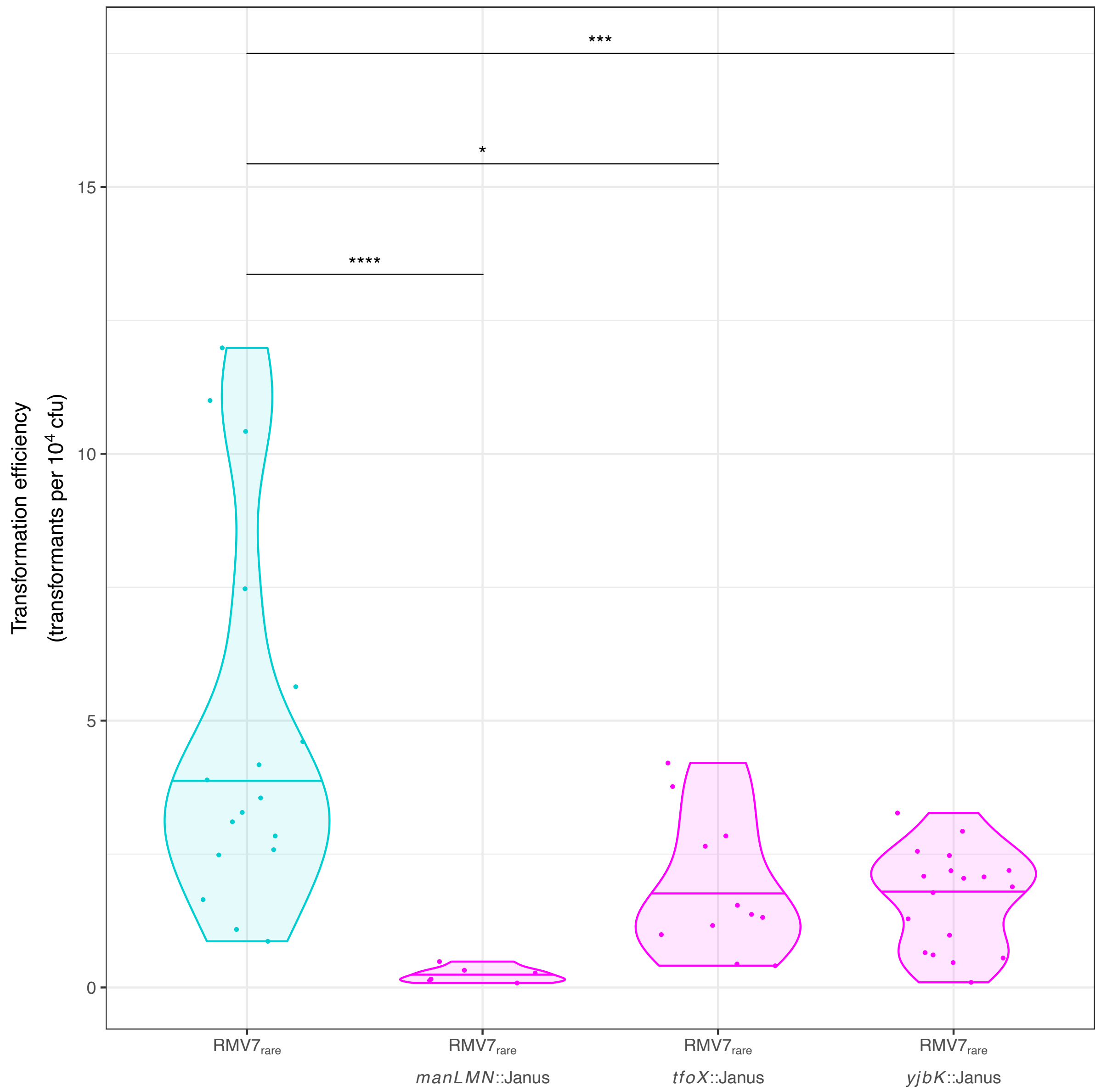

**Figure S25** Violin plot comparing the transformation efficiencies of RMV7$_{rare}$ mutants in *manLMN*, *tfoX* and *yjbK* with the parental genotype. The data shown are the same as displayed for the experiments conducted in unsupplemented media in Fig. 3A, but here the two-tailed Wilcoxon rank sum test compares the parental genotype with the mutants. Significance between results is coded as: $p < 0.05$, *; $p < 0.01$, **; $p < 10^{-3}$, ***; $p < 10^{-4}$, ****.

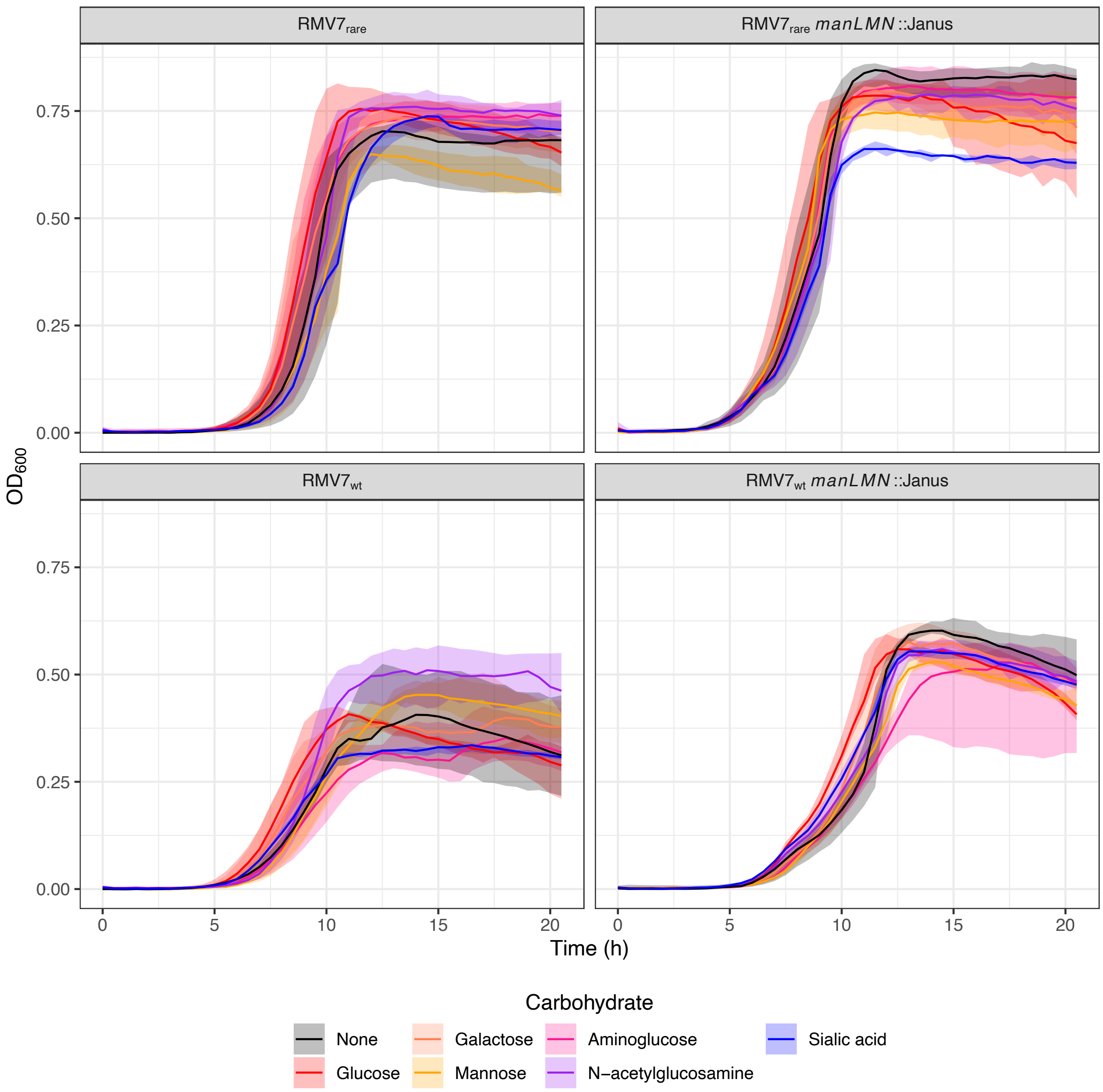

**Figure S26** Effects of ManLMN and carbon source on *in vitro* growth of $RMV7_{wt}$ and $RMV7_{rare}$. Both $RMV7_{wt}$ and $RMV7_{rare}$, and mutants of each lacking *manLMN*, were grown in unsupplemented mixed media liquid (a nutrient-rich mix of Todd-Hewitt media, with 0.5% yeast extract, and Brain-Heart Infusion). This was either unsupplemented, or supplemented with one of six additional carbon sources, indicated by the colour of the growth curve. Growth was measured using the optical density at 600 nm ($OD_{600}$) across three biological replicates. The lines represent the median optical density, and the shaded ribbon shows the range of the replicates. There was little evidence of the loss of ManLMN causing a substantial decrease in growth. The only carbon source found to increase the final population size was *N*-acetylglucosamine in $RMV7_{wt}$, which was not observed when *manLMN* was disrupted in this background

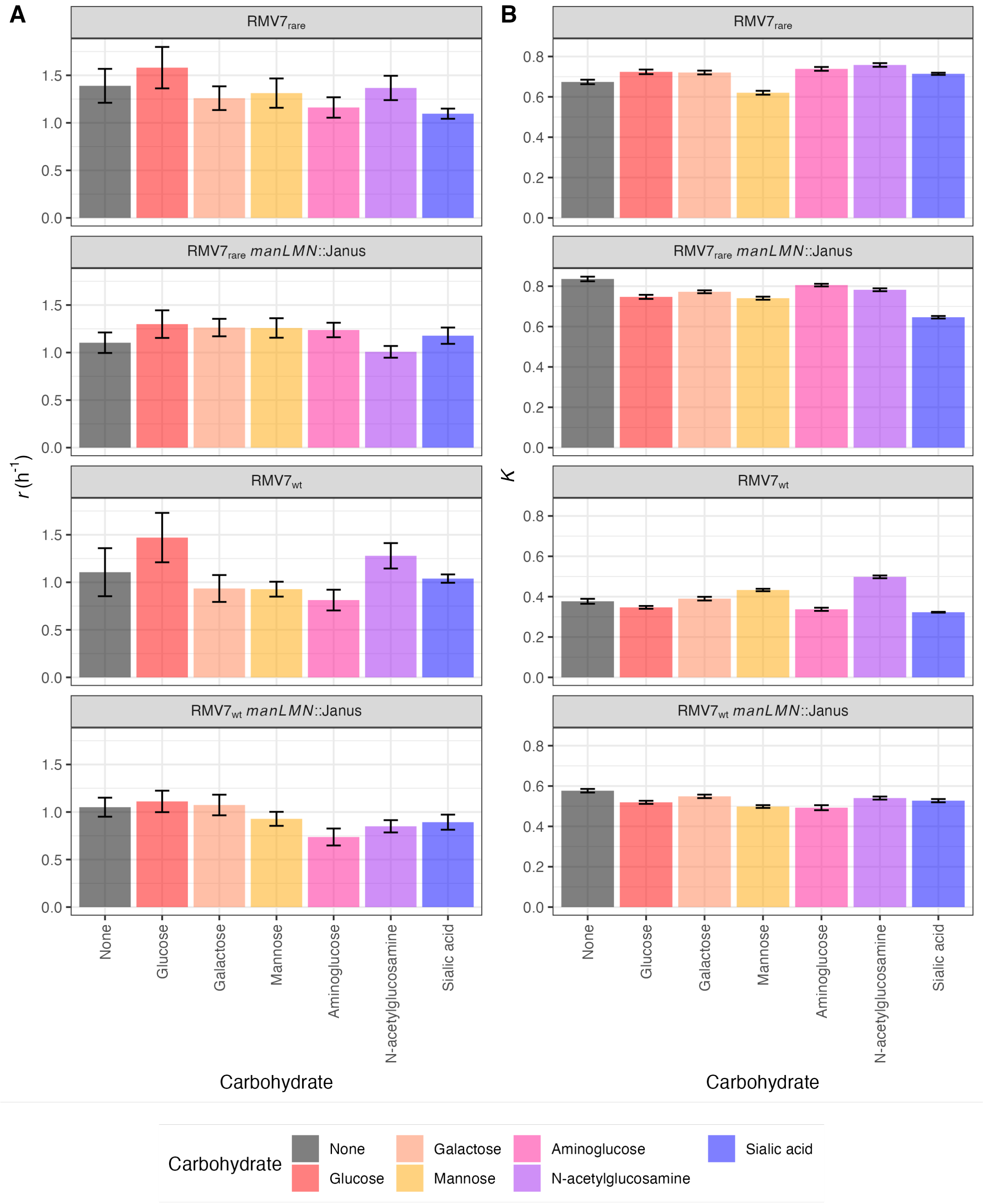

**Figure S27** Barplots summarising the effects of ManLMN and carbon source supplements on growth of $RMV7_{domi}$ and $RMV7_{rare}$. The plots are separated by variant and whether *manLMN* was disrupted by a Janus cassette insertion. The colours of the bars show the carbon source supplement in the media. The height of the bars show the estimates of logistic model parameters, and the error bars show the 95% confidence intervals of these estimates. (A) Estimates of the reproduction rate, *r*. (B) Estimates of the carrying capacity, *K*.

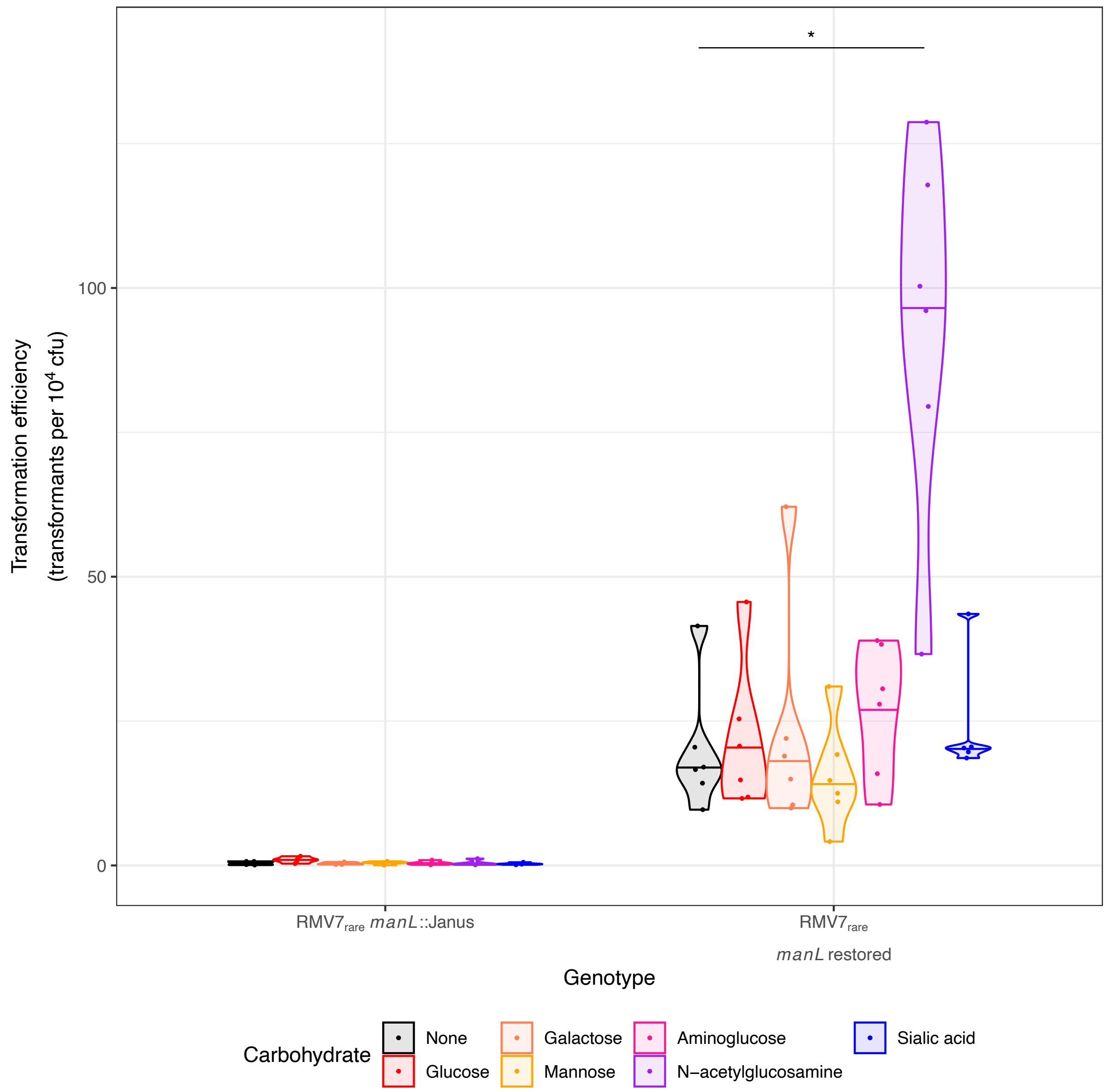

**Figure S28** Confirming the role of *manL* in regulating the competence system. The large size of the *manLMN* operon, combined with the low transformation efficiency of the *manLMN*::Janus genotype, made it impractical to restore the entire operon. Hence a second mutant was constructed in which only *manL* was disrupted with a Janus cassette. This smaller mutation could then be reversed in the more transformable $RMV7_{rare}$ cells. The data are shown as in Fig. 3A. This shows disruption of *manL* reduced the transformation efficiency of $RMV7_{rare}$ to a similar extent as disruption of the entire *manLMN* operon. Restoration of the *manL* gene resulted in a more transformable genotype that responded to GlcNAc, confirming this locus was responsible for the decrease in transformation efficiency, and lack of response to GlcNAc, in the *manL*::Janus mutant.

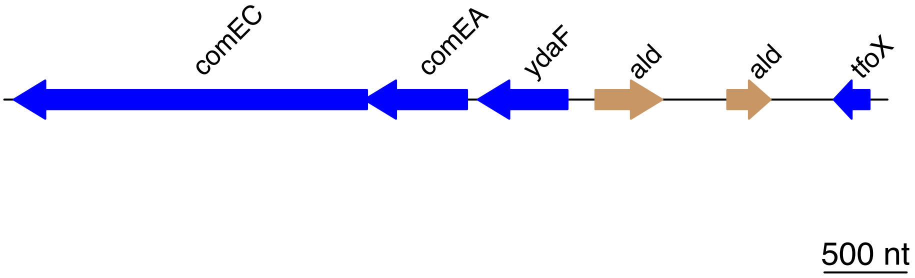

**Figure S29** Location of the *tfoX* gene in the *S. pneumoniae* chromosome. Blue arrows represent intact genes, and brown arrows represent gene fragments. The *tfoX* gene in RMV7 is found at a conserved position, shortly upstream of the *comEC* and *comEA* genes. These genes encode the main DNA entry pore of the competence machinery. The intervening gene fragments are part of a pseudogene predicted to have encoded an alanine dehydrogenase.

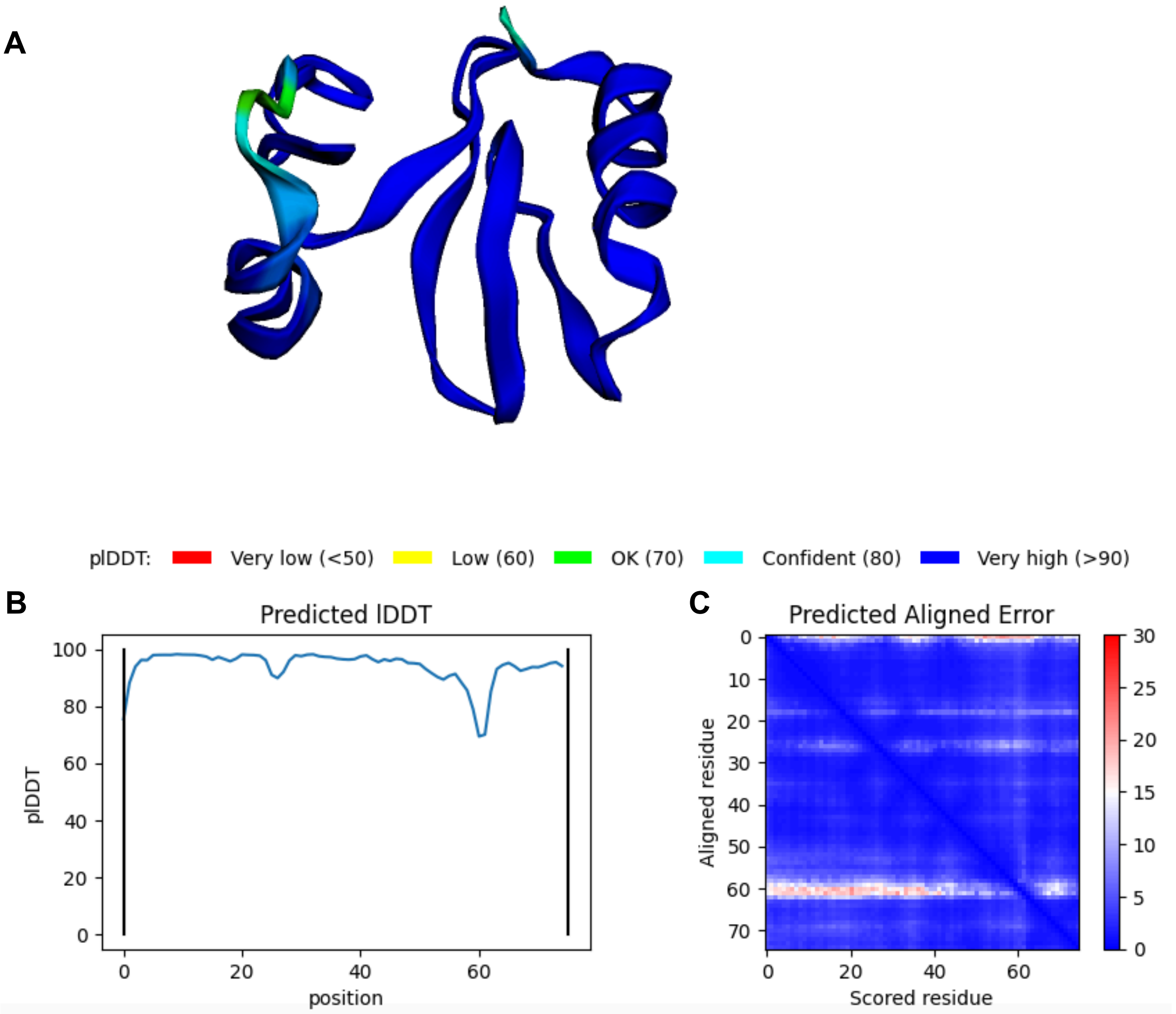

**Figure S30** Predicted structure of TfoX$_{Spn}$. (A) The amino acid sequence of RMV7$_{domi}$ *tfoX* (IONPJBJN_02097) was analysed using the AlphaFold2 pipeline with default settings. The protein structure prediction at the top shows a four-strand beta sheet flanked by alpha helices. The structure is coloured according to the confidence of the prediction, as quantified by the per-residue local distance difference test (plDDT). (B) A line graph of plDDT over the length of the protein. This shows the algorithm has high confidence in the predicted structure. (C) A heatmap showing the uncertainty from the alignment. This shows there to be few regions of uncertainty in the prediction.

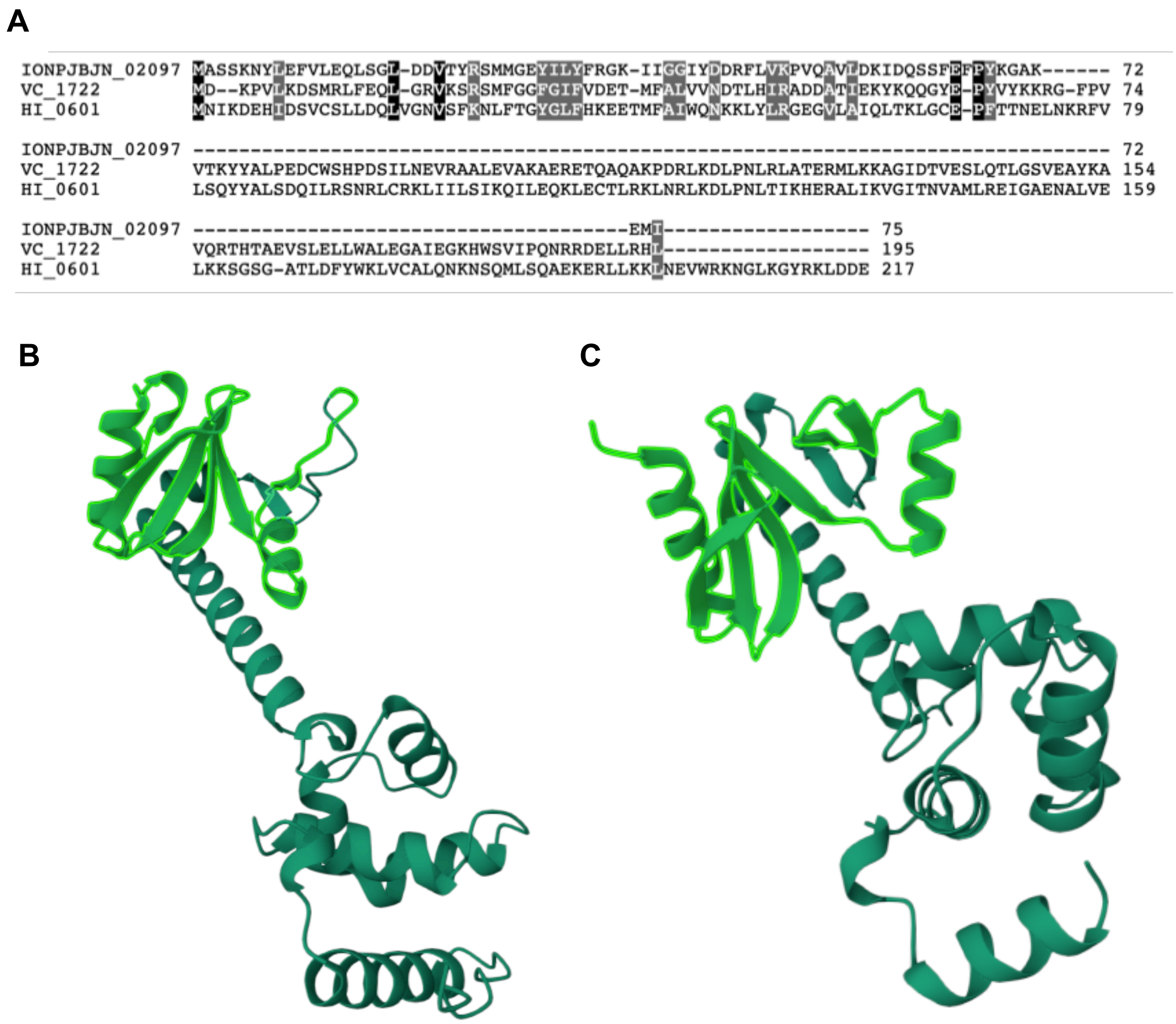

**Figure S31** (A) Alignment of the TfoX proteins of *Vibrio cholerae* (VC_1772 from *V. cholerae* ATCC 39315) and *Haemophilus influenzae* (HI_0601 from *H. influenzae* Rd) with the orthologue in RMV7, IONPJBJN_02097. Columns are coloured black where an amino acid is conserved at a position, and grey when amino acids are similar at a position. (B) The predicted structure of HI_0601, inferred using AlphaFold (AlphaFoldDB identifier AF-P43779-F1). The first 72 amino acids are highlighted in light green. These have a similar structure to that predicted for TfoX$_{Spn}$ (Fig. S30): two alpha helices linked by a four-strand beta sheet. (C) The predicted structure of VC_1772, inferred using AlphaFold (AlphaFoldDB identifier AF-Q9KRC0-F1). The first 72 amino acids are highlighted in light green. These have a similar structure to that predicted for TfoX$_{Spn}$ and the N terminal region of HI_0601.

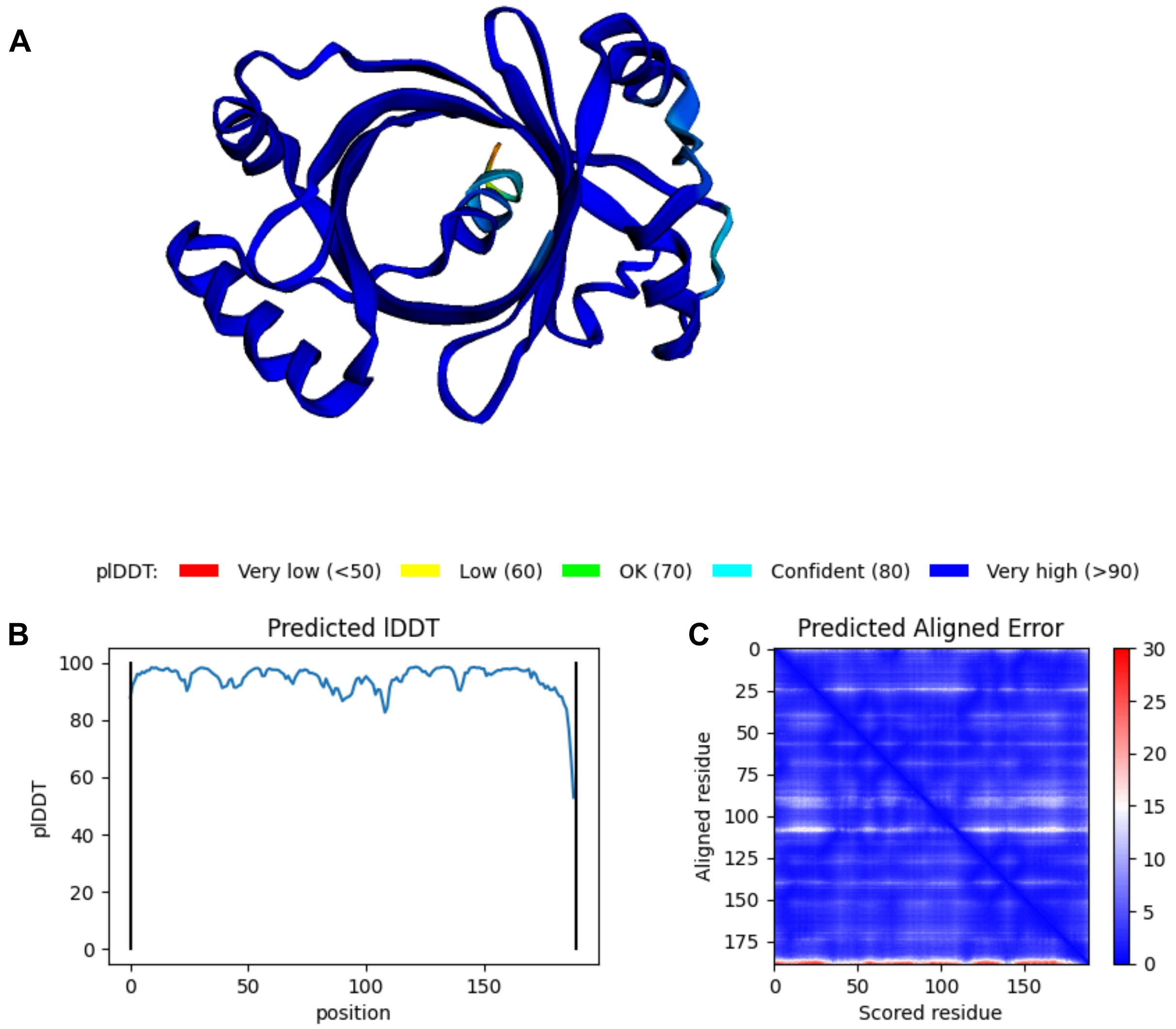

**Figure S32** Predicted structure of YjbK$_{Spn}$. (A) The amino acid sequence of RMV7$_{domi}$ *yjbK* (IONPJBJN_01639) was analysed using the AlphaFold2 pipeline with default settings. The protein structure prediction at the top shows the barrel, composed of eight beta strands, that is characteristic of CYTH proteins. The structure is coloured according to pIDDT (see Fig. S30). (B) A line graph of pIDDT over the length of the protein. This shows the algorithm has high confidence in the predicted structure. (C) A heatmap showing the uncertainty from the alignment. This shows the only region of uncertainty in the structure is at the very C terminus.

A

```
IONPJBJN_01639 MKH---LEIELKTLLKKDEYNRLKDQFTGVTPVLQ------TNYYIDTPDFELREKKVAMRIRTFEDWAELTLKVPQSVG 71
VC_2440        ME----TEIELKFFVSPDFSTILRAKIS-ETKVLQHSCRELGNTYFDTPDNWLRQHDIGLRIRRFDEVYVQTVKTAGRVV 75
HI_1598        MENIMLQEIELKLAISPQIGIELPQYLA-KFTILEHQNLFLGNTYYDYPDHFLAKQKMGLRIRQEDQELTLTLKTNGKVV 79

IONPJBJN_01639 -----NMEYNQKLQLKDAEN------------------------------------------YLSKEELPQGLVLDELAK 104
VC_2440        AGLHQRPEFNAEHHSNEPDLSLHPADIWPQGKELTQL-QAELMPLFSTNFTREQWLISMADGSQVEVAFDQGLVVAGDRQ 154
HI_1598        SGLHSRPEYNLPLIEKETPTNAQLRGLYPF----EQLPSSTLQPIFSTDFNRTFWLVEFQ-QSKIEVAFDQGKIIAGEYE 154

IONPJBJN_01639 HGI-------QSKNWQV-LGCLTTLR------------------------------------------------------ 122
VC_2440        EPICEVELELKSGQTDALFTLARQLCEHGGMRLGNLSKAARGYRLAANYSGDEIQPMALVSVDKNDTAESCFIRALEHAL 234
HI_1598        QPISEIEFELKSGNVQDLFDFVETLPFERDIYFSSASKAKRGYLLGSKQ-----------------------------FL 205

IONPJBJN_01639 ---------------------------------------------------YE--------------------------- 124
VC_2440        AHWH--YHEQIYTERENVAALHEIRHAVSYLRQLLSVYGGIIPRRASAILRQELKWLEQELQWLKEFEYLESLQEDKGYA 312
HI_1598        TDWLNKWRDFLKEEREESAVD--------------------FCAKFNAVLKMEQKLLEETLSFSPTLFSQ---------- 255

IONPJBJN_01639 ------------------------------MKTAIGL-MALDQSRYFDMTDYELELEVENHEQGKQDFRQFLEK------ 167
VC_2440        LRKLDARKFLVTALKTLQESLPQREDTLRLLSSARYTGLLLDLSRWVLTRGWQPFLDDKAREKMAQPLEAFSVKQLDRTW 392
HI_1598        -------D---------------------FMKTVERVGAFFNLYHYYDENGKILEA-MATEKQKETRLPALLES------ 300

IONPJBJN_01639 ------------------------------------------N--------QISYQKAPSKLVRFVKSMKN--------- 188
VC_2440        AELMEAFPPGKTLTVQEYLDQQYRLMRNLYTGVSFASLYDAENRQAFRMPWADLLHGI-DDLLRL--------KPLERLV 463
HI_1598        ------------------------------------------N--------QKIFVEI-RDLIRFHSETKDNEKTIEKLT 329

IONPJBJN_01639 -----------------------------------------S 189
VC_2440        DLLQGEEQDQLKRWLIRQENSILHAMEQSRTMGVEAHPYWRE 505
HI_1598        ALLKSRVY---------FE-RMIKLMELSYDL---------G 352
```

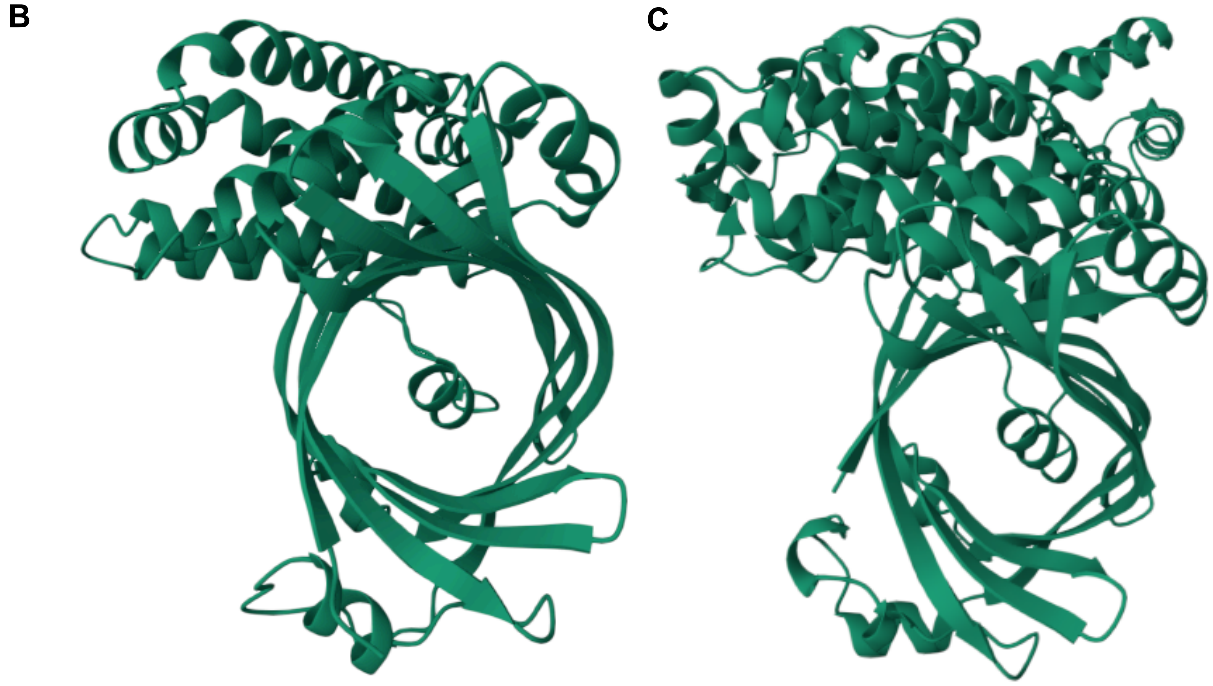

**Figure S33** (A) Alignment of CyaB adenylate cyclase proteins of *Vibrio cholerae* (VC_2440 from *V. cholerae* ATCC 39315) and *Haemophilus influenzae* (HI_1598 from *H. influenzae* Rd) with YjbK of RMV7, IONPJBJN_01639. Columns are coloured black where an amino acid is conserved at a position, and grey when amino acids are similar at a position. (B) The predicted structure of HI_1598, inferred using AlphaFold (AlphaFoldDB identifier AF-P45267-F1). This structure has a beta barrel structure, like YjbK. (C) The predicted structure of VC_2440, inferred using AlphaFold (AlphaFoldDB identifier AF-Q9KPD2-F1). This structure has a beta barrel structure, like YjbK.

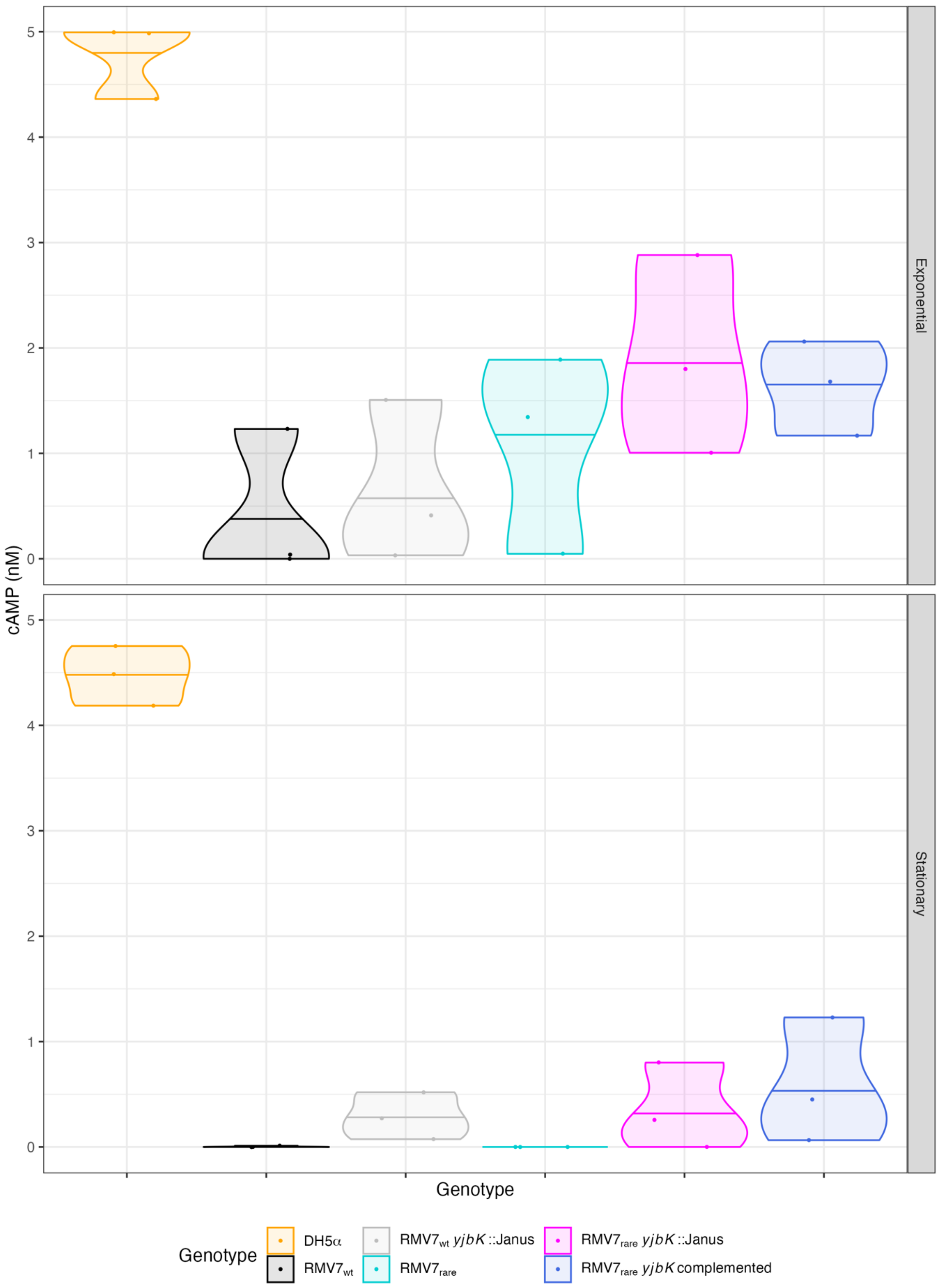

**Figure S34** Violin plots showing the concentration of 3',5'-cAMP in samples taken from exponential and stationary phase cultures of *E. coli* DH5α, and *S. pneumoniae* $RMV7_{wt}$ and $RMV7_{rare}$ genotypes differing in whether *yjbK* was intact or not. The colours indicate the genotypes from which the samples were extracted.

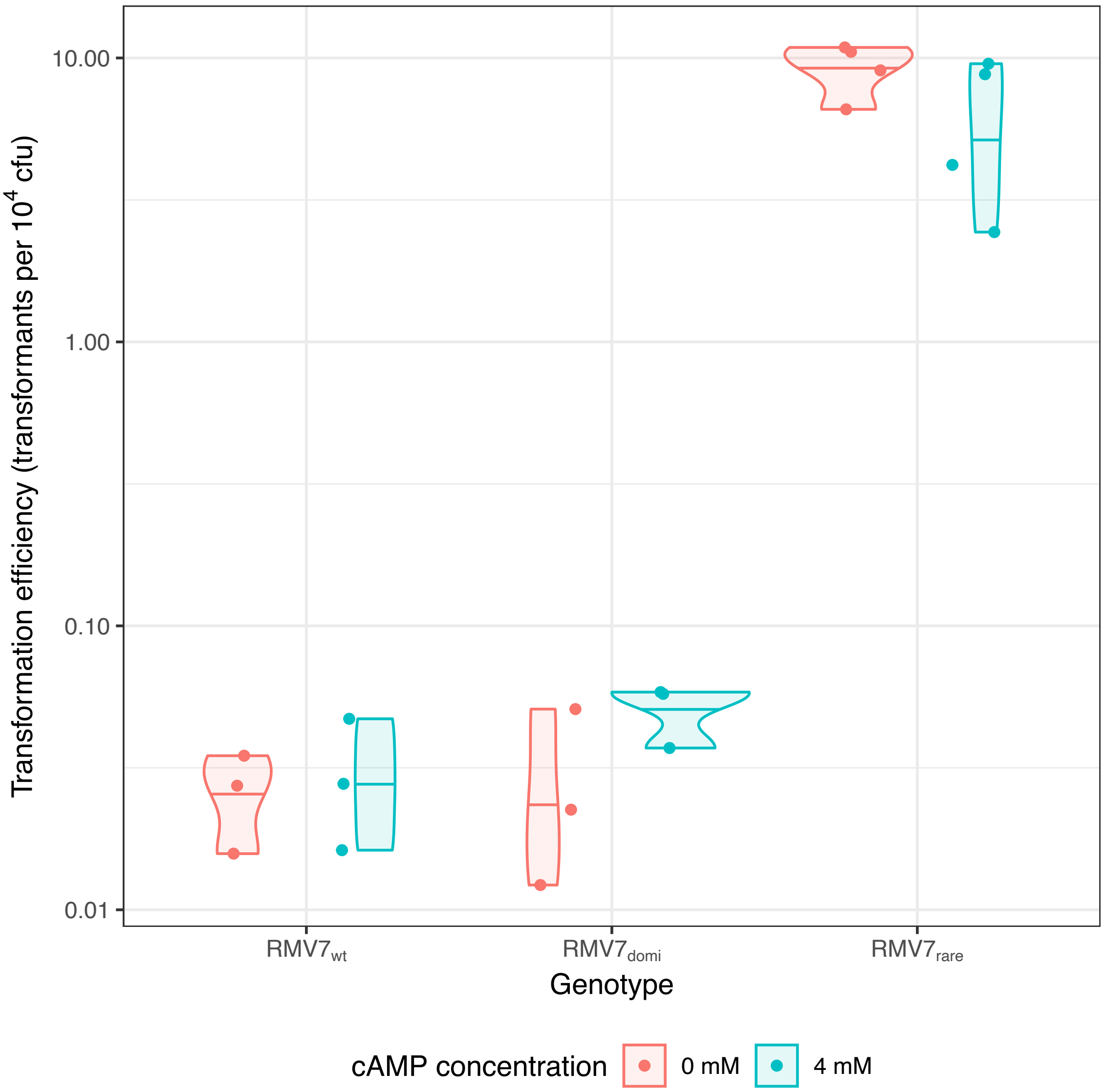

**Figure S35** Effect of exogenous 3',5'-cAMP on *S. pneumoniae* transformation efficiency. The $RMV7_{wt}$, $RMV7_{domi}$ and $RMV7_{rare}$ genotypes were transformed in liquid media with and without a supplement of 4 mM 3',5'-cAMP. This addition did not have any detectable effect on the transformation efficiency of any of the genotypes.

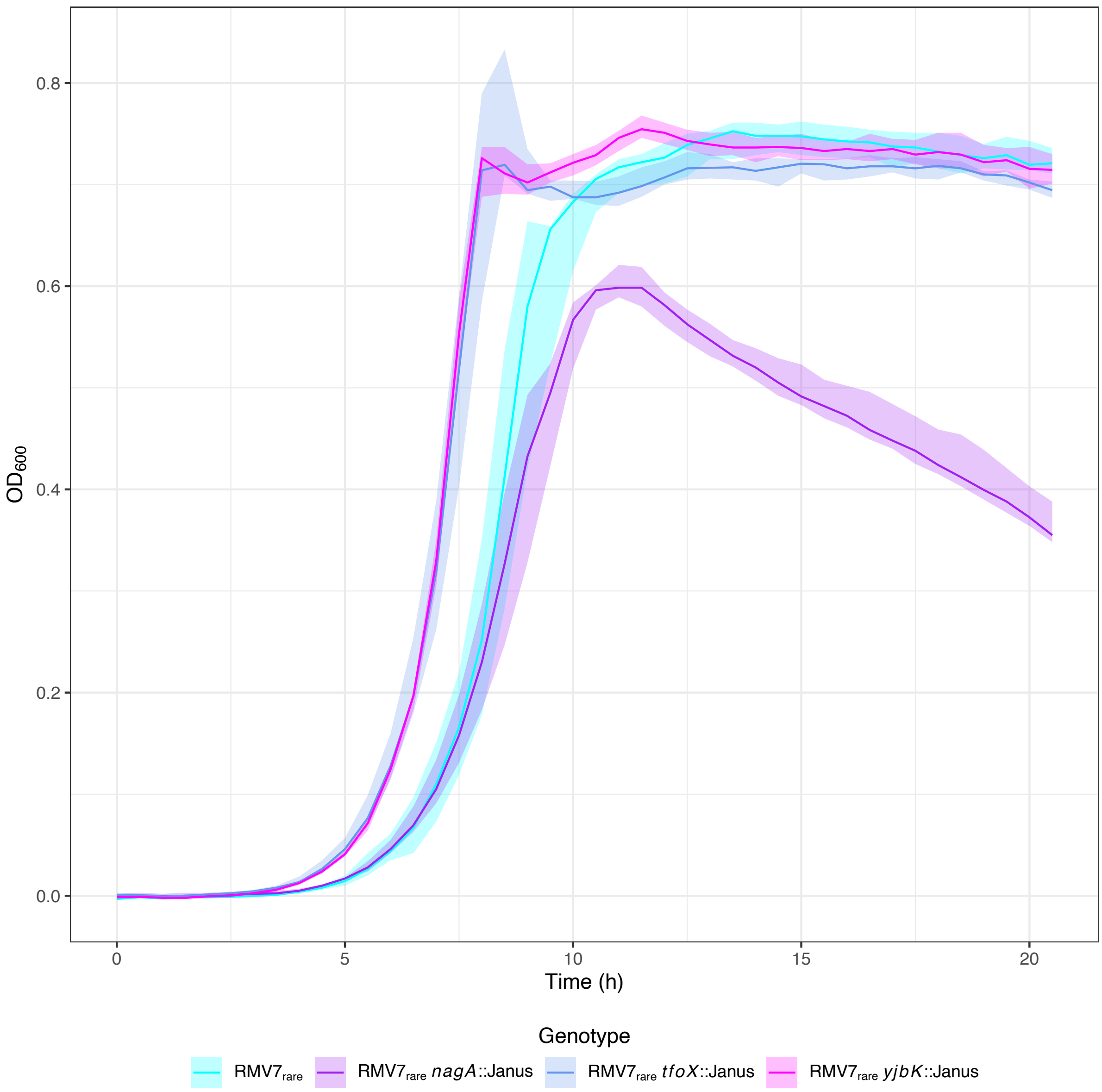

**Figure S36** Analysing GlcNAc metabolism and signalling through comparing growth curves of RMV7$_{rare}$ *nagA*::Janus, *tfoX*::Janus and *yjbK*::Janus mutants. The line graph summarises three replicate growth experiments undertaken using mixed media. The solid line shows the median recorded optical density, and the shaded area shows the range of observations.

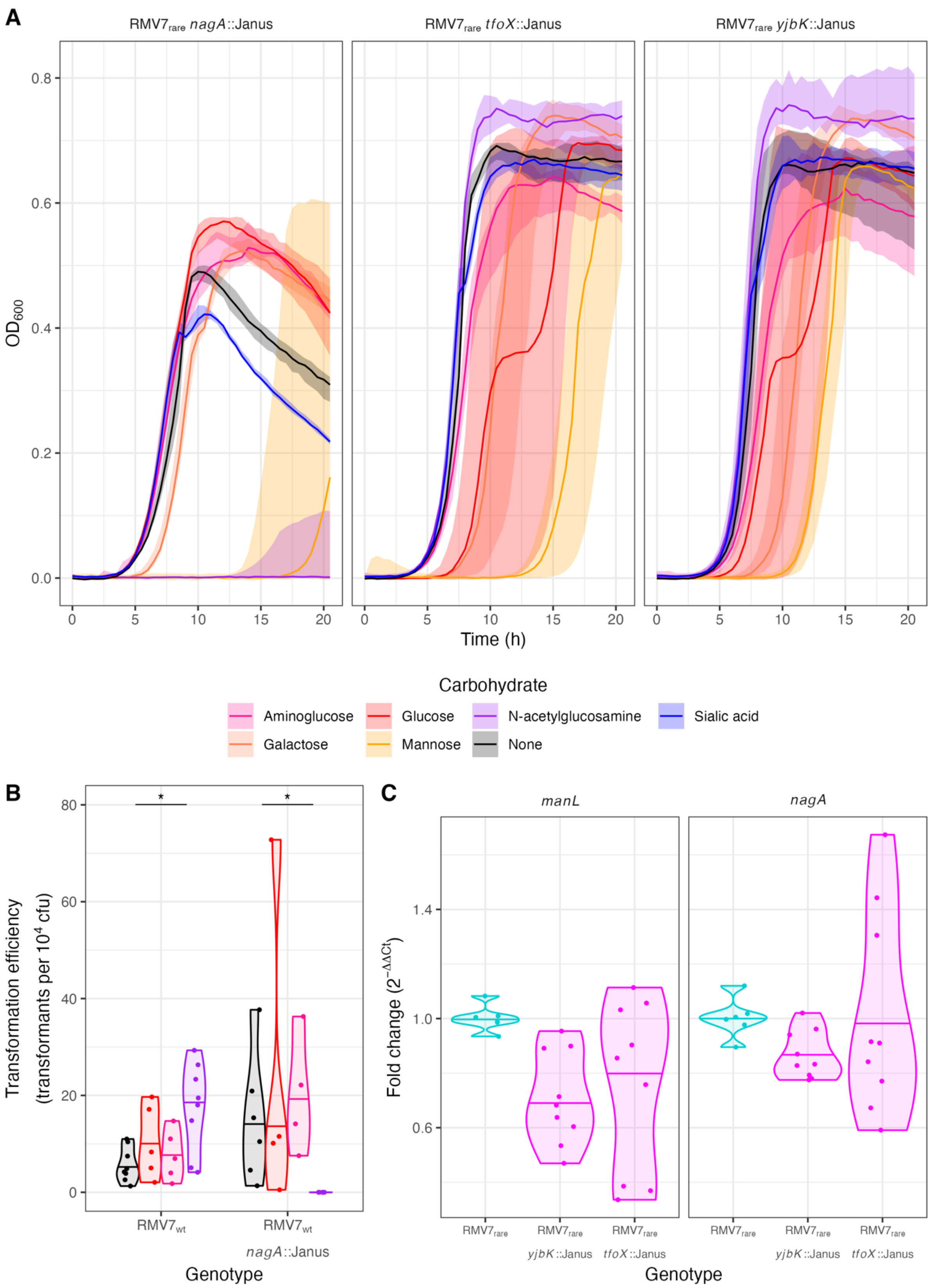

**Figure S37** Analysing GlcNAc metabolism and signalling through comparison of *nagA*::Janus, *tfoX*::Janus and *yjbK*::Janus mutants. (A) Growth curves of RMV7$_{rare}$ mutants in mixed media supplemented with different carbon sources. The *nagA*::Janus mutant exhibited a distinct growth defect in the presence of GlcNAc and mannose. This is consistent with NagA being the primary metabolic enzyme processing the cytotoxic compound GlcNAc-6-phosphate. The *tfoX*::Janus and *yjbK*::Janus mutants both grew to elevated densities in the presence of a GlcNAc supplement. (B) Transformation efficiency of $RMV7_{rare}$ *nagA*::Janus in presence of four different carbon source supplements. Consistent with the effects on growth in panel (A), no transformed colonies were recovered from the *nagA*::Janus mutant in the presence of a GlcNAc supplement. (C) Transcription of *manL* and *nagA* were measured in $RMV7_{rare}$, $RMV7_{rare}$ *tfoX*::Janus and $RMV7_{rare}$ *yjbK*::Janus grown to early exponential phase ($OD_{600}$ = 0.2). The expression of *nagA*, affecting metabolism of GlcNAc, was not affected by disruption of *tfoX* or *yjbK*. The transcription of *manL* was reduced by both mutations, suggesting the effects of *tfoX* and *yjbK* could be mediated through changing the level of the ManLMN transporter within cells.

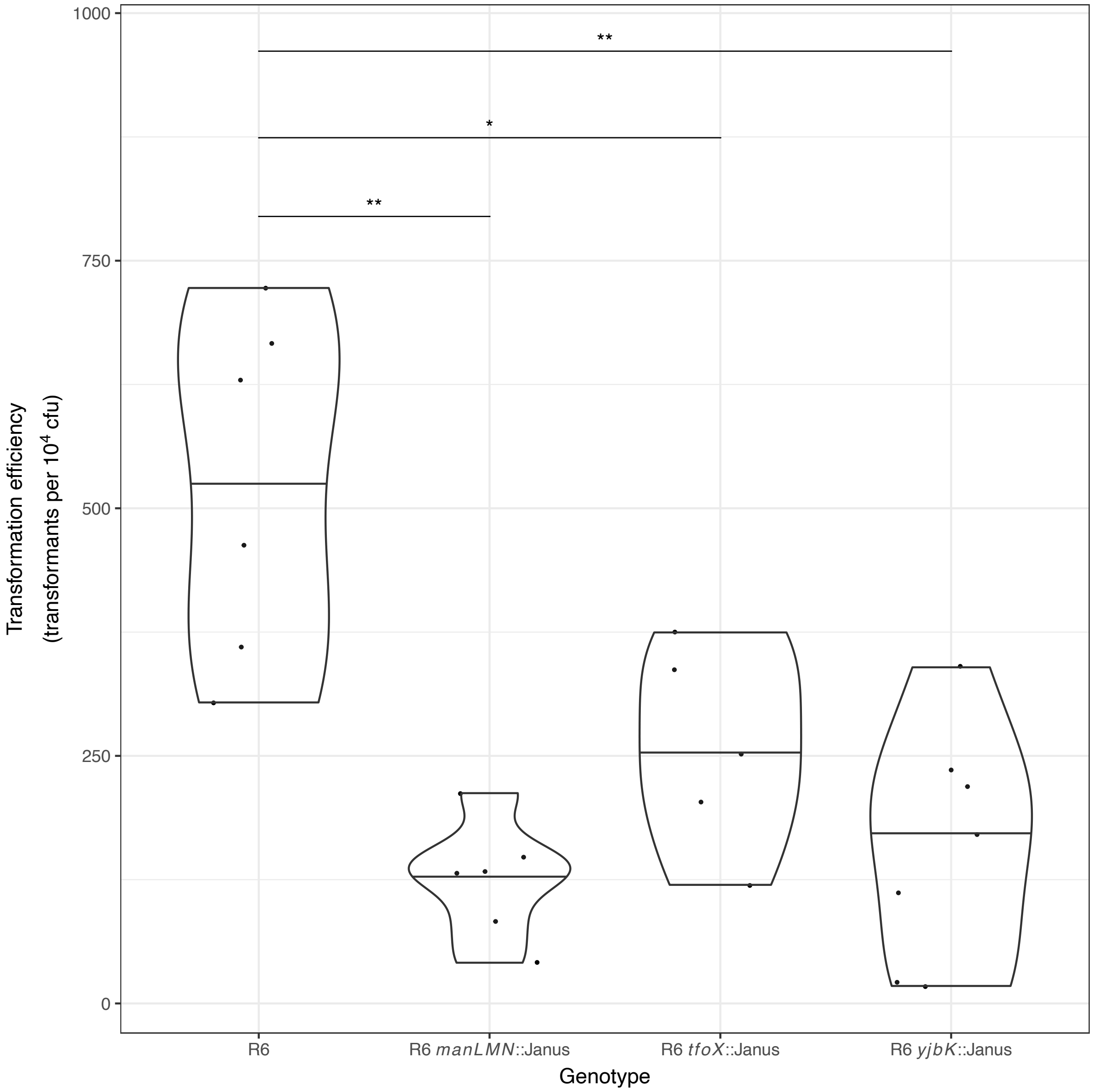

**Figure S38** Replication of the link between metabolism and transformation efficiency in the laboratory isolate *S. pneumoniae* R6. The *manLMN*, *tfoX* and *yjbK* genes were each disrupted with a Janus cassette in R6. Transformation efficiencies were measured with the bacteria grown in a nutrient-poor chemically-defined medium (see Methods). All three mutants had reduced transformation efficiencies. These differences were not observed when R6 was grown in rich media.

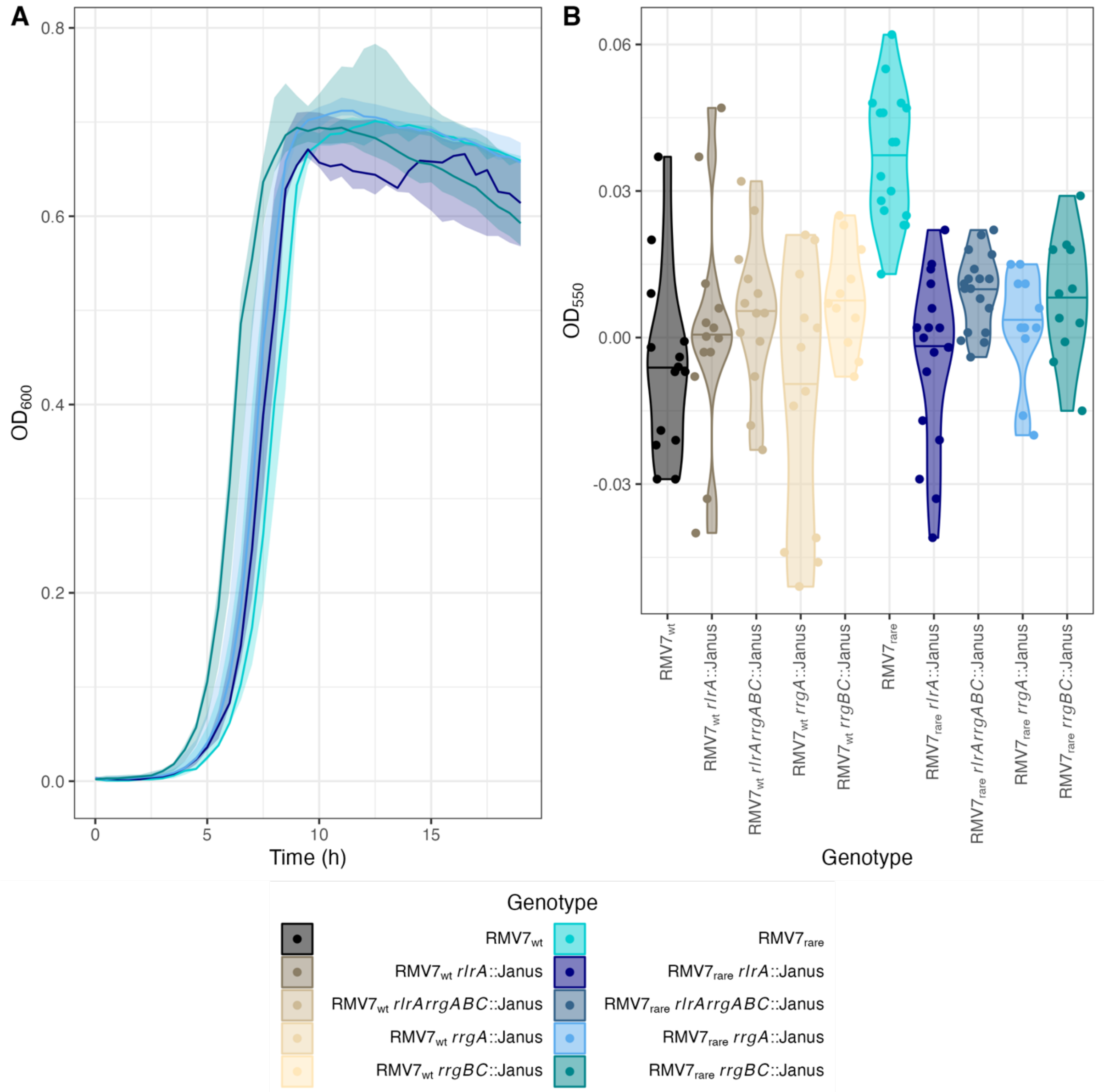

**Figure S39** Effect of type 1 pilus expression on pneumococcal phenotypes. (A) Growth curves of $RMV7_{rare}$ and mutants in which the genes for the type 1 pilus regulator (*rlrA*) or structural proteins (*rrgABC*) were disrupted using a Janus cassette. The median (solid line) and range (shaded area) of observations were calculated from three replicates. The loss of the pilus genes has little effect on growth. (B) Biofilm formation of RMV7 genotypes and mutant derivatives lacking type 1 pili. The greatest adherence of cells to the abiotic surface was observed with the $RMV7_{rare}$ variant, which expressed the type 1 pilus more strongly than the $RMV7_{wt}$ variant. Correspondingly, loss of the regulatory or structural pilus genes had little effect on biofilm formation in $RMV7_{wt}$, but substantially decreased biofilm formation by $RMV7_{rare}$.

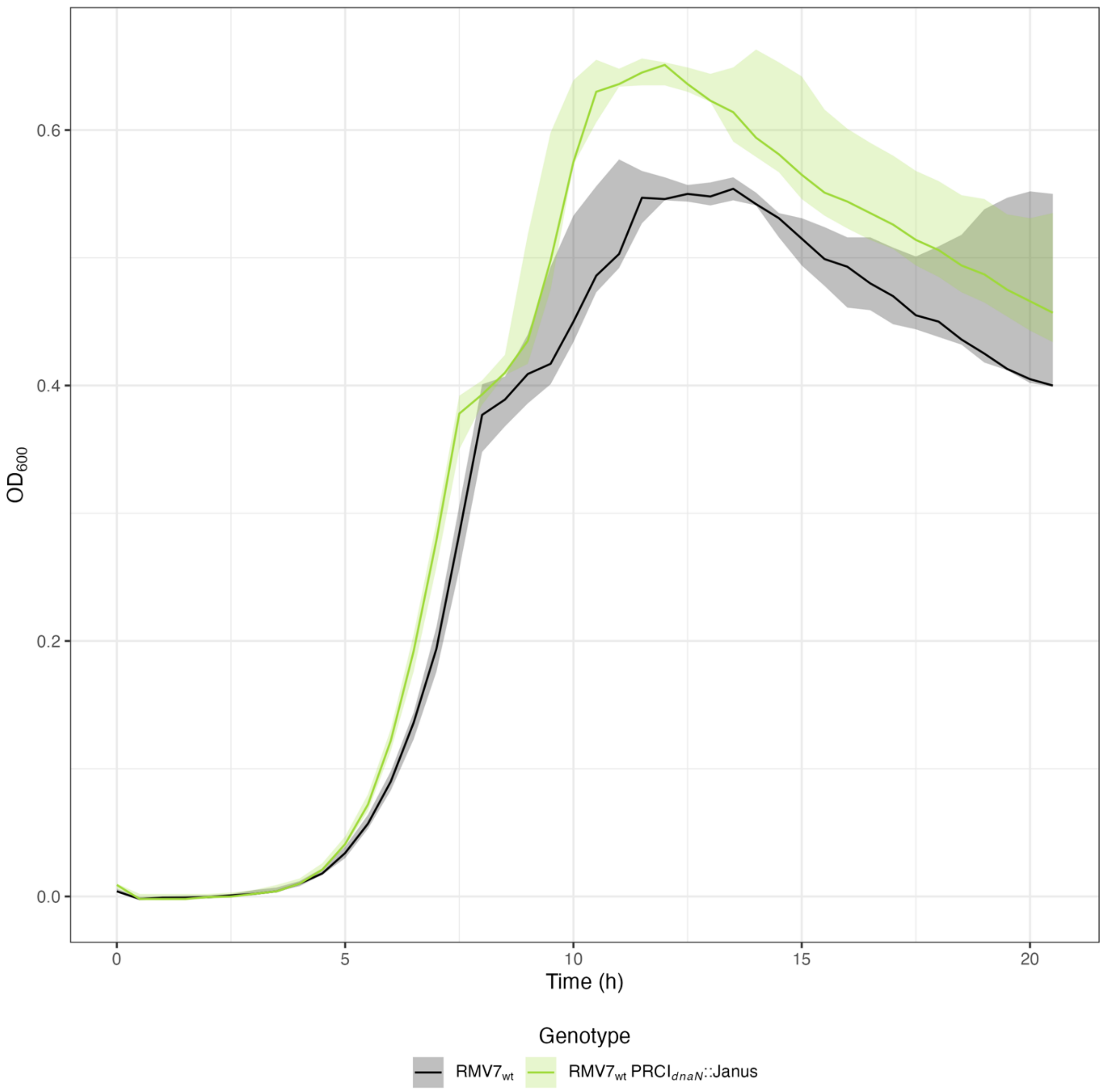

**Figure S40** Growth curves of RMV7$_{wt}$ and RMV7$_{wt}$ PRCI$_{dnaN}$::Janus in mixed media. In each plot, the lines represent the median $OD_{600}$, and the shaded ribbon shows the range of three replicates. This demonstrates that the deletion of PRCI$_{dnaN}$ increases the growth rate, and carrying capacity, of the RMV7$_{wt}$, suggesting the activity of this element causes intracellular stress.

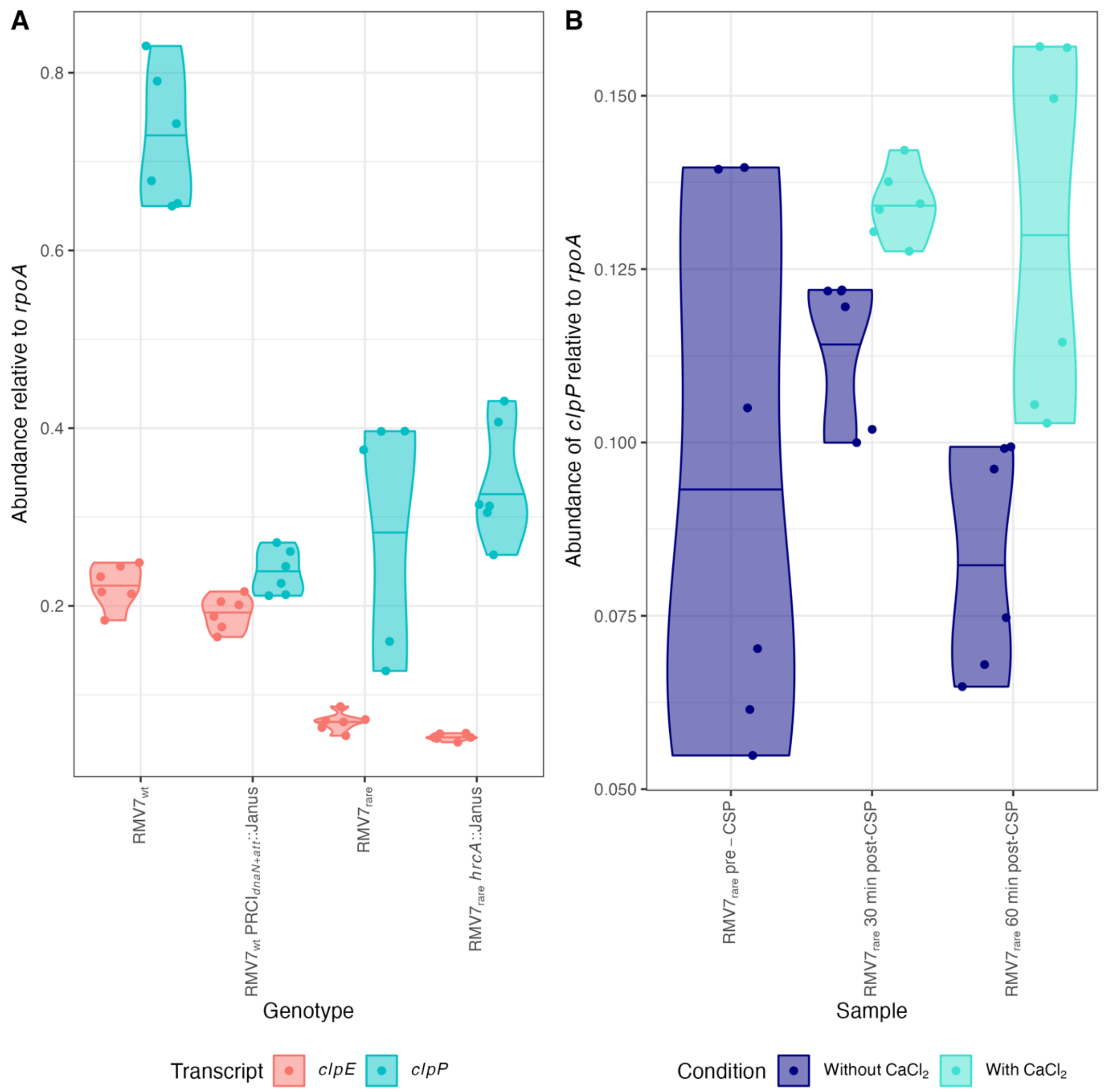

**Figure S41** Expression of *clpE* and *clpP*. (A) Violin plot showing the relative abundance of *clpE* and *clpP* transcripts in different genotypes. The *clpP* gene was most highly transcribed in $RMV7_{wt}$, in which the PRCI is highly active. The absolute expression of *clpP*, and the expression relative to *clpE*, was reduced when the PRCI was removed from $RMV7_{wt}$, suggesting expression was induced by the stress of the element's activity. Correspondingly, *clpP* expression was similarly low in $RMV7_{rare}$, in which the PRCI is less active. The deletion of *hrcA* had relatively little effect on the expression of *clpE* or *clpP* in the absence of competence induction. (B) Violin plot showing the effect of CSP and $Ca^{2+}$ on *clpP* expression in $RMV7_{rare}$. In the absence of a $Ca^{2+}$ supplement, relatively little change in *clpP* expression was observed after the induction of competence. The addition of a $Ca^{2+}$ supplement elevated expression of *clpP*. This likely reflected the increased suppression of other chaperones, due to the increased binding of HrcA to CIRCE motifs. Hence although ClpP does not appear to be part of the direct HrcA regulon (panel A), it is likely to be indirectly affected by the overlap with the functional roles of other chaperone proteins within the cell following the induction of competence.

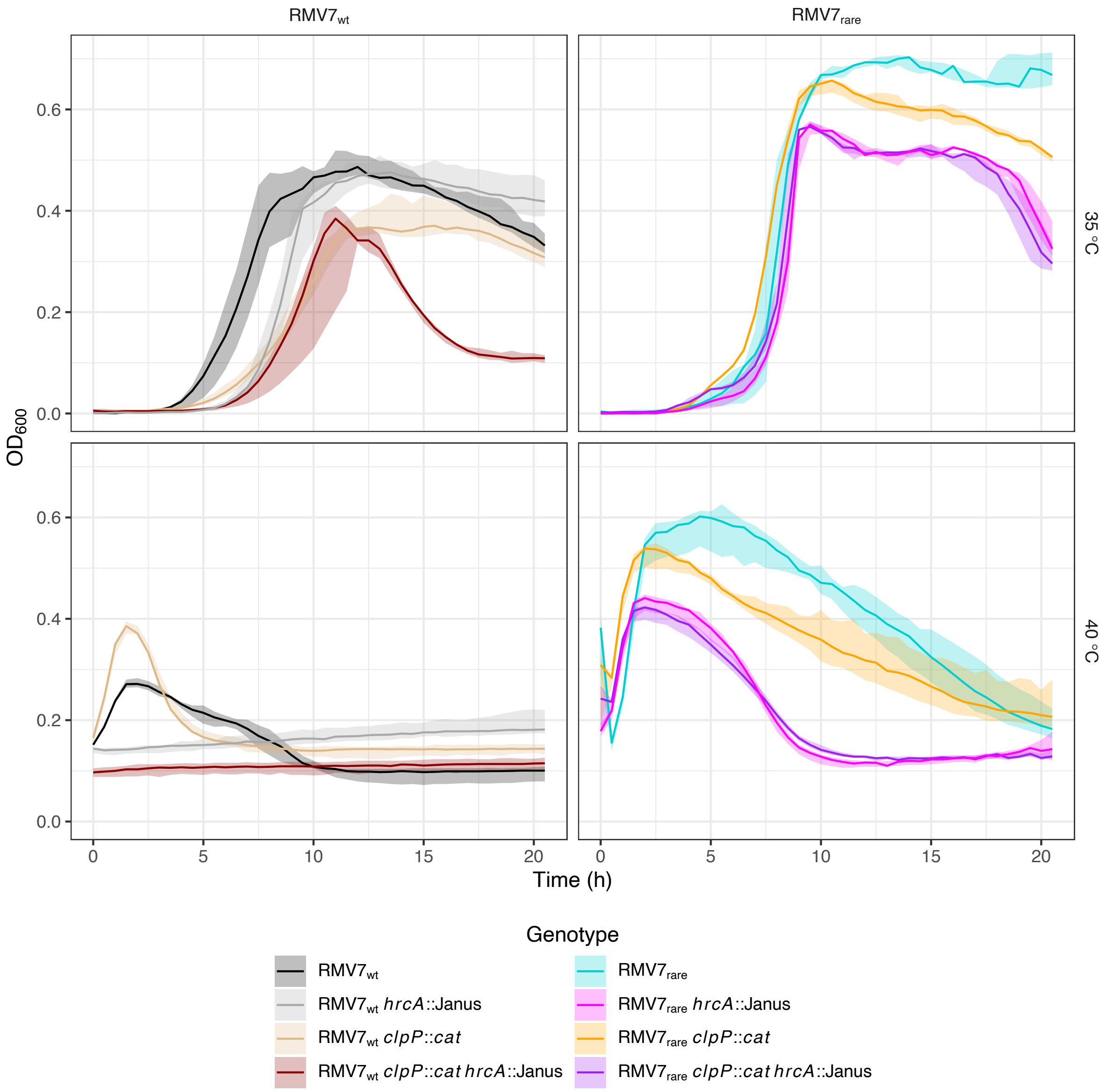

**Figure S42** Growth curves of RMV7$_{wt}$ and RMV7$_{rare}$ chaperone mutants in mixed media. In each plot, the lines represent the median OD$_{600}$, and the shaded ribbon shows the range of three replicates. The growth of RMV7$_{wt}$ genotypes is shown on the left, with RMV7$_{rare}$ genotypes on the right. The top row shows the growth of cells under standard culturing conditions, whereas the bottom row shows the growth of pneumococci at 40 °C. These “heat shock” assays required first growing the cells at 35 °C to an OD$_{600}$ of 0.2, to establish replication, before then being grown at 40 °C over a further 20 hours.

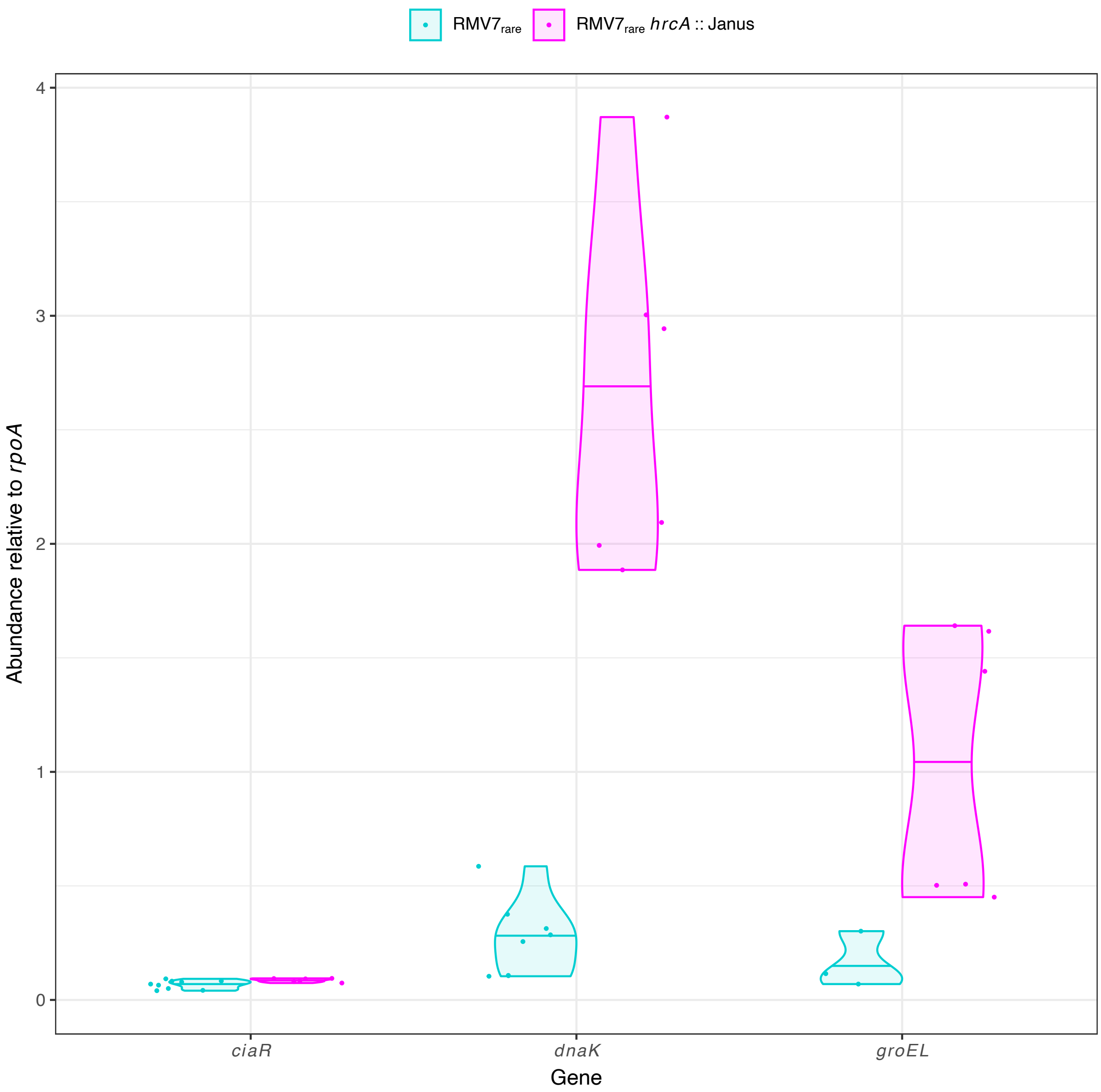

**Figure S43** Violin plots showing gene expression, as quantified by qRT-PCR, in RMV7$_{rare}$ and a mutant derivative in which the *hrcA* chaperone regulator gene was disrupted by a Janus cassette. The six points for each gene correspond to three technical replicate assays of each of two biological replicates. The horizontal line on the violin plot shows the median relative abundance for each gene in each genotype. The absence of any change in *ciaR* expression demonstrated the effect on transformation was independent of CiaRH.

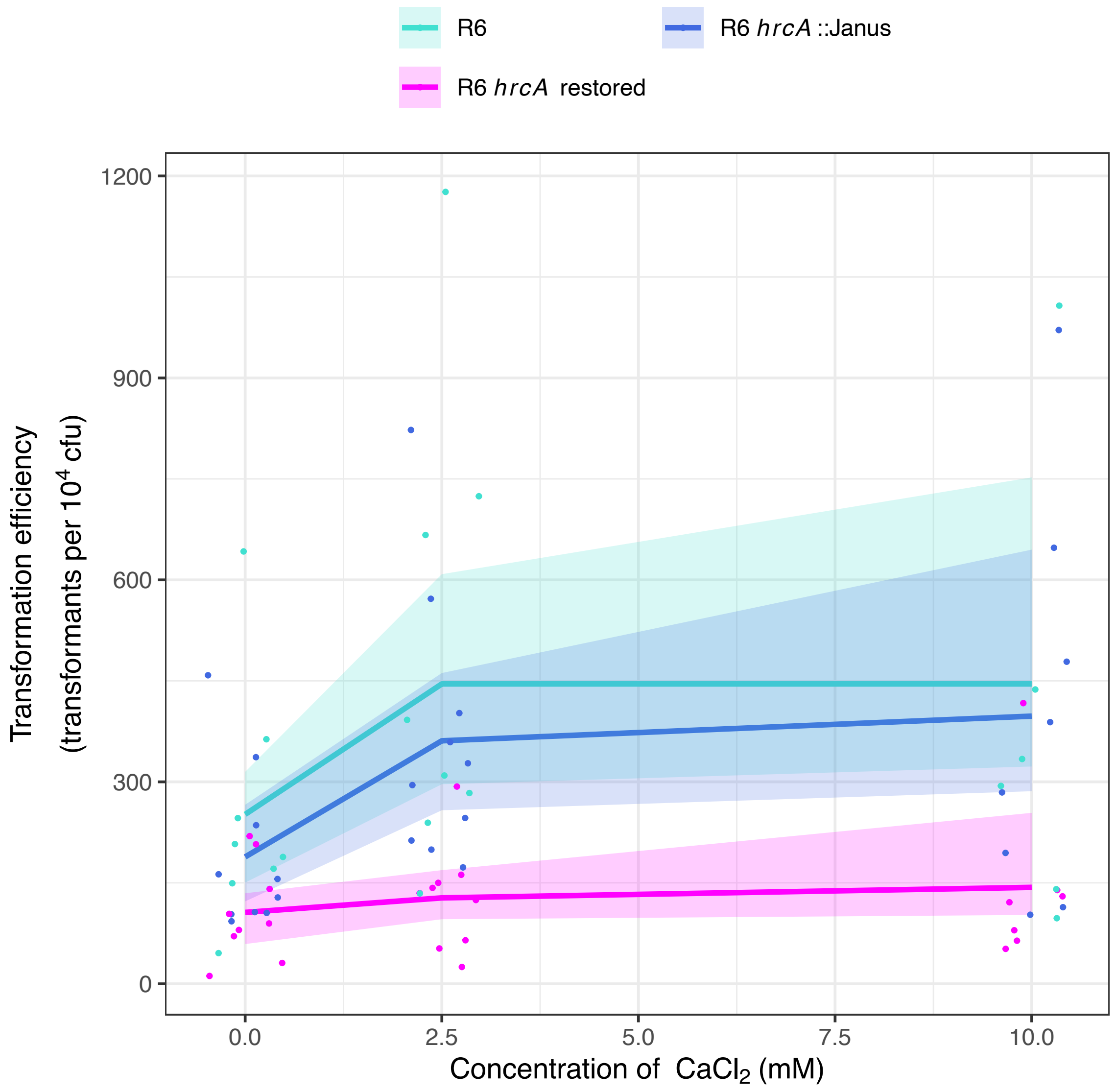

**Figure S44** Replication of the HrcA-dependent $Ca^{2+}$-activation of transformation in the laboratory isolate *S. pneumoniae* R6. The *hrcA* gene was disrupted by a Janus cassette, and then restored. Each point represents the transformation efficiency of an individual experiment. The solid lines represent the best fit of a dose-response logistic model. The shaded areas correspond to the 95% confidence intervals. Both of the *hrcA*$^+$ genotypes responded to increasing concentrations of $Ca^{2+}$ ions, but the genotype in which *hrcA* was disrupted did not.

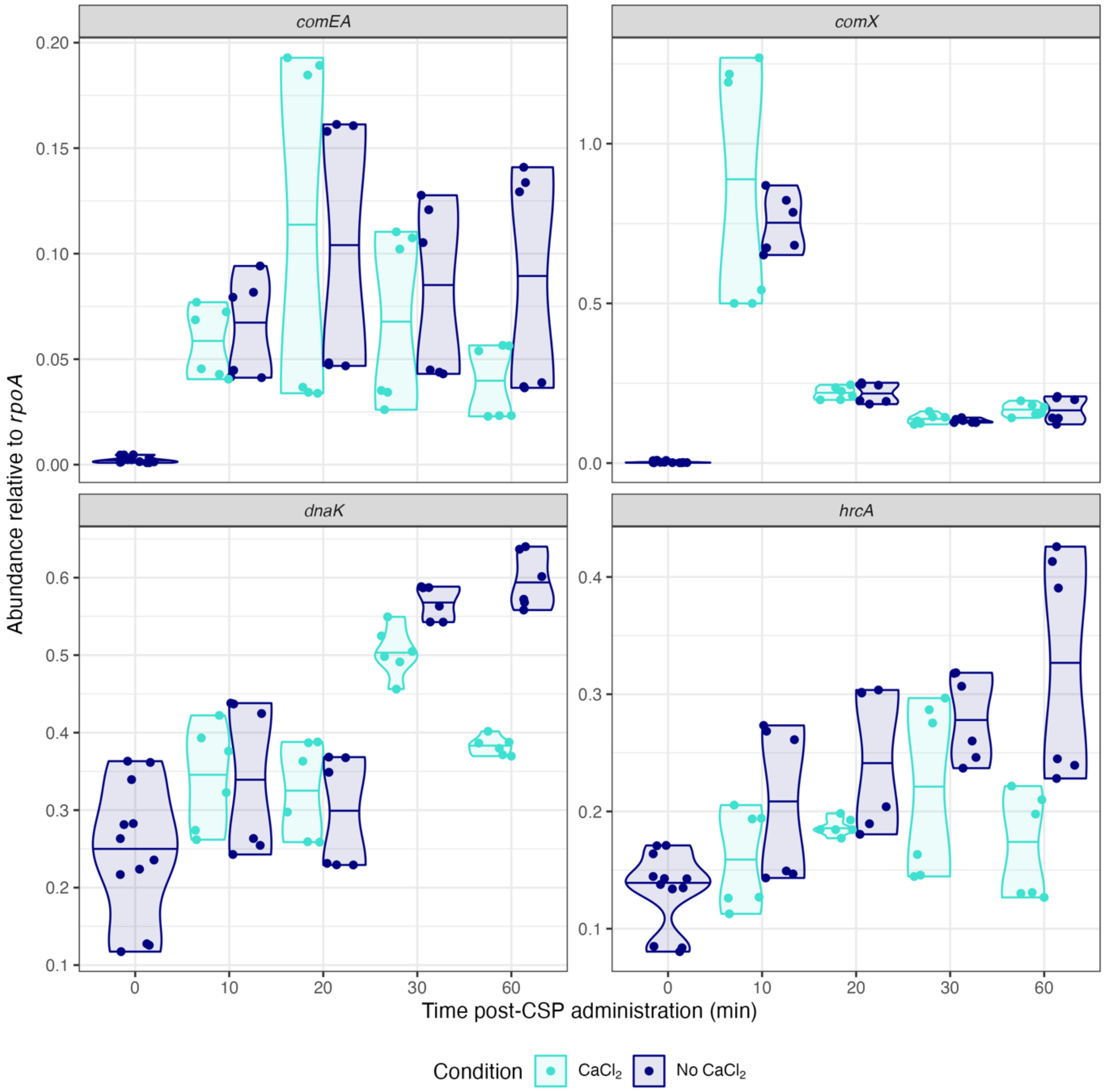

**Figure S45** Violin plots showing the effect of $Ca^{2+}$ signalling on the expression of genes regulated by HrcA after the administration of CSP. Transcription of each gene was quantified relative to *rpoA* by qRT-PCR, using three technical replicate measurements of each of two biological replicates. $Ca^{2+}$ supplementation increases HrcA's affinity for DNA, and as the protein autorepresses, this decreases *hrcA* transcription. However, this response is only evident after 30-60 min post-CSP. Similarly, *dnaK* is known to be repressed by HrcA, and there is evidence of a similar 30-60 min delay before this gene's transcription declines. By contrast, there was little evidence of $Ca^{2+}$-associated repression of either the early competence gene *comX*, or the late competence gene *comEA*. This suggests HrcA is unlikely to affect the induction of competence through repressing expression of the machinery required for transformation.

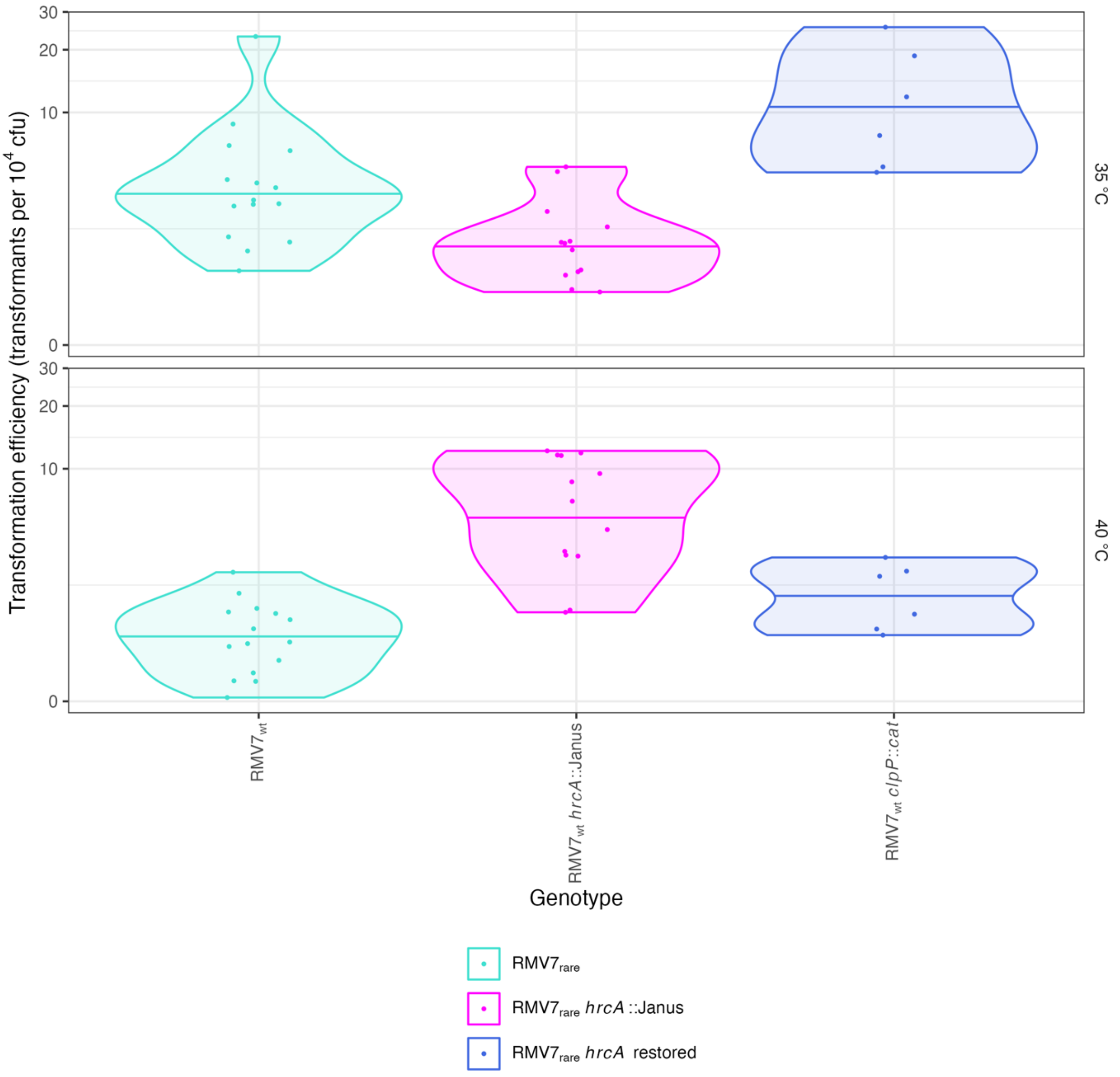

**Figure S46** Violin plots showing the transformation efficiency of $RMV7_{rare}$ and mutant derivatives during normal growth (35 °C) or during a 40 °C heat shock. The *hrcA* gene was disrupted in $RMV7_{rare}$ *hrcA*::Janus, and reinstated in $RMV7_{rare}$ *hrcA* restored. Data for $RMV7_{rare}$ and $RMV7_{rare}$ *hrcA*::Janus are reproduced from Fig. 5A. Transformation experiments for $RMV7_{rare}$ *hrcA* restored were conducted separately, but nevertheless show reinstating the *hrcA* gene reverses both the decrease in transformation efficiency at 35 °C, and the increase in transformation efficiency at 40 °C. Each point corresponds to an independent transformation experiment, and the violin plots have a horizontal line indicating the median transformation efficiency of each mutant at each temperature.

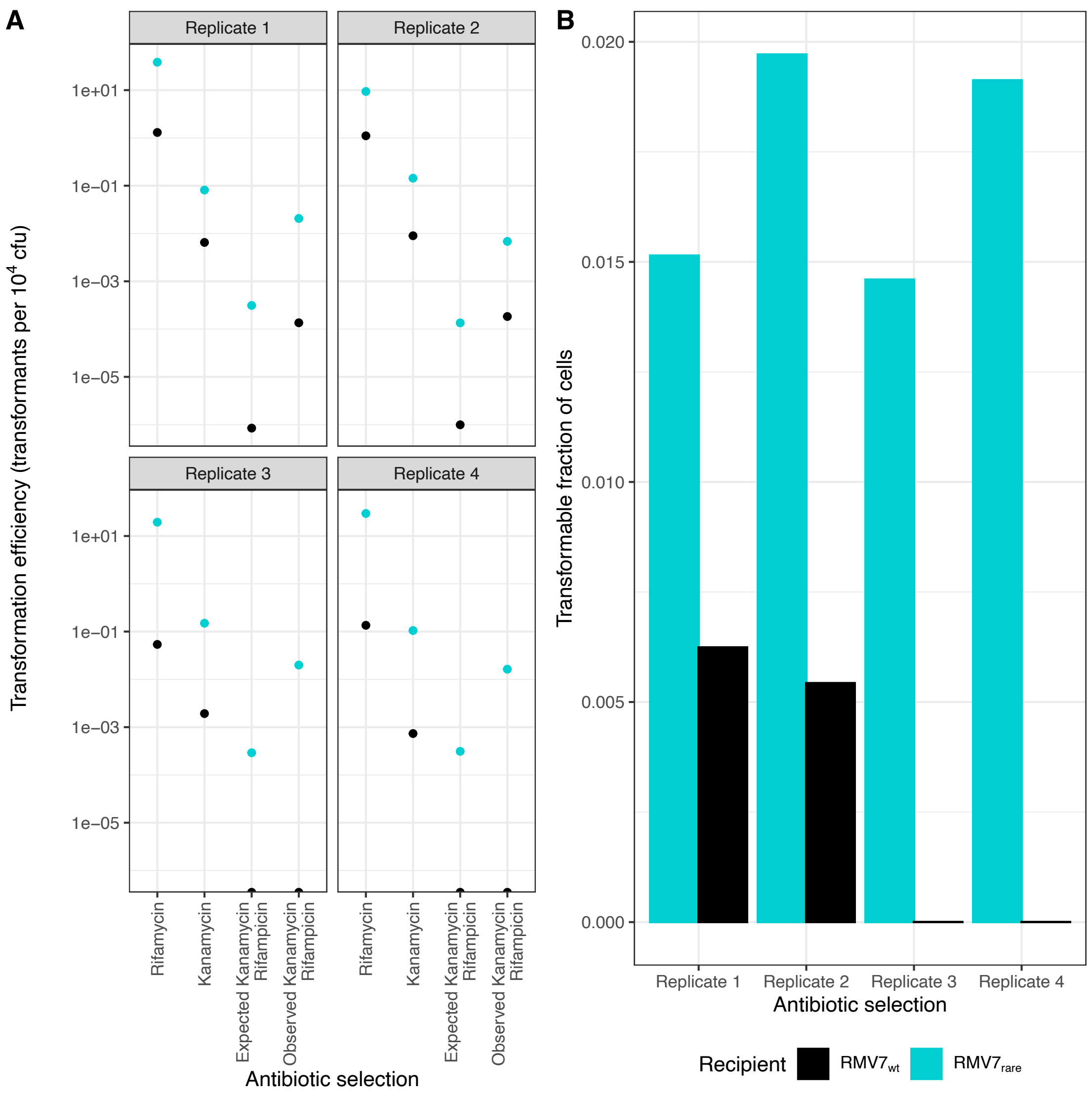

**Figure S47** Heterogeneous induction of competence across the pneumococcal population. (A) Four independent replicate experiments on separate cultures of $RMV7_{rare}$ and $RMV7_{wt}$ measured the frequency of transformation with kanamycin and rifampicin resistance markers, each on separate PCR amplicons, in the presence of GlcNAc. Assuming each marker was acquired independently, the expected frequency of double mutants was estimated from the observed frequencies of the single mutants. The observed frequencies of double mutants were higher than those expected for all four replicates with the $RMV7_{rare}$ genotype, and two of the four replicates for the less-transformable $RM7_{wt}$ genotype; in the other two replicates, the expected frequencies of double mutants were much below one cell per experiment, limiting the power to detect a deviation from this number. The excess of observed double mutants, above the expected frequency, is consistent with only a subpopulation of pneumococci within the culture being transformable. (B) Bar plot showing the estimated fraction of pneumococci that are transformable. The ratio of the expected to observed double mutant frequencies is an estimate of the transformable fraction of the pneumococcal cultures (see Methods). For $RMV7_{rare}$, the estimates are consistently close to ~1.5% of the population being transformable. For $RMV7_{wt}$, the estimates are closer to 0.5%. This relatively small fold difference between the two variants reflects the disproportionate effect of GlcNAc on $RMV7_{wt}$ relative to $RMV7_{rare}$ (Fig. 3).

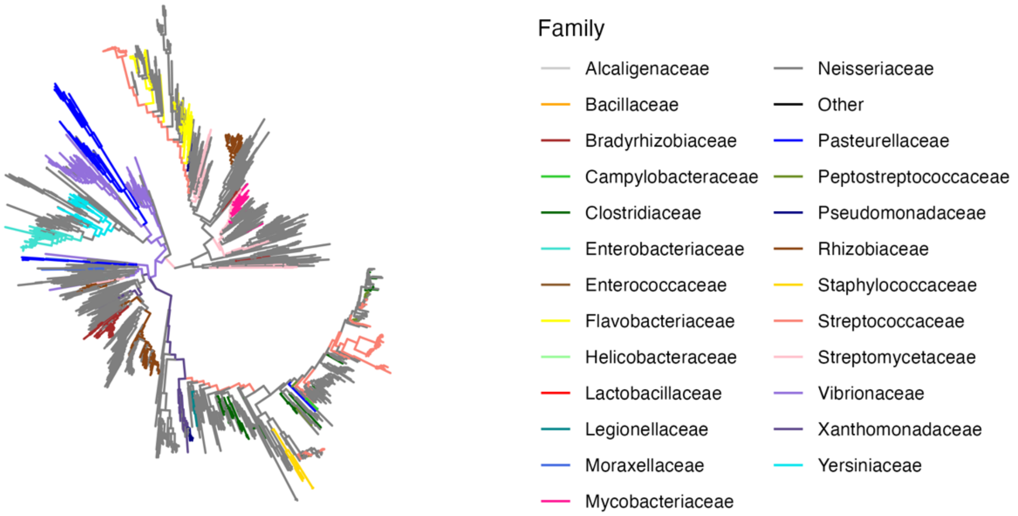

**Figure S48** Maximum likelihood phylogeny of 1,468 TfoX proteins. These sequences were identified using the EMBL SMART server and aligned with MAFFT. A phylogeny was generated with Fasttree2. Tips were assigned to families when they arose from a known taxon. These were used to partition and colour the phylogeny, according to the displayed legend, using ggtree. The clade shared by *H. influenzae* (Pasteurellaceae) and *V. cholerae* (Vibrionaceae) is distinct from that shared by *S. pneumoniae* (Streptococcaceae) and *Staphylococcus aureus* (Staphylococcaceae).

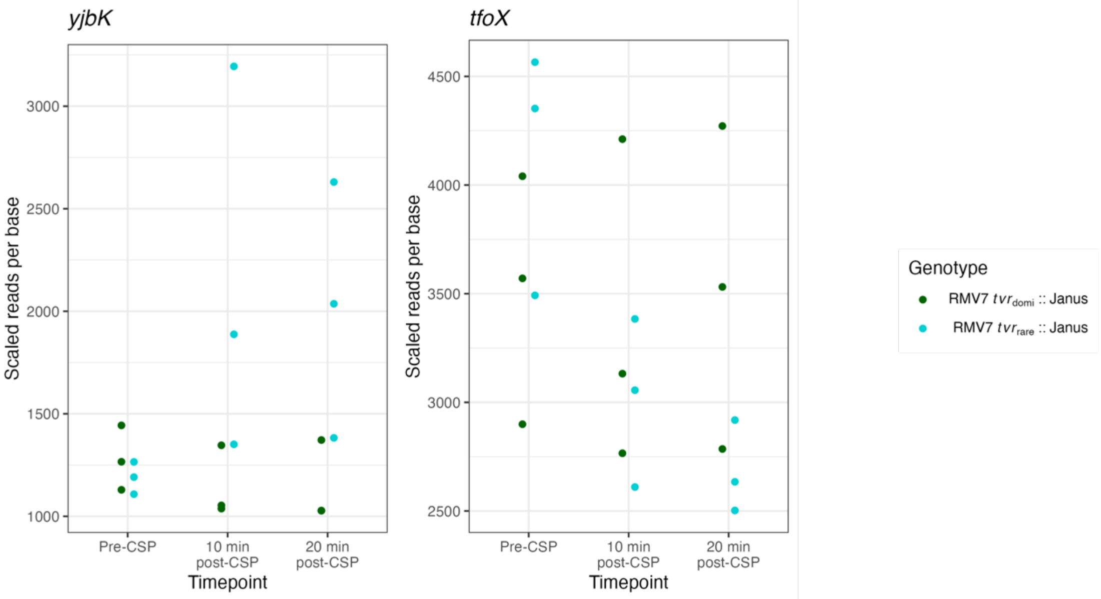

**Figure S49** Quantification of the expression of *yjbK* and *tfoX* using RNA-seq data. Data are displayed as in Fig. S11. Both genes are constitutively expressed, with the main difference between the variants being an apparent induction of *yjbK* expression after the addition of CSP.

**Supplementary Tables**

**Table S1** Single nucleotide polymorphisms distinguishing $RMV7_{domi}$ and $RMV7_{rare}$.

| Position $RMV7_{domi}$ | Allele in $RMV7_{domi}$ | Allele in $RMV7_{rare}$ | Type | Affected gene | Description |
|---|---|---|---|---|---|
| 1692817 | G | A | Non-synonymous | IONPJBJN_00426 (*ktrA*) | Non-synonymous mutation in potassium transporter |
| 1708252 | A | . | Intergenic | - | Mutation upstream of gene encoding hypothetical protein |
| 1741214 | . | C | Intergenic | - | Mutation in homopolymeric tract upstream of lytic amidase |
| 1901001 | A | G | Non-synonymous | IONPJBJN_00199 (*xerC_2)* | Non-synonymous mutation in mobile element integrase |
| 1992363 | A | G | Premature stop | IONPJBJN_00114 | Premature stop codon affecting membrane protein in $RMV7_{domi}$ |
| 279792 | G | A | Premature stop | IONPJBJN_01863 (*pstS1*) | Premature stop codon in $RMV7_{rare}$ |
| 645400 | A | C | Synonymous | IONPJBJN_01511 (*uvrC*) | Synonymous change in gene encoding DNA repair protein |
| 736520 | T | C | Variation in *tvr* locus | IONPJBJN_01427 | Variation in rearranged *hsdS* gene |
| 736578 | G | A | Variation in *tvr* locus | IONPJBJN_01427 | Variation in rearranged *hsdS* gene |
| 736581 | A | C | Variation in *tvr* locus | IONPJBJN_01427 | Variation in rearranged *hsdS* gene |
| 736620 | A | G | Variation in *tvr* locus | IONPJBJN_01427 | Variation in rearranged *hsdS* gene |
| 736681 | C | G | Variation in *tvr* locus | IONPJBJN_01427 | Variation in rearranged *hsdS* gene |
| 736916 | C | T | Variation in *tvr* locus | IONPJBJN_01426 (*xerC_3*) | Variation in *tvrR* gene within rearranged *tvr* locus |

| | | | | | |
|---|---|---|---|---|---|
| 736970 | A | G | Variation in *tvr* locus | IONPJBJN_01426 (*xerC_3*) | Variation in *tvrR* gene within rearranged *tvr* locus |
| 736990 | C | T | Variation in *tvr* locus | IONPJBJN_01426 (*xerC_3*) | Variation in *tvrR* gene within rearranged *tvr* locus |
| 737012 | T | A | Variation in *tvr* locus | IONPJBJN_01426 (*xerC_3*) | Variation in *tvrR* gene within rearranged *tvr* locus |
| 737063 | T | C | Variation in *tvr* locus | IONPJBJN_01426 (*xerC_3*) | Variation in *tvrR* gene within rearranged *tvr* locus |
| 737227 | A | G | Variation in *tvr* locus | IONPJBJN_01426 (*xerC_3*) | Variation in *tvrR* gene within rearranged *tvr* locus |
| 737341 | T | C | Variation in *tvr* locus | IONPJBJN_01426 (*xerC_3*) | Variation in *tvrR* gene within rearranged *tvr* locus |
| 925619 | C | A | Non-synonymous | IONPJBJN_01237 (*clpX*) | Non-synonymous mutation in gene encoding a protease |
| 1181193 | C | A | Non-synonymous | IONPJBJN_00977 (*feuB*) | Non-synonymous mutation in *piaA* iron transporter gene |
| 1361856 | G | T | Non-synonymous | IONPJBJN_00765 | Non-synonymous mutation in gene encoding an ion transporter protein |
| 1415649 | T | C | Non-synonymous | IONPJBJN_00700 (*gltX*) | Synonymous change in gene encoding glutamate--tRNA ligase |
| 1551834 | T | C | Non-synonymous | IONPJBJN_00578 (*phoB*) | Non-synonymous mutation in gene encoding a regulatory protein |

**Table S2** Accession codes for RNA-seq datasets.

| Accession code | Genotype | Timepoint |
|---|---|---|
| ERS3382236 | RMV7 *tvr*$_{rare}$::Janus | Pre-CSP |
| ERS3382237 | RMV7 *tvr*$_{rare}$::Janus | 10 min post-CSP |
| ERS3382238 | RMV7 *tvr*$_{rare}$::Janus | 20 min post-CSP |
| ERS3382239 | RMV7 *tvr*$_{domi}$::Janus | Pre-CSP |
| ERS3382240 | RMV7 *tvr*$_{domi}$::Janus | 10 min post-CSP |
| ERS3382241 | RMV7 *tvr*$_{domi}$::Janus | 20 min post-CSP |
| ERS3382242 | RMV7 *tvr*$_{rare}$::Janus | Pre-CSP |
| ERS3382243 | RMV7 *tvr*$_{rare}$::Janus | 10 min post-CSP |
| ERS3382244 | RMV7 *tvr*$_{rare}$::Janus | 20 min post-CSP |
| ERS3382227 | RMV7 *tvr*$_{domi}$::Janus | Pre-CSP |
| ERS3382228 | RMV7 *tvr*$_{domi}$::Janus | 10 min post-CSP |
| ERS3382229 | RMV7 *tvr*$_{domi}$::Janus | 20 min post-CSP |
| ERS3382230 | RMV7 *tvr*$_{rare}$::Janus | Pre-CSP |
| ERS3382231 | RMV7 *tvr*$_{rare}$::Janus | 10 min post-CSP |
| ERS3382232 | RMV7 *tvr*$_{rare}$::Janus | 20 min post-CSP |
| ERS3382233 | RMV7 *tvr*$_{domi}$::Janus | Pre-CSP |
| ERS3382234 | RMV7 *tvr*$_{domi}$::Janus | 10 min post-CSP |
| ERS3382235 | RMV7 *tvr*$_{domi}$::Janus | 20 min post-CSP |

**Table S3** Differential gene expression analysis (see spreadsheet)

**Table S4** Logistic model parameters estimated from growth curves.

| **Genotype** | **Condition** | **Carrying capacity estimate, *K*** | **Carrying capacity standard error** | **Growth rate estimate, *r* ($h^{-1}$)** | **Growth rate standard error ($h^{-1}$)** | **Figure** |
|---|---|---|---|---|---|---|
| RMV7$_{rare}$ | Aminoglucose | 0.734 | 0.001 | 1.206 | 0.009 | Fig. S26 |
| RMV7$_{rare}$ | Galactose | 0.719 | 0.002 | 1.292 | 0.028 | Fig. S26 |
| RMV7$_{rare}$ | Glucose | 0.718 | 0.006 | 1.624 | 0.115 | Fig. S26 |
| RMV7$_{rare}$ | Mannose | 0.616 | 0.006 | 1.352 | 0.104 | Fig. S26 |
| RMV7$_{rare}$ | N-acetylglucosamine | 0.756 | 0.004 | 1.472 | 0.064 | Fig. S26 |
| RMV7$_{rare}$ | None | 0.687 | 0.003 | 1.552 | 0.055 | Fig. S26 |
| RMV7$_{rare}$ | Sialic acid | 0.718 | 0.004 | 1.114 | 0.041 | Fig. S26 |
| RMV7$_{rare}$ *manLMN*::Janus | Aminoglucose | 0.801 | 0.005 | 1.252 | 0.062 | Fig. S26 |
| RMV7$_{rare}$ *manLMN*::Janus | Galactose | 0.773 | 0.006 | 1.265 | 0.077 | Fig. S26 |
| RMV7$_{rare}$ *manLMN*::Janus | Glucose | 0.750 | 0.007 | 1.350 | 0.106 | Fig. S26 |
| RMV7$_{rare}$ *manLMN*::Janus | Mannose | 0.738 | 0.005 | 1.261 | 0.073 | Fig. S26 |
| RMV7$_{rare}$ *manLMN*::Janus | N-acetylglucosamine | 0.784 | 0.004 | 1.001 | 0.039 | Fig. S26 |
| RMV7$_{rare}$ *manLMN*::Janus | None | 0.833 | 0.006 | 1.155 | 0.058 | Fig. S26 |
| RMV7$_{rare}$ *manLMN*::Janus | Sialic acid | 0.647 | 0.005 | 1.172 | 0.073 | Fig. S26 |
| RMV7$_{wt}$ | Aminoglucose | 0.327 | 0.003 | 0.871 | 0.043 | Fig. S26 |
| RMV7$_{wt}$ | Galactose | 0.379 | 0.002 | 1.064 | 0.040 | Fig. S26 |
| RMV7$_{wt}$ | Glucose | 0.348 | 0.006 | 1.509 | 0.231 | Fig. S26 |
| RMV7$_{wt}$ | Mannose | 0.435 | 0.003 | 0.946 | 0.043 | Fig. S26 |
| RMV7$_{wt}$ | N-acetylglucosamine | 0.497 | 0.002 | 1.290 | 0.044 | Fig. S26 |
| RMV7$_{wt}$ | None | 0.370 | 0.005 | 1.083 | 0.102 | Fig. S26 |
| RMV7$_{wt}$ | Sialic acid | 0.324 | 0.001 | 1.023 | 0.032 | Fig. S26 |
| RMV7$_{wt}$ *manLMN*::Janus | Aminoglucose | 0.518 | 0.004 | 0.725 | 0.029 | Fig. S26 |
| RMV7$_{wt}$ *manLMN*::Janus | Galactose | 0.542 | 0.007 | 1.109 | 0.099 | Fig. S26 |
| RMV7$_{wt}$ *manLMN*::Janus | Glucose | 0.515 | 0.009 | 1.081 | 0.126 | Fig. S26 |
| RMV7$_{wt}$ *manLMN*::Janus | Mannose | 0.490 | 0.007 | 0.964 | 0.087 | Fig. S26 |

| | | | | | | |
|---|---|---|---|---|---|---|
| RMV7$_{wt}$ *manLMN*::Janus | N-acetylglucosamine | 0.535 | 0.008 | 0.855 | 0.066 | Fig. S26 |
| RMV7$_{wt}$ *manLMN*::Janus | None | 0.571 | 0.010 | 1.050 | 0.114 | Fig. S26 |
| RMV7$_{wt}$ *manLMN*::Janus | Sialic acid | 0.530 | 0.007 | 0.880 | 0.067 | Fig. S26 |
| RMV7$_{rare}$ | Aminoglucose | 0.734 | 0.001 | 1.206 | 0.009 | Fig. S26 |
| RMV7$_{rare}$ | Galactose | 0.719 | 0.002 | 1.292 | 0.028 | Fig. S26 |
| RMV7$_{rare}$ | Glucose | 0.718 | 0.006 | 1.624 | 0.115 | Fig. S26 |
| RMV7$_{rare}$ | Mannose | 0.616 | 0.006 | 1.352 | 0.104 | Fig. S26 |
| RMV7$_{rare}$ | N-acetylglucosamine | 0.756 | 0.004 | 1.472 | 0.064 | Fig. S26 |
| RMV7$_{rare}$ | | 0.740 | 0.003 | 1.511 | 0.053 | Fig. S36 |
| RMV7$_{rare}$ *nagA*::Janus | | 0.491 | 0.013 | 1.764 | 0.446 | Fig. S36 |
| RMV7$_{rare}$ *tfoX*::Janus | | 0.713 | 0.004 | 2.005 | 0.135 | Fig. S36 |
| RMV7$_{rare}$ *yjbK*::Janus | | 0.736 | 0.004 | 2.022 | 0.119 | Fig. S36 |
| RMV7$_{rare}$ *nagA*::Janus | Aminoglucose | 0.495 | 0.005 | 1.080 | 0.080 | Fig. S37 |
| RMV7$_{rare}$ *nagA*::Janus | Galactose | 0.490 | 0.005 | 1.231 | 0.089 | Fig. S37 |
| RMV7$_{rare}$ *nagA*::Janus | Glucose | 0.522 | 0.008 | 1.234 | 0.145 | Fig. S37 |
| RMV7$_{rare}$ *nagA*::Janus | Mannose | 0.351 | 0.023 | 1.240 | 0.031 | Fig. S37 |
| RMV7$_{rare}$ *nagA*::Janus | N-acetylglucosamine | 367.534 | $3.46x10^9$ | 0.040 | 0.135 | Fig. S37 |
| RMV7$_{rare}$ *nagA*::Janus | None | 0.401 | 0.010 | 1.611 | 0.396 | Fig. S37 |
| RMV7$_{rare}$ *nagA*::Janus | Sialic acid | 0.329 | 0.011 | 1.979 | 0.723 | Fig. S37 |
| RMV7$_{rare}$ *tfoX*::Janus | Aminoglucose | 0.753 | 0.003 | 1.525 | 0.052 | Fig. S37 |
| RMV7$_{rare}$ *tfoX*::Janus | Galactose | 0.727 | 0.002 | 1.239 | 0.028 | Fig. S37 |
| RMV7$_{rare}$ *tfoX*::Janus | Glucose | 0.738 | 0.031 | 0.418 | 0.043 | Fig. S37 |
| RMV7$_{rare}$ *tfoX*::Janus | Mannose | 0.658 | 0.006 | 1.178 | 0.035 | Fig. S37 |
| RMV7$_{rare}$ *tfoX*::Janus | N-acetylglucosamine | 0.737 | 0.002 | 1.586 | 0.046 | Fig. S37 |
| RMV7$_{rare}$ *tfoX*::Janus | None | 0.675 | 0.003 | 1.731 | 0.074 | Fig. S37 |
| RMV7$_{rare}$ *tfoX*::Janus | Sialic acid | 0.654 | 0.003 | 1.291 | 0.047 | Fig. S37 |
| RMV7$_{rare}$ *yjbK*::Janus | Aminoglucose | 0.803 | 0.002 | 1.469 | 0.035 | Fig. S37 |
| RMV7$_{rare}$ *yjbK*::Janus | Galactose | 0.725 | 0.002 | 1.320 | 0.021 | Fig. S37 |
| RMV7$_{rare}$ *yjbK*::Janus | Glucose | 0.673 | 0.014 | 0.544 | 0.045 | Fig. S37 |
| RMV7$_{rare}$ *yjbK*::Janus | Mannose | 0.653 | 0.005 | 1.424 | 0.067 | Fig. S37 |
| RMV7$_{rare}$ *yjbK*::Janus | N-acetylglucosamine | 0.738 | 0.002 | 1.763 | 0.053 | Fig. S37 |

| | | | | | | |
|---|---|---|---|---|---|---|
| RMV7$_{rare}$ *yjbK*::Janus | None | 0.659 | 0.002 | 1.675 | 0.044 | Fig. S37 |
| RMV7$_{rare}$ *yjbK*::Janus | Sialic acid | 0.662 | 0.003 | 1.349 | 0.045 | Fig. S37 |
| RMV7$_{rare}$ | | 0.687 | 0.003 | 1.549 | 0.060 | Fig. S40 |
| RMV7$_{rare}$ *rlrA*::Janus | | 0.649 | 0.004 | 1.775 | 0.098 | Fig. S40 |
| RMV7$_{rare}$ rrg*A*::Janus | | 0.691 | 0.004 | 1.698 | 0.080 | Fig. S40 |
| RMV7$_{rare}$ *rrgBC*::Janus | | 0.659 | 0.006 | 1.829 | 0.155 | Fig. S40 |
| RMV7$_{wt}$ | | 0.488 | 0.009 | 1.174 | 0.162 | Fig. S41 |
| RMV7$_{wt}$ PRCI$_{dnaN}$::Janus | | 0.559 | 0.011 | 1.123 | 0.172 | Fig. S41 |
| RMV7$_{rare}$ | 35 °C | 0.676 | 0.004 | 1.688 | 0.097 | Fig. S43 |
| RMV7$_{rare}$ | 40 °C | 0.266 | 0.023 | 9.594 | 67.500 | Fig. S43 |
| RMV7$_{rare}$ *clpP*::*cat* | 35 °C | 0.594 | 0.008 | 1.744 | 0.228 | Fig. S43 |
| RMV7$_{rare}$ *clpP*::*cat* | 40 °C | 0.124 | 0.079 | 0.086 | 0.217 | Fig. S43 |
| RMV7$_{rare}$ *clpP*::*cat* *hrcA*::Janus | 35 °C | 0.485 | 0.013 | 2.482 | 0.764 | Fig. S43 |
| RMV7$_{rare}$ *clpP*::*cat* *hrcA*::Janus | 40 °C | 0.017 | 0.115 | 0.013 | 0.100 | Fig. S43 |
| RMV7$_{rare}$ *hrcA*::Janus | 35 °C | 0.501 | 0.011 | 2.719 | 0.692 | Fig. S43 |
| RMV7$_{rare}$ *hrcA*::Janus | 40 °C | 0.031 | 0.143 | 0.021 | 0.126 | Fig. S43 |
| RMV7$_{wt}$ | 35 °C | 0.429 | 0.008 | 1.315 | 0.203 | Fig. S43 |
| RMV7$_{wt}$ | 40 °C | 0.009 | 0.051 | 0.015 | 0.095 | Fig. S43 |
| RMV7$_{wt}$ *clpP*::*cat* | 35 °C | 0.353 | 0.004 | 0.929 | 0.070 | Fig. S43 |
| RMV7$_{wt}$ *clpP*::*cat* | 40 °C | 0.004 | 0.051 | 0.009 | 0.123 | Fig. S43 |
| RMV7$_{wt}$ *clpP*::*cat* *hrcA*::Janus | 35 °C | 0.212 | 0.016 | 2.160 | 1.801 | Fig. S43 |
| RMV7$_{wt}$ *clpP*::*cat* *hrcA*::Janus | 40 °C | 0.017 | 0.001 | 0.216 | 0.033 | Fig. S43 |
| RMV7$_{wt}$ *hrcA*::Janus | 35 °C | 0.450 | 0.003 | 1.610 | 0.107 | Fig. S43 |
| RMV7$_{wt}$ *hrcA*::Janus | 40 °C | 0.042 | 0.001 | 0.273 | 0.012 | Fig. S43 |

**Table S5** Genotypes constructed and analysed in this study.

| Isolate name | List and description of genotypes | Reference or source (this study unless specified) |
|---|---|---|
| *S. pneumoniae* R6 | R6 *rpsL*Δivr*<br>R6 *rpsL*Δivr rpoB**<br>R6 *rpsL* Δivr tfoX*::Janus<br>R6 *rpsL* Δivr manLMN*::Janus<br>R6 *rpsL* Δivr yjbK*::Janus<br>R6 *rpsL* Δivr hrcA*::Janus<br>R6 *rpsL* Δivr hrcA* restored | Apagyi *et al* 2018 |
| *S. pneumoniae* RMV7 | $RMV7_{wt}$ (RMV7 *rpsL**)<br>$RMV7_{domi}$ (RMV7 *rpsL** $tvr_{TRDIII-iii}$ *tvrR*::Janus)<br>$RMV7_{rare}$ (RMV7 *rpsL** $tvr_{TRDIII-i}$ Δ*tvrR*)<br>$RMV7_{wt}$ *rpsL* tvr*::*cat*<br>RMV7 $tvr_{domi}$::Janus (RMV7 *rpsL* tvr*::$tvr_{TRDIII-iii}$ *tvrR*::Janus)<br>RMV7 $tvr_{rare}$::Janus (RMV7 *rpsL* tvr*::$tvr_{TRDIII-i}$ *tvrR*::Janus) | Kwun *et al* 2018<br>Kwun *et al* 2018<br>Kwun *et al* 2018 |
| $RMV7_{wt}$ | $RMV7_{wt}$ *phoB*::Janus<br>$RMV7_{wt}$ *ciaRH*::Janus<br>$RMV7_{wt}$ *htrA*::Janus<br>$RMV7_{wt}$ *manLMN*::Janus<br>$RMV7_{wt}$ *tfoX*::Janus<br>$RMV7_{wt}$ *yjbK*::Janus<br>$RMV7_{wt}$ *nagA*::Janus<br>$RMV7_{wt}$ *rlrA*::Janus<br>$RMV7_{wt}$ *rlrArrgABC*::Janus<br>$RMV7_{wt}$ *rrgA*::Janus<br>$RMV7_{wt}$ *rrgBC*::Janus<br>$RMV7_{wt}$ $PRCI_{dnaN}$::Janus<br>$RMV7_{wt}$ $PRCI_{dnaN+att}$::Janus | |

| | |
|---|---|
| | RMV7$_{wt}$ PRCI$_{reg\text{-}IONPJBJN00496}$::Janus<br>RMV7$_{wt}$ PRCI$_{rep}$::Janus<br>RMV7$_{wt}$ PRCI$_{int\text{-}rep}$::Janus<br>RMV7$_{wt}$ PRCI$_{reg}$::Janus<br>RMV7$_{wt}$ PRCI$_{dnaN+att}$::Janus *hrcA*::*cat*<br>RMV7$_{wt}$ PRCI$_{dnaN+att}$::Janus *clpP*::*cat*<br>RMV7$_{wt}$ *clpP*::*cat*<br>RMV7$_{wt}$ *clpP*::*cat* *hrcA*::Janus<br>RMV7$_{wt}$ *hrcA*::*cat*<br>RMV7$_{wt}$ *hrcA*::Janus |
| RMV7$_{rare}$ | RMV7$_{rare}$ *phoB*::Janus<br>RMV7$_{rare}$ *phoB*::*phoB*$_{wt}$<br>RMV7$_{rare}$ *phoB* restored<br>RMV7$_{rare}$ *piuA*::Janus<br>RMV7$_{rare}$ *ciaRH*::Janus<br>RMV7$_{rare}$ *htrA*::Janus<br>RMV7$_{rare}$ *manLMN*::Janus<br>RMV7$_{rare}$ *manL*::Janus<br>RMV7$_{rare}$ *manL* restored<br>RMV7$_{rare}$ *tfoX*::Janus<br>RMV7$_{rare}$ *tfoX* restored<br>RMV7$_{rare}$ *yjbK*::Janus<br>RMV7$_{rare}$ *yjbK* restored<br>RMV7$_{rare}$ *nagA*::Janus<br>RMV7$_{rare}$ *manLMN*::Janus *tfoX*::*cat*<br>RMV7$_{rare}$ *manLMN*::Janus *yjbK*::*cat*<br>RMV7$_{rare}$ *yjbK*::Janus *tfoX*::*cat*<br>RMV7$_{rare}$ *rlrA*::Janus<br>RMV7$_{rare}$ *rlrA* restored<br>RMV7$_{rare}$ *rlrArrgABC*::Janus<br>RMV7$_{rare}$ *rrgA*::Janus |

| | | |
|---|---|---|
| | RMV7$_{rare}$ *rrgBC*::Janus<br>RMV7$_{rare}$ *hrcA*::Janus<br>RMV7$_{rare}$ *hrcA* restored<br>RMV7$_{rare}$ *clpP*::*cat*<br>RMV7$_{rare}$ *clpP*::*cat hrcA::*Janus<br>RMV7$_{rare}$ *hrcA*::Janus *manLMN*::*cat* | |
| *S. pneumoniae* RMV5 | RMV7 *rpsL* tvr*$_{TRDIV-iii}$ $\Delta$*tvrR*<br>RMV7 *rpsL* tvr*$_{TRDI-i}$ $\Delta$*tvrR* | Kwun *et al* 2018<br>Kwun *et al* 2018 |
| *S. pneumoniae* RMV6 | RMV7 *rpsL* tvr*$_{TRDII-iii}$ $\Delta$*tvrR*<br>RMV7 *rpsL* tvr*$_{TRDIII-i}$ $\Delta$*tvrR* | Kwun *et al* 2018<br>Kwun *et al* 2018 |
| *S. pneumoniae* RMV8 | RMV7 *rpsL* tvr*$_{TRDI-i}$ $\Delta$*tvrR*<br>RMV7 *rpsL* tvr*$_{TRDIV-ii}$ $\Delta$*tvrR* | Kwun *et al* 2018<br>Kwun *et al* 2018 |
| *E. coli* | DH5α | Sigma |

**Table S6** Oligonucleotides used in this study

| Name of oligonucleotide | Sequence | Function |
|---|---|---|
| hsdML | GCGGATGGTTTAAGTTTGGA | Checking arrangements of *tvr* locus |
| Trd III R v1 | TTCAATATAAAACTGCCATC | Checking arrangements of *tvr* locus |
| Trd IV R v2 | ATTTATCTCTGTTGAAGTTTTAGTCAG | Checking arrangements of *tvr* locus |
| Trd i R v1 | CTCCTCACTAAACAACTCATCTGA | Checking arrangements of *tvr* locus |
| Trd iii R v2 | TCCAAATGCTGTATCTACATTCCA | Checking arrangements of *tvr* locus |
| qRT rpoA For | TGGTCGTGGATATGTACCTGC | Transcriptional analysis |
| qRT rpoA Rev | CACGAGCAGGTTCCACTTGA | Transcriptional analysis |
| rpoB Full v2 For | GGCAGGACATGACGTTCAAT | Amplification of *rpoB** gene (conferring resistance to rifampicin) |
| rpoB Full v2 Rev | GCTTCCGCTTTTGTTGCTTT | Amplification of *rpoB** gene (conferring resistance to rifampicin) |
| qRT 507 For | AGGGAGAAAACTCTAATACCGT | Transcriptional analysis |
| qRT 507 For | GGCTTTTCTCAAGGTTTCAGGT | Transcriptional analysis |
| qRT hrcA For | TACCAAAACGCACGAACCTG | Transcriptional analysis |
| qRT hrcA Rev | AAAACCAGCAACACTTGGCA | Transcriptional analysis |
| qRT dnaK For | GGCTACGCTGAAGACTACCT | Transcriptional analysis |
| qRT dnaK Rev | CTGCTGCAGTTGGTTCGTTA | Transcriptional analysis |
| qRT manL For | GCGTGCCAATCAAGACTCTT | Transcriptional analysis |
| qRT manL Rev | ATTCAACACCCAAGTCACGC | Transcriptional analysis |
| qRT ciaR For | TGAACTGGGAGCGGATGATT | Transcriptional analysis |
| qRT ciaR Rev | ATCGAACTCTTTCCCCAGCA | Transcriptional analysis |
| qRT comYC For | TTGGTGGAGATGTTGGTGGT | Transcriptional analysis |
| qRT comYC Rev | TCGCCCATCTGCTTGTAACT | Transcriptional analysis |
| qRT comX For | TGGGAATTGTCGGATTGGGA | Transcriptional analysis |
| qRT comX Rev | CGCTTCTGACTTTCCTGCTT | Transcriptional analysis |
| qRT comEA For | AGGCTCTGGTTTACGTTCCT | Transcriptional analysis |
| qRT comEA Rev | TGTCCTGAGCTCGTTTTCCT | Transcriptional analysis |

| | | |
|---|---|---|
| qRT comD For | AAGCAGATGGAAGTAGCAGTT | Transcriptional analysis |
| qRT comD Rev | ATCCGACTCCGCGATTTCTT | Transcriptional analysis |
| qRT htrA For | GCGGCCCACTGATCAATATT | Transcriptional analysis |
| qRT htrA rev | ACGCGTCACTTTTCCGTTTT | Transcriptional analysis |
| qRT groEL For | CGTGGACAAAGATAGCACGG | Transcriptional analysis |
| qRT groEL Rev | CCACCTGACAATTTGGCCAA | Transcriptional analysis |
| qRT clpP For | TTATGCTGACAGGTCCGGTT | Transcriptional analysis |
| qRT clpP Rev | CCCACTTGATGCGATGACAG | Transcriptional analysis |
| qRT clpE For | GCGCGACGATGAGATTATCC | Transcriptional analysis |
| qRT clpE Rev | TCGAATCCCCGTTCCTTGAA | Transcriptional analysis |
| phoB Up For | TAAGATTGGTGTCAAAGCAGGCCAGT | Removing *phoB* |
| phoB Up Rev ApaI | AAGGGCCCTTCTCTAATCATTTCCTCGTC | Removing *phoB* |
| phoB Down For BamHI | AAGGATCCGCTTTAGATTGTATCGTGAC | Removing *phoB* |
| phoB Down Rev | TATTCTCTAGGATAATTCTAAGACTGG | Removing *phoB* |
| phoB internal For | ATGATAGGCTGGTAGGAAATTTTC | Checking the sequence of *phoB* |
| htrA Up For | TCGAACCTGCGACCGTTCGCTTA | Removing *htrA* |
| htrA Up Rev ApaI | AAGGGCCCATGTTTCATATTTGCCTCCAT | Removing *htrA* |
| htrA Down For BamHI | AAGGATCCGAATCTTAATTGACATCTATG | Removing *htrA* |
| htrA Down Rev | AATGTTACCAAACTTTATCCACAGGTT | Removing *htrA* |
| hrcA Up For | GGCGTAAATTTTGCTATAATATTTTCG | Removing *hrcA* |
| hrcA Up Rev ApaI | AAGGGCCCGCTAGAGTTAATAGACTCTTGC | Removing *hrcA* |
| hrcA Down For BamHI | AAGGATCCAATCAAGTCAATGTGGTCAAC | Removing *hrcA* |
| hrcA Down Rev | TGGTTCGCTTCGCTCACTATTACAAA | Removing *hrcA* |
| yjbK Up For | CACGATAATCTGTATAGAAGCCCAAATGT | Removing *yjbK* |
| yjbK Up Rev ApaI | AAGGGCCCCAATAGTGTTTTCAATTCAAT | Removing *yjbK* |
| yjbK Down For BamHI | AAGGATCCAGTATGAAAAATAGCTGAAAT | Removing *yjbK* |
| yjbK Down Rev | ATATTAGTATTGTCAAGAAGCTCAGCAGC | Removing *yjbK* |
| ciaRH Up For | CGTCATAACTATGACCGTCTTGGTTTTG | Removing *ciaRH* |
| ciaRH Up Rev ApaI | AAGGGCCCCAGACCTAGGTCATCCTCAA | Removing *ciaRH* |
| ciaRH Down For BamHI | AAGGATCCAATCTTTGAAGTGAAGATTGC | Removing *ciaRH* |
| ciaRH Down Rev | CCAGAAGTTAATCTGACGACTGCTAGT | Removing *ciaRH* |

| | | |
|---|---|---|
| tvr Up For | ACAACCTTTGACTACGCCTATACACC | Removing *tvr* locus |
| tvr Up Rev ApaI | AAGGGCCCATCTTCCCTTTTCTTTAGTT | Removing *tvr* locus |
| tvr Down For BamHI | AAGATATCATTCTAAAGTGATTGCCATGC | Removing *tvr* locus |
| tvr Down Rev | AAGCCCTTCTTGCTAAGAACGACATTC | Removing *tvr* locus |
| tfoX Up For | GGCGCAAAATACGACCATCATG | Removing *tfoX* |
| tfoX Up Rev ApaI | AAGGGCCCGTTCTTACTTGATGCCAT | Removing *tfoX* |
| tfoX Down For BamHI | AAGGATCCGAAATGATTTGAGTGGAAG | Removing *tfoX* |
| tfoX Down Rev | TGCAAAAGCACCGGAATTGGTGA | Removing *tfoX* |
| manLMN Up For | GCCAAATCCAAGGACGTATGGTACTCGA | Removing *manLMN* |
| manLMN Up Rev ApaI | AAGGGCCCCATCACGCGACTAGCTTGGTT | Removing *manLMN* |
| manLMN Down For BamHI | AAGGATCCAACTTGATTCGTATGTCATTC | Removing *manLMN* |
| manLMN Down Rev | AGGAATCAGATTTTAGAACCTGGTTCC | Removing *manLMN* |
| manL Down For BamHI | AAGGATCCAACAAAGCCAATGTCAAATAA | Removing *manL* |
| manL Down Rev | TCCAAGACCCTTGTAAGAAGGTTG | Removing *manL* |
| piuA Up For | TAAGGATAAGAGGTCCCACTTAAAAC | Removing *piuA* |
| piuA Up Rev ApaI | AAGGGCCCAGAATTAGAAGAACAAGCGCT | Removing *piuA* |
| piuA Down For BamHI | AAGGATCCGTAATTCCAGATAATACACCG | Removing *piuA* |
| piuA Down Rev | CTTGCAATTATTGCAAGTGATCTTATC | Removing *piuA* |
| nagA Up For | ACAGACTTAACTGTCAATGGCTTGAAG | Removing *nagA* |
| nagA Up Rev ApaI | AAGGGCCCCACAAGTTCCAAGTAACCA | Removing *nagA* |
| nagA Down For BamHI | AAGGATCCGCTAAATCCGTTCACATCGA | Removing *nagA* |
| nagA Down Rev | CGTCTAGGTCTACCATAATCAATTTAGC | Removing *nagA* |
| rlrA Up For | CATTCTTCTCAACTTCAACAGTCCATT | Removing *rlrA* |
| rlrA Up Rev ApaI | AAGGGCCCCAGTTCATCTAGTTCAATAG | Removing *rlrA* |
| rlrA Down For BamHI | AAGGATCCTTAGATTTGCATGTTAGCCA | Removing *rlrA* |
| rlrA Down Rev | CTGAGCAATGAAAGCCAATTTCCC | Removing *rlrA* |
| rrgA Up For | GAAATATACTCTTTTTGTGTAAATTC | Removing *rrgA* |
| rrgA Up Rev ApaI | AAGGGCCCTTCAGGCGTTTCTGCTAAA | Removing *rrgA* |
| rrgA Down For BamHI | AAGGATCCACAGGTGAAGATGGTAAG | Removing *rrgA* |
| rrgA Down Rev | TTTTCATCAATAATTTCATTATTAG | Removing *rrgA* |
| rrgBC Up For | CCTAAGGAAGTAGAAAAGAACACAGTG | Removing *rrgBC* |

| | | |
|---|---|---|
| rrgBC Up Rev ApaI | AAGGGCCCGGCAGCAAGCATTGTTAAAAA | Removing *rrgBC* |
| rrgBC Down For BamHI | AAGGATCCTTGATGCTTGTTGCCATTTTG | Removing *rrgBC* |
| rrgBC Down Rev | GGTCAAATCCGTAAACATCTTAGCCGT | Removing *rrgBC* |
| clpP Up For | AATTGAAGTTATTAATCACCCACTGA | Removing *clpP* |
| clpP Up Rev ApaI | AAGGGCCCAGAACGTTCTCCACGGCTTGT | Removing *clpP* |
| clpP Down For BamHI | AAGGATCCAGCGCCCAGGAAACACTTGAA | Removing *clpP* |
| clpP Down Rev | AATGGAACACCTGCTTTTGTAGCGTTC | Removing *clpP* |
| PRCI Up For v2 | TCGTAATTCCTAGCCGTTCTCTACGCG | Removing PRCI with *att* sites |
| PRCI Up Rev ApaI v2 | AAGGGCCCTCAGTCAATAATTTTATTCA | Removing PRCI with *att* sites |
| PRCI Down For BamHI v2 | AAGGATCCGTGTTATAATATTAGGGATTG | Removing PRCI with *att* sites |
| PRCI Down Rev v2 | TTCTTCCTCAGCACGCGCAGAAATAAC | Removing PRCI with *att* sites |
| PRCI Up For | CCAACGATACCTGCTGTCA | Removing PRCI |
| PRCI Up Rev ApaI | AAGGGCCCAGCAGGGTAAATACGGCAGA | Removing PRCI |
| PRCI Down For BamHI | AAGGATCCCTTTAACCCTGCCCGTGATG | Removing PRCI |
| PRCI Down Rev | AAAAGGCTAATCGTTGGGAAATT | Removing PRCI |
| PRCI Regulator Up For | GTATACAACCGTCAACGATTGGGTAA | Removing PRCI’s regulator |
| PRCI Regulator Up Rev ApaI | AAGGGCCCGTGTATTCTATATGTACGATG | Removing PRCI’s regulator |
| PRCI Regulator For BamHI | AAGGATCCCGAATAACATCTAACGATTTT | Removing PRCI’s regulator |
| PRCI Regulator Down Rev | CGCCCTTGAGTCTTTTGGGATGATGAT | Removing PRCI’s regulator |
| dinD Up For | CCCTATTCCGATGAAAATACTGTTAAC | Removing IONPJBJN_00496 |
| dinD Up Rev ApaI | AAGGGCCCAATAAAGTATCCTTTCTAAAA | Removing IONPJBJN_00496 |
| tvrR Inf Up For | AATCACCATTACGATTCCAAGTGAATTT | Removing N-terminal 267bp of *tvrR* |
| tvrR Inf Up Rev ApaI | GGGGGCCCAATTTCTCACTTTCTTATTCA | Removing N-terminal 267bp of *tvrR* |
| tvrR Inf Up R BamHI | TTGGATCCAATTTCTCACTTTCTTATTCA | Removing Janus from *tvrR*::Janus mutant |
| tvrR Inf Down For BamHI | TTGGATCCTTAAGGAGTTATTTAGCAAATT | Removing N-terminal 267bp of *tvrR* |
| tvrR Inf Down Rev | CTTTTAAGAGATGAATTTTTGGTGTGA | Removing N-terminal 267bp of *tvrR* |
| tvrR Up For | AATCACCATTACGATTCCAAGTGAATTT | Removing C-terminal 681bp of *tvrR* |
| tvrR Up Rev ApaI | TTGGGCCCACTACAATCCTTTTCAGACTG | Removing C-terminal 681bp of *tvrR* |
| tvrR Down For BamHI | TTGGATCCAAGCAAATCCCGATATTCCGA | Removing C-terminal 681bp of *tvrR* |
| Janus For ApaI | TTGGGCCCCCGTTTGATTTTTAATGGATAATGTG | Amplifying Janus |

| | | |
|---|---|---|
| Janus Rev BamHI | ATGGATCCCCTTTCCTTATGCTTTTGGACG | Amplifying Janus |
| Cat For ApaI | AAGGGCCCAGTGGGATATTTTTAAAATAT | Amplifying *cat* |
| Cat Rev BamHI | AAGGATCCTTATAAAAGCCAGTCATTAGG | Amplifying *cat* |

**Table S7** Correspondence between figures and files available from https://figshare.com/projects/Diverse_regulatory_pathways_modulate_bet_hedging_of_competence_induction_in_epigenetically-differentiated_phase_variants_of_Streptococcus_pneumoniae/171060.

| Figure | Dataset | File |
|---|---|---|
| 1A | Experimental data from the analysis of RMV7 phase variants | Comparison_of_RMV_phase_variants.csv |
| 1C | Experimental data from the analysis of RMV7 phase variants | RMV7_phase_variant_frequencies.csv |
| 1D | Experimental data from the analysis of RMV7 phase variants | RMV7_domi_rare_wt_transformation_frequencies.csv |
| 1E | Experimental data from the analysis of RMV7 phase variants | RMV7_domi_rare_wt_biofilm_thicknesses.csv |
| 1F | Experimental data from the analysis of RMV7 phase variants | RMV7_tvr_domi_rare_passage_transformation_frequencies.csv |
| 2 | RNA-seq data from a comparison of RMV7 epigenetic phase variants | RMV7_RNA_seq_statistics.csv |
| 3A | Experimental data from the analysis of RMV7 phase variants | RMV7_rare_carbohydrates_transformation_efficiency.csv |
| 3B | Experimental data from the analysis of RMV7 phase variants | RMV7_wt_carbohydrates_transformation_efficiency.csv |
| 3C | Experimental data from the analysis of RMV7 phase variants | RMV7_rare_carbohydrates_biofilm.csv |
| 3D | Experimental data from the analysis of RMV7 phase variants | RMV7_rare_GlcNAc_double_mutants_transformation_efficiencies.csv |
| 3E | Experimental data from the analysis of RMV7 phase variants | Comparison_of_RMV_phase_variants_in_GlcNAc.csv |
| 4A | Experimental data from the analysis of RMV7 phase variants | RMV7_wt_PRCI_mutants.csv |
| 4B | Experimental data from the analysis of RMV7 phase variants | RMV7_wt_PRCI_chaperone_mutants.csv |
| 4C | Experimental data from the analysis of RMV7 phase variants | RMV7_wt_PRCI_mutant_gene_expression.csv |

| | | |
|---|---|---|
| 5A | Experimental data from the analysis of RMV7 phase variants | RMV7_heat_shock_transformation.csv |
| 5B | Experimental data from the analysis of RMV7 phase variants | RMV7_rare_CaCl2_titration.csv |
| 5C | Experimental data from the analysis of RMV7 phase variants | RMV7_rare_CaCl2_effect_on_hrcA.csv |
| 5D | Experimental data from the analysis of RMV7 phase variants | RMV7_rare_comEAX_expression.csv |
| 5E | Experimental data from the analysis of RMV7 phase variants | RMV7_rare_manLMN_hrcA_double_mutants.csv |
| S1 | Agarose gel electrophoresis separation of PCR amplicons from RMV7 tvr loci | RMV7_wt_tvr_variation.jpg |
| S1 | Agarose gel electrophoresis separation of PCR amplicons from RMV7 tvr loci | RMV7_domi_rare_tvr_domi_tvr_rare_tvr_arrangements.jpg |
| S2A | Experimental data from the analysis of RMV7 phase variants | RMV7_wt_spontaneous_resistance_frequency.csv |
| S2B | Experimental data from the analysis of RMV7 phase variants | RMV7_overnight_transformation_efficiencies.csv |
| S3 | Morphology of RMV7 variant colonies | RMV7_domi_x10_magnification.jpg |
| S3 | Morphology of RMV7 variant colonies | RMV7_domi_x4_magnification.jpg |
| S3 | Morphology of RMV7 variant colonies | RMV7_rare_x10_magnification.jpg |
| S3 | Morphology of RMV7 variant colonies | RMV7_rare_x4_magnification.jpg |
| S3 | Morphology of RMV7 variant colonies | RMV7_wt_x10_magnification.jpg |
| S3 | Morphology of RMV7 variant colonies | RMV7_wt_x4_magnification.jpg |
| S4A | Experimental data from the analysis of RMV7 phase variants | RMV7_phoB_transformation_efficiency.csv |
| S4B | Experimental data from the analysis of RMV7 phase variants | RMV7_pstS_transformation_efficiency.csv |
| S5B | Agarose gel electrophoresis separation of PCR amplicons from RMV7 tvr loci | RMV7_domi_rare_tvr_domi_tvr_rare_tvr_arrangements.jpg |
| S5C | Experimental data from the analysis of RMV7 phase variants | RMV7_tvr_mutant_transformation_efficiencies.csv |
| S5D | Experimental data from the analysis of RMV7 phase variants | RMV7_tvr_mutant_biofilm_thicknesses.csv |

| | | |
|---|---|---|
| S6 | RNA-seq data from a comparison of RMV7 epigenetic phase variants | RMV7_RNA_seq_fragment_lengths.csv |
| S7 | RNA-seq data from a comparison of RMV7 epigenetic phase variants | RMV7_RNA_seq_expression.csv |
| S8 | RNA-seq data from a comparison of RMV7 epigenetic phase variants | RMV7_RNA_seq_statistics.csv |
| S9 | RNA-seq data from a comparison of RMV7 epigenetic phase variants | RMV7_RNA_seq_statistics.csv |
| S10 | RNA-seq data from a comparison of RMV7 epigenetic phase variants | RMV7_RNA_seq_JSDs.csv |
| S11 | RNA-seq data from a comparison of RMV7 epigenetic phase variants | RMV7_RNA_seq_expression.csv |
| S12 | RNA-seq data from a comparison of RMV7 epigenetic phase variants | RMV7_RNA_seq_expression.csv |
| S13 | RNA-seq data from a comparison of RMV7 epigenetic phase variants | RMV7_RNA_seq_expression.csv |
| S14 | Experimental data from the analysis of RMV7 phase variants | RMV7_qRTPCR.csv |
| S15 | RNA-seq data from a comparison of RMV7 epigenetic phase variants | RMV7_RNA_seq_expression.csv |
| S16 | Experimental data from the analysis of RMV7 phase variants | RMV7_motif_separation_expression_change_comparison.csv |
| S18 | Experimental data from the analysis of RMV7 phase variants | RMV7_piuA_transformation_efficiency.csv |
| S19 | Experimental data from the analysis of RMV7 phase variants | RMV7_motif_distribution.csv |
| S20 | RNA-seq data from a comparison of RMV7 epigenetic phase variants | RMV7_RNA_seq_expression.csv |
| S21 | RNA-seq data from a comparison of RMV7 epigenetic phase variants | RMV7_RNA_seq_expression.csv |
| S22 | RNA-seq data from a comparison of RMV7 epigenetic phase variants | RMV7_RNA_seq_expression.csv |
| S23 | RNA-seq data from a comparison of RMV7 epigenetic phase variants | RMV7_RNA_seq_expression.csv |

| | | |
|---|---|---|
| S24 | Experimental data from the analysis of RMV7 phase variants | RMV7_ciaRH_htrA_transformation_efficiency.csv |
| S25 | Experimental data from the analysis of RMV7 phase variants | RMV7_rare_carbohydrates_transformation_efficiency.csv |
| S26 | Experimental data from the analysis of RMV7 phase variants | RMV7_manLMN_growth_curves.csv |
| S27 | Experimental data from the analysis of RMV7 phase variants | RMV7_manLMN_growth_curve_statistics.csv |
| S28 | Experimental data from the analysis of RMV7 phase variants | RMV7_rare_manL_carbohydrate_transformation_efficiency.csv |
| S34 | Experimental data from the analysis of RMV7 phase variants | RMV7_cAMP_concentrations.csv |
| S35 | Experimental data from the analysis of RMV7 phase variants | RMV7_cAMP_transformation_efficiency.csv |
| S36 | Experimental data from the analysis of RMV7 phase variants | RMV7_rare_nagA_tfoX_yjbK_growth_curves.csv |
| S37A | Experimental data from the analysis of RMV7 phase variants | RMV7_rare_nagA_tfoX_yjbK_carbohydrate_growth_curves.csv |
| S37B | Experimental data from the analysis of RMV7 phase variants | RMV7_nagA_transformation_efficiency.csv |
| S37C | Experimental data from the analysis of RMV7 phase variants | RMV7_manL_nagA_expression_levels.csv |
| S38 | Experimental data from the analysis of RMV7 phase variants | R6_manLMN_tfoX_yjbK_transformation_efficiency.csv |
| S39A | Experimental data from the analysis of RMV7 phase variants | RMV7_pilus_growth_curves.csv |
| S39B | Experimental data from the analysis of RMV7 phase variants | RMV7_pilus_biofilm_thicknesses.csv |
| S40 | Experimental data from the analysis of RMV7 phase variants | RMV7_PRCI_growth_curves.csv |
| S41A | Experimental data from the analysis of RMV7 phase variants | RMV7_clpPE_expression.csv |
| S41B | Experimental data from the analysis of RMV7 phase variants | RMV7_rare_clpP_expression_with_CaCl2.csv |

| | | |
|---|---|---|
| S42 | Experimental data from the analysis of RMV7 phase variants | RMV7_heat_shock_growth_curves.csv |
| S43 | Experimental data from the analysis of RMV7 phase variants | RMV7_chaperone_expression_with_CaCl2.csv |
| S44 | Experimental data from the analysis of RMV7 phase variants | R6_CaCl2_titration.csv |
| S45 | Experimental data from the analysis of RMV7 phase variants | RMV7_qRTPCR_CaCl2_expression_effect.csv |
| S46 | Experimental data from the analysis of RMV7 phase variants | RMV7_heat_shock_hrcA_restoration.csv |
| S47 | Experimental data from the analysis of RMV7 phase variants | RMV7_recombination_heterogeneity.csv |
| S48 | Maximum likelihood phylogeny of TfoX N terminal domain | TfoX_N_terminal_domain.tre |
| S49 | RNA-seq data from a comparison of RMV7 epigenetic phase variants | RMV7_RNA_seq_expression.csv |
